# The spectrum of covariance matrices of randomly connected recurrent neuronal networks

**DOI:** 10.1101/2020.08.31.274936

**Authors:** Yu Hu, Haim Sompolinsky

## Abstract

A key question in theoretical neuroscience is the relation between the connectivity structure and the collective dynamics of a network of neurons. Here we study the connectivity-dynamics relation as reflected in the distribution of eigenvalues of the covariance matrix of the dynamic fluctuations of the neuronal activities, which is closely related to the network’s Principal Component Analysis (PCA) and the associated effective dimensionality. We consider the spontaneous fluctuations around a steady state in a randomly connected recurrent network of stochastic neurons. An exact analytical expression for the covariance eigenvalue distribution in the large-network limit can be obtained using results from random matrices. The distribution has a finitely supported smooth bulk spectrum and exhibits an approximate power-law tail for coupling matrices near the critical edge. We generalize the results to include connectivity motifs and discuss extensions to Excitatory-Inhibitory networks. The theoretical results are compared with those from finite-size networks and and the effects of temporal and spatial sampling are studied. Preliminary application to whole-brain imaging data is presented. Using simple connectivity models, our work provides theoretical predictions for the covariance spectrum, a fundamental property of recurrent neuronal dynamics, that can be compared with experimental data.

## 1 Introduction

Collective dynamics in networked systems are of great interest, with numerous applications in many fields, including neuroscience, spin glasses, social and ecological networks [57]. Many studies of neuronal networks have focused on how certain statistics of dynamics depend on the network’s connectivity structure [48, 18, 55], including the population average [27] and variance [11] of pairwise correlations. Although powerful and directly comparable with experimental data, these are *local* features of dynamics, and can therefore be estimated just from the local measurements of the activity of involved neurons. However, important salient aspects of the dynamics are captured only at the *global* scale. Probing these aspects experimentally requires simultaneously recorded activities of a population of neurons, which has recently become increasingly feasible.

An important example of a global aspect of population dynamics is the eigenvalues of the covariance matrix, which are complicated nonlinear functions of all matrix elements. These eigenvalues arise naturally when performing the widely used Principal Component Analysis (PCA) of population activity, where they correspond to the amount of variance contained in each principal component of the activity. Another example that has received substantial recent interest [34, 31, 46, 50, 43] is the effective dimensionality of neural population activity, which can be defined based on the moments of the covariance eigenvalues. Many recent experimental studies have observed a low dimensional dynamics of neurons in the brain [45, 15], and theoretical investigations have illustrated the importance of having a low dimensionality for brain function and computation [14], such as when representing stimuli [12] and generating motor outputs [15].

As the experimental techniques of measuring the activity of large population of neurons in biological networks become increasingly available, new opportunities arise for studying how the network’s connectivity structure affects these global aspects of population dynamics.

In this work, we study the eigenvalue distribution (i.e., spectrum) of the covariance matrix of spontaneous activity in a large recurrent network of stochastic neurons with random connectivity. We focus on several basic and widely used models of random connectivity, including independent and identical Gaussian distributed connectivity [48] (Section 3.1), networks with connectivity motifs [49, 38, 58, 27, 26] (Section 3.3), and random Excitation-Inhibition (EI) networks (Section 3.5). Random connectivity has been a fundamental model in theoretical studies of neuronal network dynamics[31, 48, 28]. It can be motivated as a minimal model to capture the highly complex, disordered connections observed in many neuronal circuits, such as in the cortex. Some aspects of these covariance spectra might be distinct from those under ordered, deterministic connectivity (Section 4.1).

The dynamics considered here is simple where the activity fluctuations around the steady-state are described by a linear response [30, 52]. Despite the simple dynamics and minimal connectivity model, we find the resulting spectrum has a continuous bulk of nontrivial shape exhibiting interesting features such as a power-law long tail of large eigenvalues (Section 3.2), and strong effects due to the non-normality of the connectivity matrix (Section S5.2). These covariance spectra highlight interesting population-level structures of neuronal co-fluctuations shaped by recurrent interactions that were previously unexplored.

Using the theory of the covariance spectrum, we derive closed-form expressions for the effective dimensionality (previously known for the simple random iid Gaussian connectivity [11]) We show that the continuous bulk spectrum has the advantage over low-order statistics such as the dimensionality thanks to its robustness to low rank perturbations (Sections 3.3 to 3.5 and 3.8).

Our analytically derived eigenvalue distributions can be readily compared to real activity data of recurrent neural circuits or simulations of more sophisticated computational models. We provide ready-to-use code to facilitate such applications (Section 5.7). An example of such an application for a whole-brain calcium imaging data is presented in Section 3.8.

## 2 Model

### 2.1 Neuronal networks with random recurrent connectivity

We consider a recurrent network of linear rate neurons driven by noise

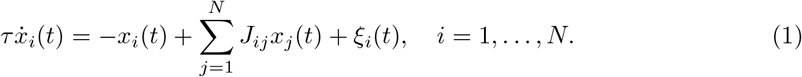

Here *x_i_*(*t*) is the firing rate of neuron *i*. *J_ij_* describes the recurrent interaction from neuron *j* to *i*. *τ* is a time constant describing how quickly the firing rates changes in response to inputs. The network is driven by independent Gaussian white noise *ξ_i_*(*t*) with variance *σ*^2^, that is, the expectation 〈*ξ_i_*(*t*)*ξ_j_*(*t* + *τ*)〉 = *σ*^2^*δ_ij_δ*(*τ*).

We focus on the structure of long time scale co-fluctuations in the network, which are described by the *long time window* covariance 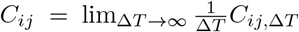. *C*_*ij*,Δ*T*_ is the covariance of the summed activity over a window of Δ*T*: 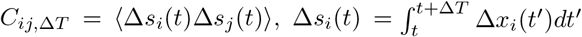. For biophysical neurons, *C*_*ij*,Δ*T*_ typically settles to its limiting value when Δ*T* > 50ms [5]. It can be shown [16] that the long time window covariance *C* (also the zero-frequency covariance, see Section 3.6) is

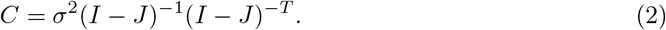

Here *I* is the identity matrix, and *A*^−1^, *A^T^* are the matrix inverse and transpose (*A^−T^* = (*A*^−1^)^*T*^). For simplicity we will set *σ*^2^ = 1 unless stated otherwise. The covariance matrix *C* can also be estimated from experimental data consisting of simultaneously recorded neurons (Methods). We consider generalizations beyond the long time window covariance in Section 3.6.

Our analysis and results start from the covariance-connectivity relation Eq. (2), which also describes, or closely approximates, the network dynamics in other models (Section 5.2) including networks of integrate-and-fire or inhomogeneous Poisson neurons [22, 23, 39, 52], fixed point activity averaged over whitened inputs, and structural equation modeling in statistics [1].

For many biological neural networks, such as cortical local circuits, the recurrent connectivity is complex and disordered. Random connectivity is a widely used minimal model to gain theoretical insights on the dynamics of neuronal networks [48, 55]. We first consider a random connectivity where

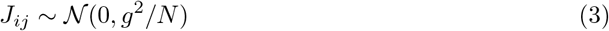

are drawn as independent and identically distributed (iid) Gaussian variables with zero mean and variance *g*^2^/*N* (referred subsequently as the *iid Gaussian connectivity*). The covariance spectrum follows directly from results in random matrices [4, 47]. We then show how to generalize to other types of random connectivity, including: those with connectivity motifs (Section 3.3), Erdos-Rényi random connectivity, networks with excitation and inhibition (Section 3.5). The theory we derived assumes the network is large and is exact as *N* → ∞, and we verify their applicability to finite-size networks numerically.

### 2.2 Covariance eigenvalues and dimensionality

Principal Component Analysis (PCA) is a widely used analysis of population dynamics, where the activity is decomposed along orthogonal patterns or Principal Components (PCs). The PCs are the eigenvectors of the covariance matrix *C* (Eq. (36)), and the associated eigenvalues *λ_i_* are positive and show the amount of activity or variance distributed along the modes. In this work, we focus on the distribution of these covariance eigenvalues, described by the (empirical) probability density function (pdf) *p_C_*(*x*) which is defined through the equality 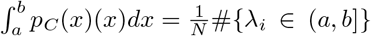 for all *a,b*. We also refer to *p_C_*(*x*) as the spectrum (which should not be confused with the frequency spectrum in Fourier transform). We will derive the limit of *p_C_*(*x*) as *N* → ∞ and study how it depends on the connectivity parameters such as *g* = *N*var(*J_ij_*).

The shape of *p_C_*(*x*) can provide important theoretical insights on interpreting PCA. For example, it can be used to separate outlying eigenvalues corresponding to low dimensional externally driven signals from small eigenvalues corresponding to fluctuations amplified by recurrent connectivity interactions [17] (Section 3.8). the spectrum is also closely related to the effective *dimension* of the population activity. In many cases, the linear span of the activity fluctuations is full rank, *N*. Nevertheless, most of the variability is embedded in a much lower dimensional subspace. A useful measure of the effective dimension, known as the participation ratio [31, 42] is given by

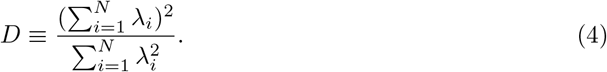

which can be calculated from the first two moments *p_C_*(*x*). We will also derive explicit expressions for *D* in random connectivity models.

## 3 Results

### 3.1 Continuous bulk spectrum with finite support

For networks with iid Gaussian connectivity (Section 2.1), there is one parameter *g* describing the overall connection strength. For stability of the fixed point and the validity of the linear response theory around it, *g* is required to be less than 1 [48]. The parameter *σ* in Eq. (2) just scales all *λ_i_* and thus is hereafter set to 1 for simplicity. Our main theoretical result is the following expression for the probability density function (pdf) of the covariance eigenvalues in the large *N* limit (Supplementary Materials),

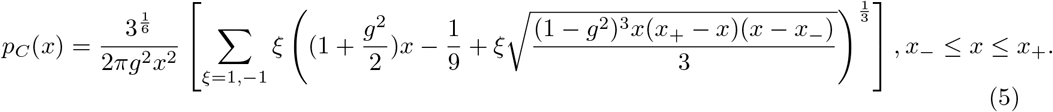

where

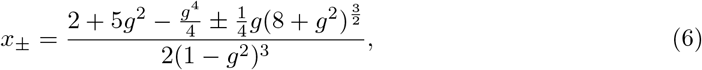

and *p_C_*(*x*) = 0 for *x* > *x*_+_ and *x* < *x*_−_. The distribution has a smooth, unimodal shape and is skewed towards the left (Fig. 1C). Near both support edges, the density scales as 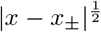 (Supplementary Materials).

**Figure 1:**
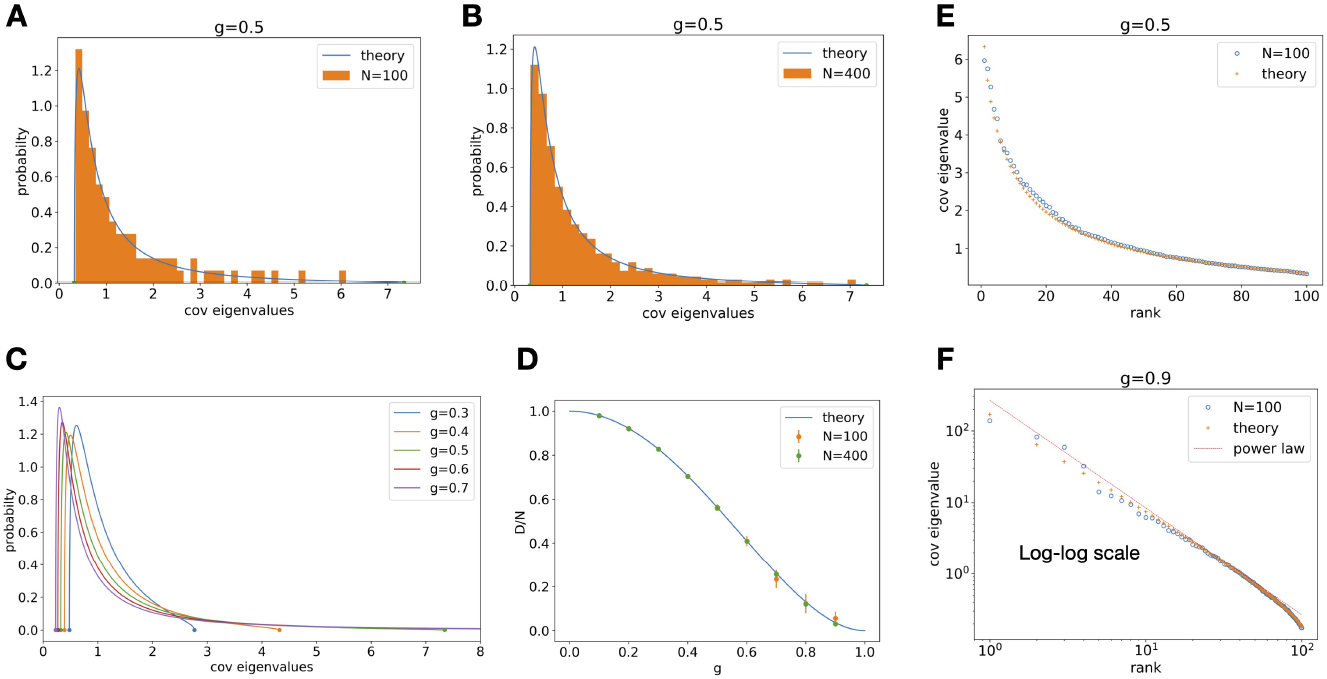
Covariance spectrum under random Gaussian connectivity. **A.** Compare theory (Eq. (5)) with finite-size network covariance using Eq. (2) at *N* = 100, *g* = 0.5. The histogram of eigenvalues is a single realization of the random connectivity. **B**. Same as A. at *N* = 400. **C**. Covariance eigenvalue distribution at various value of *g*. As *g* increases the distribution develops a long tail of large eigenvalues. **D**. Dimension (normalized by network size) vs *g*. The dots and error bars are mean and sd over repeated trials from finite-size networks (Eq. (2) and use Eq. (4)). Note some error bars are smaller than the dots **E**. Covariance eigenvalues vs. their rank (in descending order). The circles are covariance eigenvalues from a single realization of the random connectivity with *N* = 100 (Eq. (2)). The crosses are predictions based on the theoretical pdf (Eq. (5)). **F.** Same as E. but for *g* = 0.9 and on the log-log scale. The red dashed line is the power law with exponent −3/2 derived from Eq. (5), see Section 3.2

The above result for the distribution *p_C_*(*x*) follows from the derivation of the circular law distribution of the eigenvalues of the random matrix *J* [19, 4, 47, 21]. However, to the best of our knowledge, this is the first exposition of the explicit expression for the spectrum of *C*, (Eq. (5), which is essential for fitting to empirical data Section 3.8) and for the study of network dynamics. We emphasize that *p_C_*(*x*) does not have a *simple* relation to the spectrum of *J* because *J* is a non-normal matrix (i.e., *J^T^J* ≠ *JJ^T^*). This point is further elaborated in Section 3.3.2. Although the above result is derived in the large *N* limit, it matches pretty well the spectrum of *C* in networks of sizes of several hundred, as demonstrated in our numerical results, Fig. 1AB. In PCA and other analyses, the covariance eigenvalues are plotted in descending order vs. their rank [50, 35]. We can use the theoretical pdf Eq. (5) to predict this *rank plot* by numerically solving the inverse cumulative distribution function (cdf), i.e., quantile function, at probability 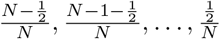. The closed form pdf (Eq. (5)) allows for using the highly efficient Newton’s method to compute the quantiles. Figure 1EF shows a good agreement between the theory Eq. (5) and a single realization of a *N* = 100 random network.

Interestingly, as we will show, the shape of the derived pdf is qualitatively different from that of the well known Marchenko–Pastur law ([32], Eq. (34) in Methods) which describes the spectrum arises from finite samples of iid Gaussian noise. This highlights that the nontrivial correlations generated by the recurrent dynamics is a distinct factor in shaping the covariance spectrum.

### 3.2 Long tail of large eigenvalues near the critical coupling

As *g* approaches the critical value of 1, the upper limit of the support *x*_+_ diverges as (1 – *g*^2^)^−3^ (Section 5.3 in Methods). This corresponds to an activity PC with diverging variance and is consistent with the stability requirement of *g* < 1. Note that the lower edge *x*_−_ is always bounded away from 0 and has a limit of 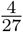 as *g* → 1. Analyzing the shape of *p_C_*(*x*) for large *x* in the critical regime *g* → 1 yields a long tail of large eigenvalues, following a power law (Fig. 2AB, Methods)

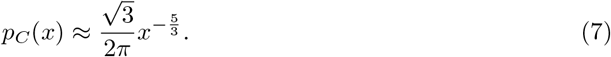

**Figure 2:**
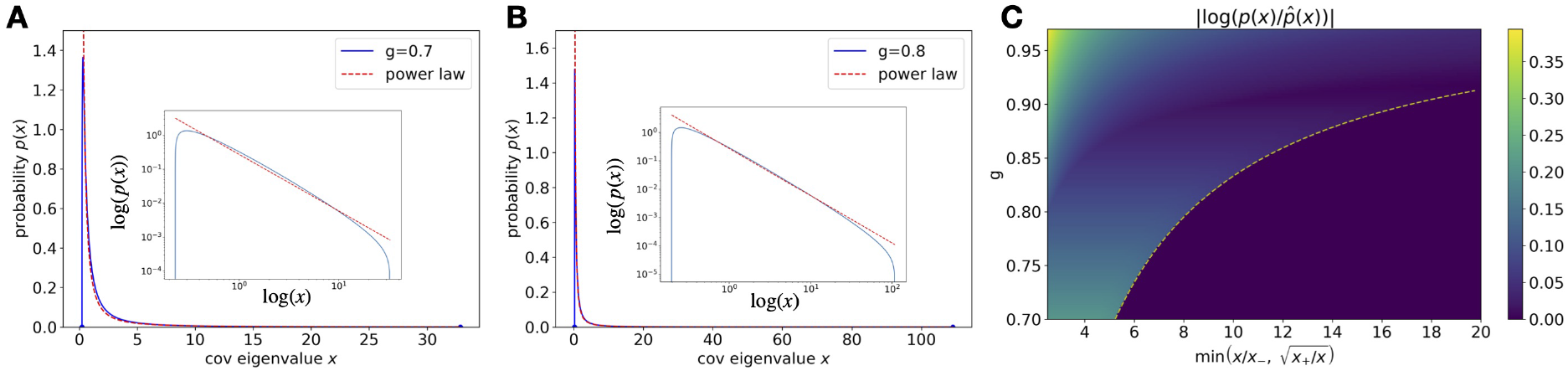
Approximate power-law tail. **A.** The exact pdf (solid line) of the covariance spectrum compared with the power-law approximation (dashed line, Eq. (7)) at *g* = 0.7. Inset shows the log-log scale. **B**. Same as A. for *g* = 0.8. The approximation improves as *g* approaches the critical value 1. **C**. The log error between the exact pdf and approximation 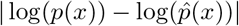 as a function of *g* and “distance” from the support edges. We quantify this “distance” as the minimum ratio of *x/x*_−_ and 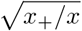 (more details and motivations in Supplementary Materials). The plot shows the log error is small when this ratio is large, which means *x* being far away from the edges. The dashed line shows the attainable region of the ratio which increases with *g*.

To better elucidate the range of validity of the above power law, we consider the regime where 1 – *g*^2^ ≪ 1 and *x* ≫ 1. Define 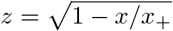 where *x*_+_ ∝ (1 – *g*^2^)^−3^ is the upper edge of the support of *p_C_*(*x*). Then,

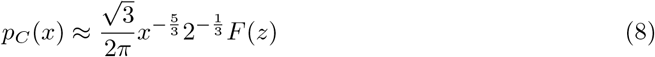

where *F*(*z*) = (1 + *z*)^1/3^ – (1 – *z*)^1/3^. Thus, far from the spectrum upper edge, *z* → 1 and we obtain Eq. (7), whereas near the upper edge *z* → 0 and 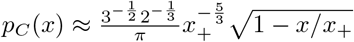, which is the expected square-root singularity near the edge.

The power-law approximation of the probability density function Eq. (5) translates to an approximation for the cumulative distribution function 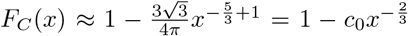. This also means a power law in the rank plot with exponent of −3/2 when connection strength *g* is close to the critical value (Fig. 1F), providing an alternative mechanism based on recurrent circuits for the experimental observations in [50, 35].

Because the probability density is small in the power-law tail, large eigenvalues can appear to be sparsely located (Fig. 1A) and potentially mistaken for statistical outliers. This underscores the importance of knowing the exact distribution and support edges for interpreting PCA results of population activity, topics which we revisit later (Fig. 8). Note that a long tail in the spectrum is a distinct feature of correlations arising from the recurrent network dynamics. For example, for the Marchenko-Pastur law that is often used for modeling empirical covariance spectra, the upper edge of its support relative to the mean is bounded by 4 (Methods). In contrast, the same ratio for *p_C_*(*x*) (Eq. (5)) can be arbitrarily large as *O*((1 – *g*^2^)^−2^) (see below for calculating the mean). This highlights the difference between covariance generated by finite samples of noise and correlations generated by the recurrent dynamics.

The long tail of the eigenvalue distribution is also reflected by a low effective dimension (Eq. (4)). In (Supplementary Materials) we show that the mean and second moment of the eigenvalue distribution above, are given by

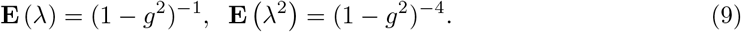

which yields for the dimension

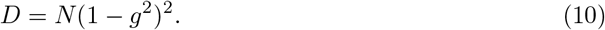

In particular, the *relative dimension* with respect to the network size *D/N* vanishes as *g* approaches 1 (Fig. 1D). In comparison, *D/N* for the Marchenko–Pastur law (Eq. (34)) is at least 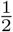.

While these low-order moments can be derived from previous methods (see e.g., [11] and Supplementary Materials), our method allows for the derivation of higher-order moments, such as,

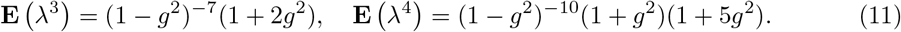

and in general,

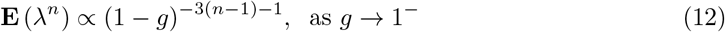

### 3.3 Impact of the asymmetry in connectivity

We next consider generalizations of random connectivity beyond the iid Gaussian model (Section 2.1). An important feature of biological neural networks is the presence of motif structures [49, 38], which correspond to overabundance of certain subgraphs, relative to their frequency in an edge-shuffled network (i.e., an iid random graph with matching connection probability). Certain motifs (e.g., diverging, converging, and chain [25]) can be shown to emerge from low rank perturbations of a random *J* hence, as explained in Section 3.4, they do not affect the bulk spectrum of *C* (see example in Fig. 5BD and Supplementary Materials).

Here we study motifs in the form of correlations between reciprocal components of *J*, which is equivalent to varying the degree of asymmetry of *J* [47]. In this case, each component of *J* is 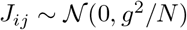 but *J_ij_* and *J_ji_* are correlated,

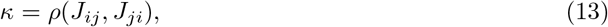

with −1 ≤ *κ* ≤ 1. All other correlations are zero.

#### 3.3.1 Symmetric and anti-symmetric random networks

First, we consider two extreme cases for the reciprocal motifs: *κ* = 1 corresponding to *J_ij_* = *j_ji_*, and *κ* = −1 corresponding to anti-symmetric matrix (or skew-symmetric *J_ij_* = – *j_ji_*). These cases are much simpler to analyze, because *J* is a *normal matrix* so *p_C_*(*x*) can be derived from the well known eigenvalue distribution of *J* ([47]). For symmetric random connectivity,

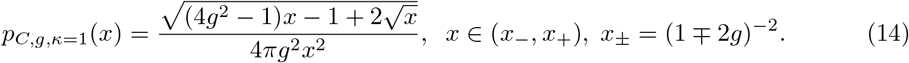

Here stability requires that 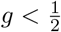. For anti-symmetric random connectivity,

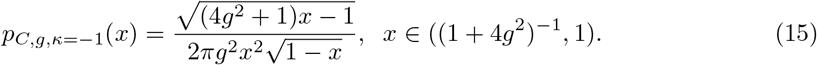

Here the network is stable for all *g*. The derivations are given in the Supplementary Materials. From the above equations, we see that *p_C_*(*x*) of the symmetric random network (Fig. 3A) has a power-law tail analogous to Eq. (7) as *g* → 1/2 (i.e., large *x*) but with a different exponent from the iid case (Eq. (7)),

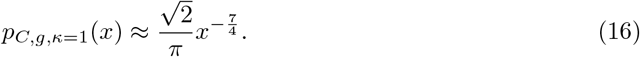

**Figure 3:**
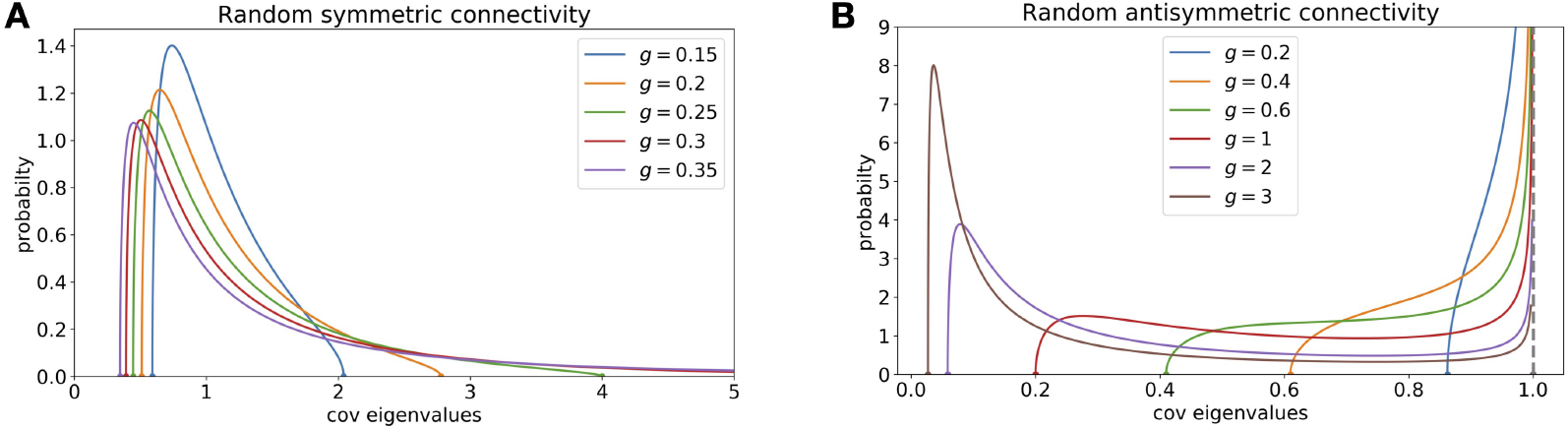
Covariance spectrum for the symmetric and anti-symmetric random connectivity. **A.** The pdf of a covariance spectrum with random symmetric *J* with different *g* (note 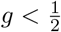 for stability). **B.** Same as A., but for random anti-symmetric *J_ij_* = – *J_ji_*. The pdf diverges at *x* = 1 as 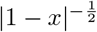.

The *p_C_*(*x*) of the anti-symmetric random network (Fig. 3B) does not have a long tail as the upper limit of the support is always 1.

#### 3.3.2 Connectivity with general asymmetry

For the Gaussian random connectivity with *κ* = *ρ*(*J_ij_*,*J_ji_*), – 1 < *κ* < 1, we have derived an implicit equation for *p_C,g,κ_*(*x*) in the large *N* limit based on the results in [47] (Eq. (S60) in Supplementary Materials). Although a closed-form expression can be derived using the root formula for quartic equations, it seems quite cumbersome, hence we show here the numerical solutions of this equation. For a fixed *g*, as *κ* increases, the distribution broadens on both sides (Fig. 4C). Intuitively, these effects might be due to the change in the critical *g* for stability, which is now given by *g_c_* = (1 + *κ*)^−1^ (based on the spectrum of *J* [47]). This motivates us to compare the distributions *p_C,g,κ_*(*x*) with the same *relative coupling strength g_r_* = *g/g_c_* = *g*(1 + *κ*), which is also the maximum real part of *J*’s eigenvalues [47]. As shown in Fig. 4D, when fixing the relative *g_r_* the distribution narrows as κ increases.

**Figure 4:**
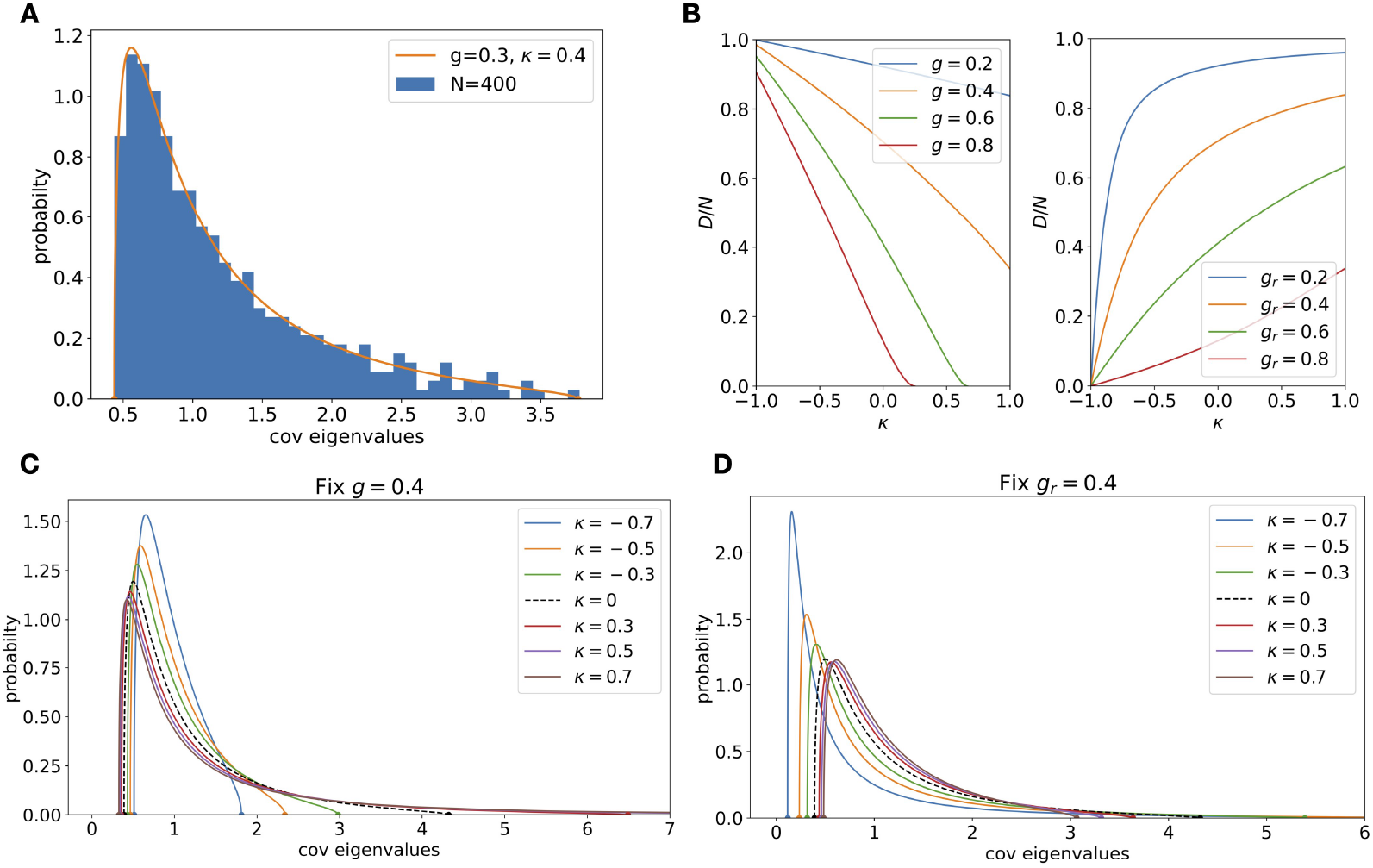
Impact of reciprocal motifs. **A**. Compare theoretical covariance spectrum for random connectivity with reciprocal motifs and a finite-size network covariance using Eq. (2)(*g* = 0.4, *κ* = 0.4, *N* = 400). **B**. The impact of reciprocal motifs on dimension for various *g_r_* = *g/g_c_* (Eq. (18)). For small *g_r_*, the dimension increases sharply with *κ*. **C**. The spectra at various *κ* while fixing *g* = 0.4. The black dashed line is the iid random connectivity (*κ* = 0). **D**. Same as C. but fixing relative *g_r_* = 0.4 to control the main effect (see text). The changes in shape are now smaller and the support narrows with increasing *κ*.

Consistent with the above results, we have shown (Supplementary Materials) that for all intermediate values of −1 < *κ* < 1, the critical covariance spectrum has an approximate power-law tail with the same exponent as the iid random case (Eq. (7))

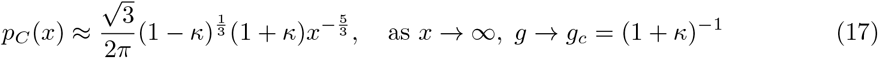

The shape changes of *p_C,g,κ_*(*x*) with reciprocal motifs are also reflected by the dimension measure, for which we derived a closed-form expression (Supplementary Materials)

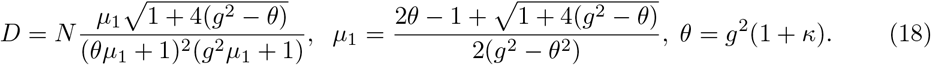

Here *μ*_1_ is the mean of the distribution. Comparing with Eq. (10), this shows the nontrivial dependence of dimension on the reciprocal motif strength *κ*. As *g* → *g_c_* = (1 + *κ*)^−1^, 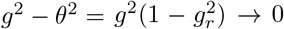. The numerator of *μ*_1_ is at least 2*θ* – 1 ≥ 2/(1 + *κ*) – 1 > 0. Therefore *μ*_1_ diverges as 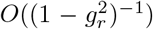. Since 1 + 4(*g*^2^ – *θ*) ≥ 1 + 4(*g*^2^ – *g*) ≥ (1 – 2/(1 + *κ*))^2^ > 0, we have *D/N* vanishes as 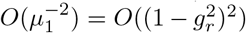. The above limits under the critical *g* are similar to the *κ* = 0 case, Eq. (10). Consistent with the shape changes, the dimension decreases with reciprocal motifs when fixing *g* and increases when fixing *g_r_* (Fig. 4B).

The general asymmetric random connectivity also provides an example of the strong effect of *J* being a non-normal matrix on the covariance spectrum. By continuity, one may expect that as *κ* decreases towards −1, the shape of *p_C,g,κ_*(*x*) will become similar to that of the anti-symmetric network *p*_*C,g,κ*=-1_(*x*), which is bimodal for sufficiently large *g* (i.e., has another peak in addition to the divergence at 1, Fig. 3B). Indeed, assuming a normal *J* predicts a covariance spectrum that is bimodal with a non-smooth peak in a large region of −1 < *κ* < 0 and *g* (Supplementary Materials). Intriguingly, the actual spectrum *p_C,g,κ_*(*x*) is unimodal for all but a minuscule region of (*κ,g*) where *κ* < −0.95 (Supplementary Materials).

### 3.4 Adding low rank connectivity structure

An important property of the spectrum of *C* is the robustness of its bulk component to the addition of low rank structured connectivity. Many connectivity structures that are important to the dynamics and function of a recurrent neuronal network can be described by a full rank random component plus a low rank component. [33, 44]. For example, such components may arise from Hebbian learning [2]. A simple case is where we add a rank *k* structured matrix that is deterministic or independent to the random component [51, 24]. As shown in the Supplementary Materials, in large networks, the bulk covariance spectrum remains unchanged, but the low rank component may give rise to at most 2*k* outlying eigenvalues. This is illustrated by the example of rank-1 perturbation to *J* with iid Gaussian entries in Fig. 5CD, where the expected location of the outliers in the covariance spectrum can be predicted analytically (Fig. 5EF, Supplementary Materials). This is in contrast to the spectrum of *J*, where the same perturbations can lead to an unbounded number of randomly located eigenvalues [41, 51] (Fig. 5AB). In sum, the bulk spectrum of covariance is robust against low rank perturbations to the connectivity. Note, however, the relevance of the bulk spectrum for the network dynamics depends on the location of outliers. Outliers to the right of the bulk spectrum may indicate potential instability of the dynamics even for *g* < 1, as discussed in the example below.

**Figure 5:**
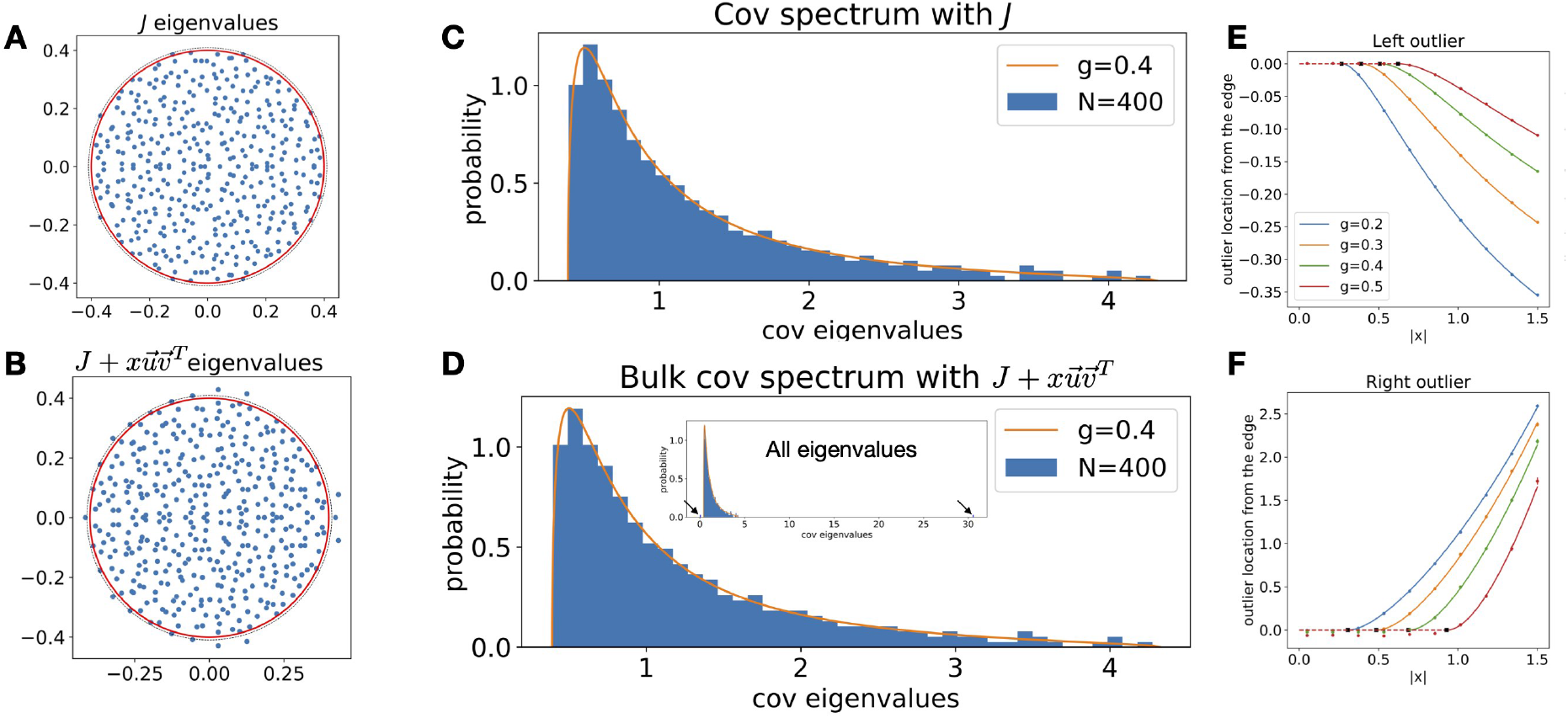
Robustness of the covariance spectrum to low rank perturbations of the connectivity. **A**. Eigenvalues of a Gaussian random connectivity *J* (Eq. (3)), *g* = 0.4, *N* = 400. As *N* → ∞, the limiting distribution of eigenvalues is uniform in the circle with radius *g* ([19] red solid line). The black dashed line is the 0.995 quantile of the eigenvalue radius calculated from 1000 realizations. **B**. Same as A. but for the rank-1 perturbed 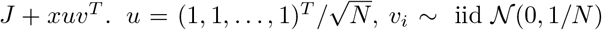 and *x* = 4.03. This example also corresponds to adding diverging motifs (Section 3.3, Supplementary Materials). **C**. The histogram of covariance eigenvalues (Eq. (2)) under the *J* in A. **D**. The bulk histogram of eigenvalues with *J* + *xuv^T^* has little change and remains well described by the Gaussian connectivity theory (red line, Eq. (5)). Besides the bulk, there are two outlier eigenvalues to the left and right (inset, arrows) **E,F**, Analytical predictions (solid and dashed lines) of the outlier locations given *g* and |*x*| when *u, v* are (asymptotically) orthogonal unit vectors that are independent of *J* (see other cases in Supplementary Materials). The y-axis is the outlier location subtracting the corresponding edge *x*_±_, Eq. (6), so it is zero for small |*x*| before the outlier emerges (dashed line). The dots are the mean of the smallest (for the left outlier) or largest (right outlier) eigenvalues averaged across 100 realizations of the random *J, N* = 4000. The errorbars are the standard error of the mean (SEM, many are smaller than the dots).

### 3.5 Sparse Excitatory–Inhibitory networks

The Gaussian random connectivity has a non-zero connection weight for all pairs of neurons with probability 1, where many biological networks are sparsely connected. In addition, each neuron has both excitatory (positive) and inhibitory (negative) weights, in contrast to many neuronal networks that obey Dale’s Law, namely all neurons are either excitatory (with all outgoing weights positive) or inhibitory (with negative outgoing weights). We consider here a simple model of E-I network, consisting of *N*/2 excitatory and *N*/2 inhibitory neurons. The probability of each connection *J_ij_* to be nonzero, which may depend on the types of neurons *i* and *j*, is *K_αβ_*/*N*, *α*, *β* = E,I. Thus, the mean number of inputs to a neuron of type *α* from a population of type *β* is *K_αβ_*/2. All excitatory non-zero connections are of strength 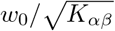 and the inhibitory connections are 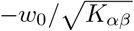. We assume that *K_αβ_* = *k_αβ_K* where *k_αβ_* = *O*(1) and *K* ≪ *N*.

To map this architecture on to the one studied above, we adopt the framework of [28] and consider the equivalent Gaussian connectivity with matching variance for each *J_ij_* which is 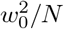 (to the leading order for *N* ≫ 1). Hence we can define the effective synaptic gain as 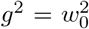 for all neurons. The mean of the connections between a presynaptic neuron of type *β* and postsynaptic *α* is 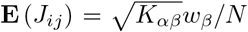 where *w_E_* = *w*_0_ and *w_I_* = –*w*_0_. Thus, we can write *J_ij_* as a zero-mean iid Gaussian matrix with uniform variance 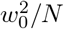 and a rank-2 matrix of the means. As stated in Section 3.4, in such a case the bulk spectrum of the neurons’ covariance matrix is the same as Eq. (5). In addition there are at most 4 outlier eigenvalues. For *K* ≫ 1, from the analysis of [28], the stability of the recurrent dynamics of a linear network with the above connectivity amounts to the requirement that all eigenvalues of the 2 × 2 matrix 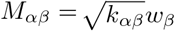 have negative real parts. Fulfilling this condition by choosing appropriate values for *k_αβ_* (see example in Fig. 6A and Supplementary Materials) guarantees that the outlier(s) due to the nonzero means are to the left of the bulk covariance spectrum so that the largest eigenvalue is *x*_+_(*g*), Eq. (6). For *K* = *O*(1), the results in [51] show that the above condition is sufficient but not necessary for stability. For example, when all *k_αβ_* are equal to *k*, which corresponds to a balance of excitation and inhibition [41], all eigenvalues of *M* are 0 and the dynamics is stable for *g* < 1 for large *N*. At the same time, there can be two outlying eigenvalues on the two sides of the bulk covariance spectrum (Fig. 6B), whose expected location can be predicted (Section 3.4 and Supplementary Materials). Several additional examples including all inhibitory networks are considered in the Supplementary Materials.

**Figure 6:**
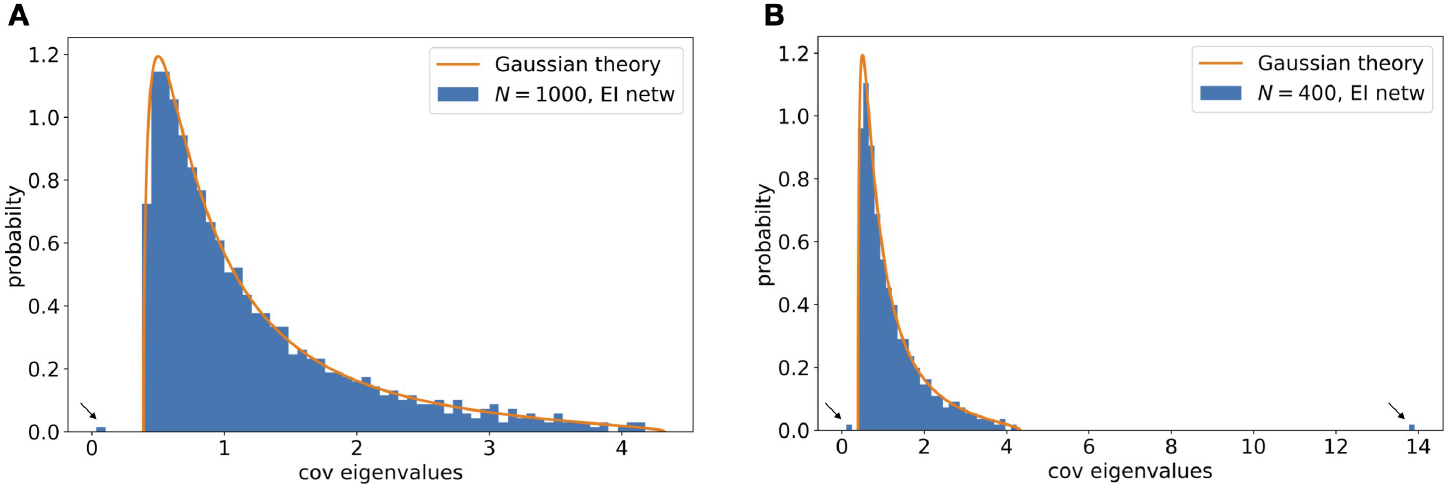
EI networks. **A**. One realization of the covariance eigenvalues by Eq. (2) with an EI network satisfying the stability condition (see text). The bulk spectrum is well described by the Gaussian random connectivity theory (solid line, Eq. (5)). There is one small outlier to the left of the bulk (arrow). The parameters are *g* = *w*_0_ = 0.4, 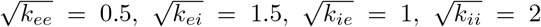, *K* = 60, *N* = 1000. To improve the accuracy of the theory to finite *K, N*, here we use a slightly modified connection weight, 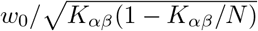, for all excitatory non-zero connections, and similarly 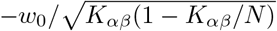 for inhibitory connections. **B**. Similar as A but for balanced EI network (see text) with *k_αβ_* = *k* = 1, *g* = 0.4, *K* = 40, *N* = 400. Note there are two outliers on both sides of the bulk.

### 3.6 Frequency dependent covariance

We have so far focused on the long time window covariance matrix. This would be especially suitable for neural activity recordings with limited temporal resolution such as calcium imaging [53]. Temporal structures of correlation beyond the slow time scale can be described by the frequency covariance matrix (or coherence matrix)

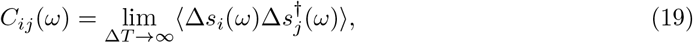

where 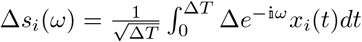 is the Fourier transform of the neural activity and *z*^†^ is the complex conjugate. *C_ij_*(*ω*) can also be calculated by the Fourier transform of the time-lagged cross-correlation functions *C_ij_*(*τ*) = 〈*x_i_*(*t*)*x_j_*(*t* – *τ*)〉 (Wiener-Khinchin theorem). Analogous to Eq. (2) *C*(*ω*) obeys [16],

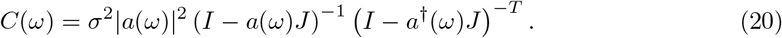

Here *z*^†^ is the complex conjugate and |*z*| is the norm for a complex *z*. The transfer function 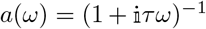 summarizes the dynamics of single neurons in the network and corresponds to a response filter of *e^−t/τ^/τ, t* > 0 for the model of Eq. (1) (see also Section 5.2). The long time window covariance we have studied corresponds to *C*(*ω* = 0) (Wiener-Khinchin theorem).

For the iid Gaussian random connectivity *J*, we can show that the spectrum of *C*(*ω*) is given by the same Eq. (5) for *C*(0) (up to a constant scaling) by replacing *g* with a frequency dependent *g*(*ω*) (compare with Eq. (3))

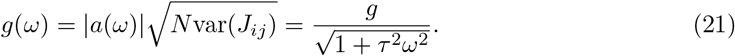

We can use Eq. (21) to study the scaling of frequency as *g* approach the critical value of 1. In many cases, we can expect that the neuronal and synaptic dynamics lead to a smooth effective low-pass filtering of the recurrent input, such that for small frequency *g*(0) – *g*(*ω*) ∝ *ω*^2^. For the specific *g*(*ω*) in Eq. (21), this can be directly verified. The low-pass filtering implies that the frequencies showing a critical covariance spectrum are those with 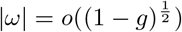.

Note, however, the simple replacement by an effective *g* may not apply to a connectivity matrix that does not have iid entries. For example, for networks with non-zero reciprocal motifs, the covariance spectrum changes qualitatively with frequency (Supplementary Materials).

### 3.7 Sampling in time and space

The theoretical spectra we have discussed are based on the exact covariance matrix (Eq. (2)). For neural data, this is equivalent to the limit of the sample covariance 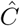 (Eq. (36)) when the number of time samples *M* is much larger than the number of neurons *N*. Note that if the activity data is first averaged or summed over a time window/bin (Δ*T* in Eq. (36)) before calculating the sample covariance, then *M* is the number of bins. However, many large-scale neural recordings are in the so-called *high dimensional* regime, where *N* and *M* are comparable, that is, the ratio *α* = *N/M* remains finite or even greater than 1 for large *N*, *M*. It is thus important to study this effect of temporal sampling on the covariance eigenvalues to better relate to experimental data [34].

We refer to 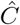 and 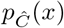 as the *time-sampled* covariance and spectrum. The relation between the original spectrum *p_C_*(*x*) and the time-sampled spectrum 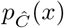 for a finite *α* ≥ 0 has been studied in [9]. The authors derived a general relation between the generating function of the eigenvalue distribution 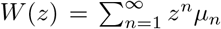, where *μ_n_* is the *n*-th moments of the eigenvalue distribution, and the counterpart 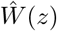 for the sampled distribution,

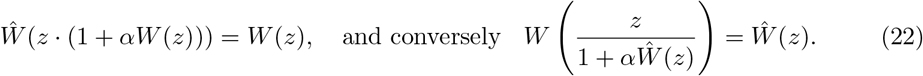

We give an alternative derivation of this result using free probability [54, 36] (Supplementary Materials), which allows us to also generalize to the spatial sampling case (see below). For simplicity, here we describe the results for 0 ≤ *α* ≤ 1. For *α* > 1 where time samples are severely limited, the spectrum of the *N* – *M* nonzero eigenvalues can be calculated with small modifications (Supplementary Materials).

One corollary of Eq. (22) is a simple formula for how the (relative) dimension changes under time sampling

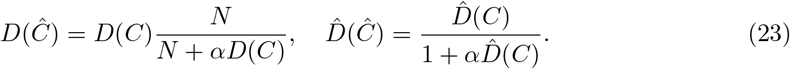

These formulas show that both *D* and 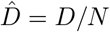 decrease with *α* (fewer time samples).

The relations Eqs. (22) and (23) apply to any covariance matrix spectrum. For example, it reproduces the time-sampled dimension derived in [34] for a different model of covariance *C*. We now apply Eq. (22) to the case of iid Gaussian connectivity to derive specific results of the time-sampled spectrum 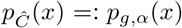. Here Eq. (22) becomes a cubic equation and can be solved analytically (Supplementary Materials). Consistent with the dimension, when *α* increases from 0 to 1, the support of the time-sampled distribution expands from both sides (Fig. 7A). In particular, for any fixed *α* < 1 (so 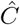 is positive definite), the left edge of the support *x*_−_ decreases with *g* but is always bounded away from 0 even as *g* → 1, where 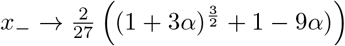 (see also figure in Supplementary Materials). Interestingly, the approximate power law of *p_C_*(*x*) (Eq. (7)) still holds under time sampling for any fixed *α* as *g* → 1 (Supplementary Materials).

**Figure 7:**
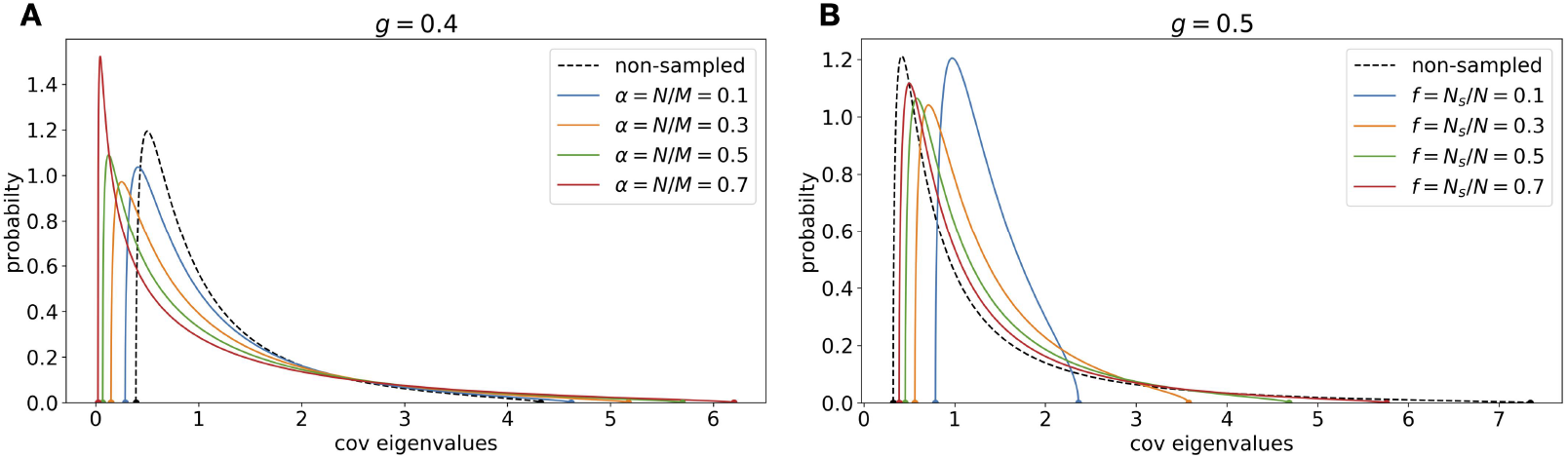
Effects of sampling in time and space on the covariance spectrum. **A**. For the iid Gaussian random connectivity, how different levels of time samples *α* change the spectrum (Eq. (22)). The non-sampled case corresponds to *α* = 0. *g* is fixed at 0.4. **B**. Same as A. but for the spatial subsampling (Eq. (S104)), at *g* = 0.5. The non-sampled case corresponds to *f* = 0.

Another challenge for fitting to neural data is that often only a subset of neurons are observed instead of the entire local recurrent network. The unobserved neurons have an impact on the dynamics and affect the eigenvalues of the observed covariance matrix. We study this by considering randomly selecting *N_s_* = *fN*, 0 < *f* ≤ 1 neurons and define their covariance 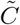 as the *space-sampled* covariance. Using the free probability approach, we derive similar results as Eqs. (22) and (23) (Supplementary Materials) and apply them to derive the spectrum and dimension for the iid Gaussian connectivity under spatial sampling. In particular, the relative dimension 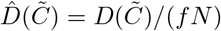 increases with spatial sampling (i.e., decrease *f*) which is consistent with the shape of the spectrum where its support narrows in (Fig. 7). Lastly, the power-law feature is also preserved under spatial sampling. For any fixed 0 < *f* ≤ 1, we show that as *g* → 1 and *x* → ∞ (see example figure with *g* close to 1 in Supplementary Materials)

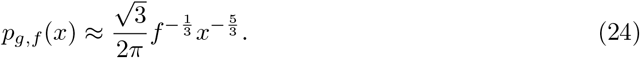

### 3.8 Fitting the theoretical spectrum to data

Our theory for the bulk covariance spectrum can be fitted to neural activity whenever the covariance eigenvalues can be calculated. The best-fitting theoretical spectrum can be found by minimizing the *L*^2^ or *L*^∞^ error between the empirical and theoretical cumulative distributions (Methods) with respect to parameters such as *g*. We note that the availability of closed-form or analytic solutions of the theoretical distributions makes this optimization highly efficient.

In many settings, the value of the baseline neuronal variance *σ*^2^ in Eq. (2) is not known. But this can be easily addressed by scaling both the observed and the predicted eigenvalues to have a mean equal to 1. After fitting the connectivity parameter *g* for the normalized eigenvalues, *σ*^2^ can then be estimated using the original means of data and theory. For our theoretical spectra, the mean *μ* of covariance eigenvalues is available in closed-form (Eqs. (9) and (18)), and the scaled pdf is easily found as *p_R_*(*x*) = *μp_C_*(*μx*).

Furthermore, the recorded neural activity is sometimes normalized for each neuron (e.g., by converting activity to z-scores). In this case, we need to analyze the eigenvalues of a *correlation matrix* whose entries are normalized as 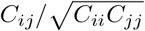. Interestingly, we found that the correlation eigenvalue distribution for our random connectivity models in the large *N* limit is the same as the rescaled *p_R_*(*x*) above. This is because the diagonal entries of *C* become uniform (thus converge to *μ*) for large *N* (Supplementary Materials).

The fitting of the spectrum is also robust to outliers in the covariance eigenvalues (Section 3.4). In Supplementary Materials we demonstrate an example where a rank-2 component is added to the covariance *C*. Since in practice the rank of the perturbation is unknown *a priori*, we use *all* eigenvalues in the fitting, and the fitted *g* is highly accurate despite the presence of outliers. We can also use the fitted *g* to help identify the outliers by separating them out based on the upper edge of *p_C_*(*x*) support [40, 17].

We conclude with a preliminary application to whole-brain calcium imaging data in larval zebrafish. In [10], activities of the majority of the neurons in the larval zebrafish brain were imaged simultaneously during presentations of various visual stimuli and grouped into *functional clusters* based on their response similarity. These clusters reveal potential neural circuits and, in some cases, reveal a good match with known brain nuclei. Here we select a few clusters that contain a large number of neurons and are anatomically localized (Fig. 8B). For each cluster, we calculate its sample correlation matrix during the spontaneous condition (no stimulus was presented) and then fitted the eigenvalues to the time-sampled spectrum with iid Gaussian connectivity (Section 3.7). Despite the simplicity of the model with only one parameter to tune, the results show good agreement with data and is significantly better than fitting using the Marchenko-Pastur law (Fig. 8C), which models spatially independent noise (Section 5.4). Therefore, our theory provides a quantitative mechanistic explanation of how a long tail of covariance eigenvalues or equivalently low dimensional activity occurs in recurrent neural circuits.

**Figure 8:**
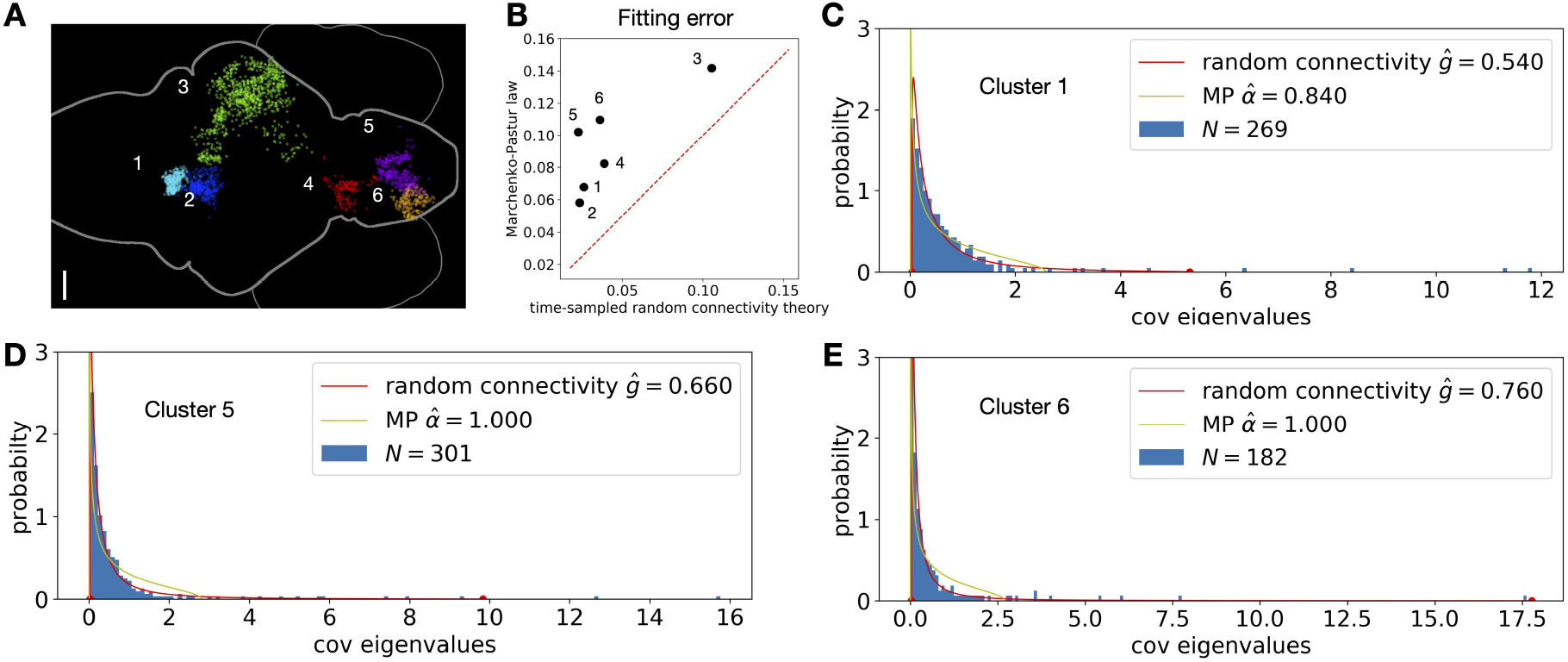
Fitting the theoretical spectrum to data. **A.** The anatomical map of neurons (dots) in the example functional clusters (different colors) across a larval zebrafish brain (scale bar is 50 *μ*m, see text and [10]). **B**. Comparing the fitting error of the time-sampled random connectivity theory (Section 3.7) and the Marchenko-Pastur law. The errors are measured by Eq. (37). The red dashed line is the diagonal. **C-E.** Comparing the fitted theoretical spectra with data (histogram). The red and yellow curves are random connectivity theory and the Marchenko-Pastur law, respectively, both with one parameter to tune. See more details in Methods. Fitting results for all other clusters are in Supplementary Materials.

## 4 Discussion

In this work, we studied the eigenvalue density distribution of the covariance matrix of a randomly connected neuronal network, whose activity is approximated as noise driven linear fluctuations around a steady state. We derived an explicit expression for eigenvalue distribution in the large-network limit analytically in terms of the statistics of the connectivity such as coupling strength and motifs. Our results also include closed-form expressions for the dimension measure generalizing known results [11]. Knowing the exact shape and support of the bulk eigenvalue distribution can facilitate separating outlying eigenvalues corresponding to low dimensional structure (coming from other unmodeled effects such as external input) [40] (Fig. S11 in Supplementary Materials). The shape of the bulk spectrum reflects structured amplification of the neuronal noise by the random recurrent interactions and is robust to low rank perturbations to the connectivity or to the activity (Supplementary Materials). As the connection strength increases towards the critical edge of stability, the spectrum exhibits a power-law tail of large eigenvalues, with exponent –5/3 in pdf (or –3/2 in eigenvalue vs. rank plot). Intriguingly, this power law persists even when the shape of the spectrum is modified by connectivity motifs or due to finite temporal and spatial sample size. In contrast, when we move away from the asymmetric, random connectivity model, the exponent of the power law (if any) becomes different: –7/4 for symmetric random connectivity (Eq. (16)), –2 for a normal connectivity *J^n^* with matching eigenvalue distribution as iid Gaussian *J* (Supplementary Materials), and –*d*/4 – 1 for a *d*-dimensional ring network (see below). Based on these results, we conjecture that a power-law tail, whenever present for any covariance spectrum, reflects the qualitative nature of the connectivity and is a robust feature that will survive both temporal and spatial sampling with the same exponent (precise statement in Section S9.3, Eq. (S108), Supplementary Materials).

The interpretation bulk spectrum corresponds to smaller eigenvalues in the PCA analysis of neural activity data, their meaning and relation to circuit connectivity. Unlike the large eigenvalues [37], the interpretation of the bulk spectrum of PCA of neural activity data has received little attention. A notable exception is a recent work [7] which studied the power law of covariance spectrum of data near criticality based on the renormalization group method. Our theory thus provides an important benchmark to compare with experimental data and advocates the bulk covariance spectrum as a powerful global description of collective dynamics in neuronal networks.

One limitation of the work is the assumed dynamic regime where fluctuations of the neuronal activity are described by the linear response theory [30, 52] around a fixed point. Extensions and comparisons to highly nonlinear activity such as chaotic dynamics [48] are left for future research. Future work could also consider more general network architectures such as multiple populations of EI networks [28] and incorporating distant dependent connectivity patterns based on known cortical microcircuit architectures [6, 8, 29].

### 4.1 Ordered vs. disorder connectivity

We have studied the covariance spectrum under random connectivity, which is used as a model for complex recurrent networks. Here we ask whether features of these spectra are distinct results of the connectivity being random. To address this, we briefly explored the covariance spectra from several widely used examples of ordered connectivity for comparison.

First, consider a ring network [6] with translation invariant long-range connections, where the connection strength between neurons depends smoothly on their distance (Fig. 9A-inset, see Methods). In the large-network limit, the covariance spectrum becomes a delta distribution at 1 with a few discretely located large eigenvalues (Fig. 9A). Next, we consider the ring network with short-range, in particular, Nearest-Neighbor (NN) connections. The covariance spectrum is now continuous (no outliers) and supported on an interval, but the pdf diverges at both edges as 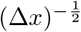 (Fig. 9B).

**Figure 9:**
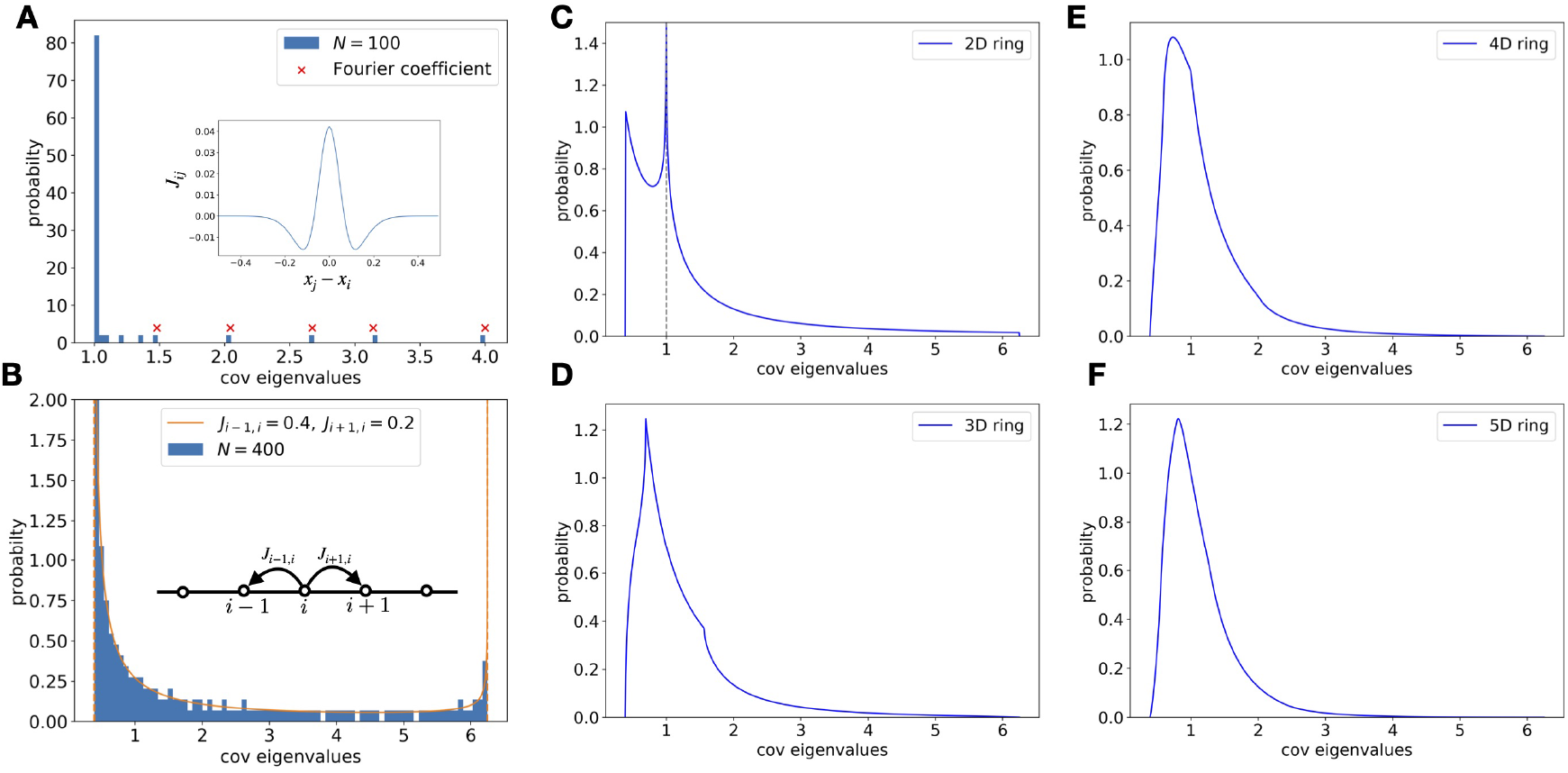
Covariance spectra under some deterministic connectivity models. **A**. Histogram of the covariance eigenvalues of a ring network with a long-range connection profile (inset, *N* = 100). Most eigenvalues are close to 1 and the rest of eigenvalues converge to discrete locations predicted by top Fourier coefficients (crosses) of the connection profile (Eq. (35)). **B**. Same as A. but for a ring network with Nearest-Neighbor connections: *J*_*i*−1_ = 0.4, *J*_*i*+1,*i*_ = 0.2. The solid line is theoretical spectrum in large *N* limit which has two diverging singularities at both support edges. The effect of such singularities is also evident in the finite-size network at *N* = 400 (a single realization). **C-F**. Higher dimensional Nearest-Neighbor ring network (*ad* = 0.6, see Methods). As the dimension increases, the singularities in the pdf become milder and less evident, and the overall shape becomes qualitatively similar to the random connectivity case (Fig. 1).

To seek further examples of ordered connectivity leading to a qualitatively similar covariance spectrum as the random connectivity, we consider the *d*-dimensional generalization of the NN ring (Methods). As dimension *d* increases, the smoothness of the pdf within and at the edges of the support increases, and the covariance spectrum becomes qualitatively similar to the random case [20] (Fig. 9C-F). Interestingly, as the connection strength approaches its critical value for stability, the covariance spectra also exhibit a power-law tail 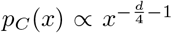 (Supplementary Materials; 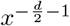 is also possible under other cases [56]). To match the exponent of the random network *d* would be 8/3 ≈ 2.67. These comparisons suggest that the covariance spectrum’s overall smooth density and long tail shape is a shared property in highly connected networks with high rank connectivity matrices, including random networks and high dimensional short-range spatially invariant networks.

## 5 Methods

### 5.1 Models of random connectivity

Here is a summary of results on various random connectivity models.

- iid Gaussian random connectivity 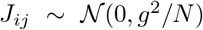: closed-form pdf and endpoints (Eq. (5)), including the frequency-dependent covariance spectrum (Section 3.6), and a power-law tail approximation (Eq. (7)).
- Gaussian random connectivity with reciprocal motifs/asymmetry *κ* = *ρ*(*J_ij_, J_ji_*) (Section 3.3): analytic solution and endpoints (quartic root, Supplementary Materials) and a power-law tail approximation (Eq. (17)). For special case of symmetric and ant-symmetric connectivity, closed-form pdf Eqs. (14) and (15), including a frequency dependent covariance spectrum (Supplementary Materials).
- Erdős-Rényi and certain EI network Section 3.5: same bulk spectrum as the iid Gaussian case.

For all cases, the mean *μ* and the dimension *D* are derived in closed-form (Eqs. (9), (10) and (18).

For simplicity, we do not require *J_ii_* to be zero (i.e., no self-coupling), but allow it, for example in the iid Gaussian model, to be distributed in the same way as other entries *J_ij_*. In large-network limit, since individual connections are weak (e.g., 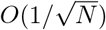), allowing this self-coupling or setting *J_ii_* = 0 does not affect the covariance spectrum (Supplementary Materials).

**Table 1:** List of notations.

| Notation | Description |
| --- | --- |
| $N$ | number of neurons |
| $C$ | covariance matrix, Eq. (2) |
| $p_C(x)$ | pdf of eigenvalues of $C$, Section 2.2 |
| $x_{\pm}$ | support edges of $p_C(x)$ |
| $J$ | matrix of connection weights Section 2.1 |
| $\sigma^2$ | variance of white noise input, Section 2.1 |
| $g$ | connection strength $\text{var}(J_{ij}) = g^2/N$, Eq. (3) |
| $g_c$ | maximum $g$ constrained by stability |
| $g_r$ | $g/g_c$ |
| $\kappa$ | $\rho(J_{ij}, J_{ji})$ reciprocal motif cumulant, Eq. (13) |
| $\mu$ | mean of eigenvalues $\frac{1}{N} \sum_{i=1}^N \lambda_i$ |
| $D$ | dimension, Eq. (4) |
| $M$ | number of time samples, Eq. (36) |
| $\alpha$ | $N/M$, Section 3.7 |
| $N_s$ | number of sampled neurons, Section 3.7 |
| $f$ | $N_s/N$, Section 3.7 |

### 5.2 Applications to alternative neuronal models and signal covariance

Although the relation between *C* and *J* (Eq. (2)) is derived here in a linear rate neuron network Eq. (1), it also arises in other models of networked systems.

#### Linearly interacting Poisson neurons

This is also called a multivariate Hawkes model [23]. This is a simple model for spiking neuron networks, but is versatile enough to capture for example the temporal spiking correlations seen in other more sophisticated nonlinear spiking neuron networks [39, 22]. A time-dependent Poisson firing rate is calculated as a filtered input spike train *S_j_*(*t*) (sum of delta functions), and spikes are then drawn as a Poisson process given *y_i_(*t**),

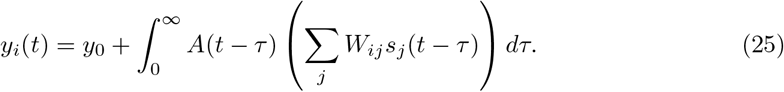

Here we consider a homogeneous network where the baseline firing rate *y*_0_ and response filter *A*(*t*) is the same for all neurons.

The exact long time window spike count covariance matrix of this network can be shown to be [23]

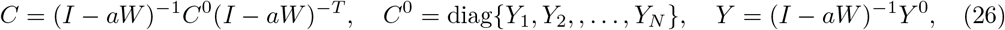

which is valid if the time varying *y_i_*(*t*) does not often become negative (for example when any negative connections *W_ij_* are small compare to *y*_0_). Here 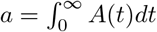, *Y*^0^ and *Y* are vectors of baseline and perturbed (with recurrent connections) firing rates of the neurons respectively. If we assume that the effective connection strength *aW* is weak so that we can approximate *Y* with *Y*^0^, then (26) becomes

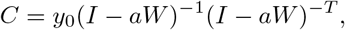

the same as Eq. (2) with *J* = *aW* (note that for Poisson process var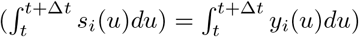.

Another condition that ensures a uniform *Y* and does not restrict connections to be weak is a *row balance* condition of W sometimes assumed in EI networks [41],

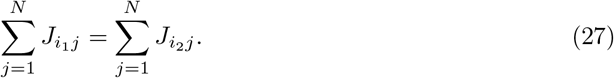

This is not unreasonable to assume, for example, considering the homeostatic mechanisms of neurons.

#### Integrate-and-Fire neurons

As shown in [52, 22] using the linear response theory [30], the covariance structure of a network of generalized integrate-and-fire (IF) neurons

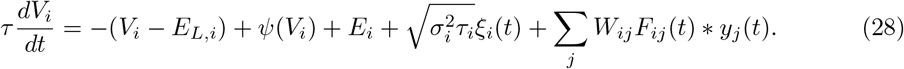

Here *V_i_* is the membrane potential and a spike is generated when *V_i_* reaches a threshold. *y_i_*(*t*) = ∑_k_ *δ*(*t* – *t_i,k_*) is the spike train, and 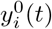 in Eq. (29) is the “unperturbed” spike train in absence of recurrent connections *W*. Different choices of *ψ*(*V*) realize types of IF neurons, such as *ψ*(*V*) = Δ_*T*_ exp((*V* – *V_T_*)/Δ_*T*_) for the exponential IF neurons. During the asynchronous firing of neurons (no strong synchronized firing across the network), Eq. (28) can be well approximated by

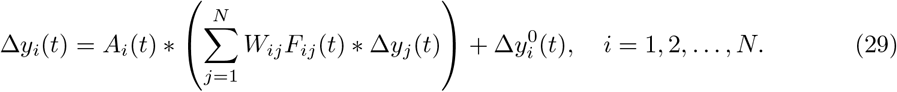

Here 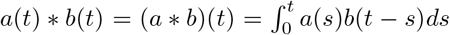 denotes convolution. *W* = {*W_ij_*} is the matrix of recurrent connection weights). *A_i_*(*t*) is the linear response kernel for neuron *i* (e.g., an exponential decay) *F_ij_*(*t*) is the temporal kernel of the synapse. For simplicity, we assume that both *A* and *F* are uniform across the network.

It it shown [52, 22] that the long time window spike count covariance matrix *C* (in fact also the frequency covariance Eq. (19)) is well approximated by

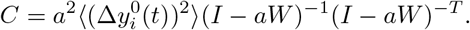

Here the scalar 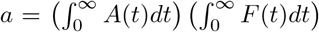 summarizes the cellular and synaptic dynamics. 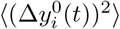 can be thought of as the baseline neuronal variance in the absence of recurrent connections (*A* = 0 in Eq. (29)). This expression of the covariance matrix again matches with Eq. (2).

#### Fixed points over whitened input

The covariance we considered so far describes the structure of fluctuations of spontaneous dynamics without or under fixed external input, often referred as *noise covariance* [3]. We can also consider a *signal covariance* in a network of firing rate neurons

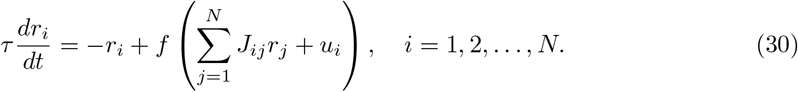

Here *u_i_* is the external input to neuron *i*. Assume the network settles to a steady state where all neurons have a firing rate *r_i_* > 0, and approximate the nonlinearity as *f*(*x*) ≈ *x*, then the fixed point firing rates are

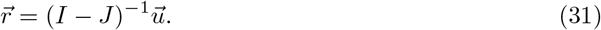

Now consider the network activity across an ensemble of input patterns, which has whitened statistics [13],

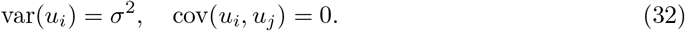

It is easy to see that the covariance of firing rates 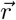 is given by *σ*^2^(*I* – *J*)^−1^(*I* – *J*)*^−T^*, which is the same as Eq.(2).

We note that Eq. (31) or equivalently 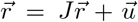 appears in broader contexts beyond neuroscience and is studied in the field of linear structural equation modeling (SEM) [1].

### 5.3 Power-law approximation of the eigenvalue distribution

The power-law property of *p_C_*(*x*) for iid *J_ij_* under critical *g* is probably known in random matrix (private communication), by results from the equivalent distribution studied in [4], although we do not know of a specific reference. We include a derivation based on the explicit expression of *p_C_*(*x*) in the Supplementary Materials that is outlined below.

First note the limits of the support edges. As *g* → 1^−^, 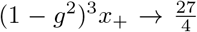. For the lower edge, 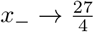 can be found by the Taylor expansion of (1 – *g*^2^) or note that (1 – *g*^2^)^3^*x*_+_*x*_−_ = 1. Consider a *x* that is far away from the support edges as *g* → 1, given the above, this means,

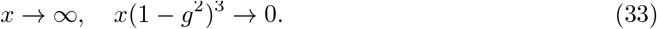

Note that since *x*_+_/*x*_−_ ~ (1 – *g*^2^)^−3^, there is plenty range of *x* to satisfy the above for strong connections when *g* is close to 1. Under these limits, Eq. (5) greatly simplifies as various terms vanish leading to (Supplementary Materials)

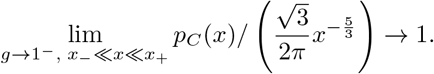

This explains the validity of the power-law approximation away from support edges. If we are only interested in the leading-order power-law tail in the critical distribution (i.e., *g* → 1^−^ and *then x* → ∞), then there is a simpler alternative derivation that can we also apply to other connectivity models (see Supplementary Materials).

### 5.4 Comparison with the Marchenko–Pastur distribution

The Marchenko-Pastur distribution is widely used for modeling covariance eigenvalues arising from noise [32, 17, 40]. It is also the limit of the time-sampled spectrum *p_g,α_*(*x*) (Fig. 7 and Section 3.7) at weak connections *g* = 0. The Marchenko-Pastur law has one shape parameter *α*. We focus on the case when the covariance is positive definite which restricts 0 < *α* < 1 (otherwise there is a delta distribution at 0) and the pdf is

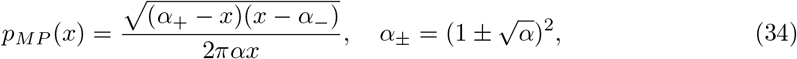

The first two moments are 1 and 1 + *α*, from which we know the dimension is 1/(1 + *α*) has a lower limit 1/2. The upper edge *α*_+_ is bounded by 4.

### 5.5 Deterministic connectivity

#### 5.5.1 Ring network with short- and long-range connections

In a ring network, neurons are equally spaced on a circle (can be physical or functional space) and neuron *i* is associated with a location *x_i_* = *i/N, i* = 0,…, *N* – 1. The connection between two neurons *j* and *i* only depends on the location difference *x_i_* – *x_j_* is therefore translation invariant.

For long-range connections, the connectivity has a shape determined by a fixed smooth periodic function *f*(*x*) on [0,1),

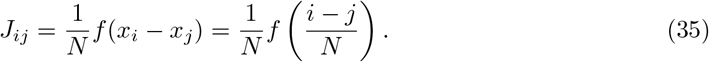

In the large-network limit, the covariance eigenvalues have an approximate delta distribution at 1 except for a finite number of discretely located larger eigenvalues (Fig. 9A). A precise statement of this result is described in Supplementary Materials. The outlying eigenvalues correspond to the leading Fourier coefficients of *f*(*x*).

For the Nearest-Neighbor (NN) connectivity, only *J*_*i*-1,*i*_ and *J*_*i*+1,*i*_ are non-zero and remain fixed as *N* → ∞.

#### 5.5.2 Multi-dimensional ring network

For a *d*-dimensional ring, the neurons are equally spaced on a *d*-dimensional lattice

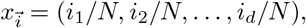

which is periodic along each coordinate. We focus on the NN connectivity where each neuron is connected to 2*d* neighboring neurons with strength 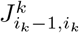 and 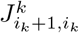 along direction *k*. We show that the probability density function at both support edges scales as 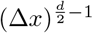 (for comparison, the random network edges scale as 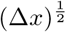). This means for dimension *d* ≥ 2, there is no singularity at the support edges (Fig. 9).

To characterize the shape of the covariance spectrum (Fig. 9C-F), we further simplify by setting 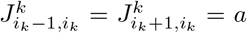 (see also Supplementary Materials for motivations based on 1D ring) and analytically derived *p_C_*(*x*) (Supplementary Materials). For small dimensions *d* ≤ 3, there are distinct “inflection points” within the support. As *d* increases, this non-smooth feature becomes less evident and becomes hard to identify in empirical eigenvalue distributions from a finite-size network (not shown).

### 5.6 Fitting the theoretical spectrum to data

For neural activity data, *C* can be calculated from a large number of time samples of binned spike count *s_i_*(*t*) (assuming bin size is Δ*T* large),

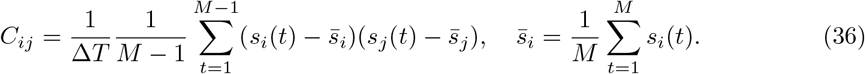

For calcium imaging data, the fluorescence is approximately integrating the spikes over the indicator time constant. So we can still apply Eq. (36) by plugging in the fluorescence signal in place of *s_i_*(*t*) to calculate the covariance *C* (omit the constant factor Δ*T* which does not affect fitting to the theory, Section 3.8).

We fit the theoretical spectrum to empirical eigenvalues by finding the connectivity parameter *g* that minimizes the error between the *cumulative distribution functions* (cdf) 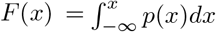. This avoids issues such as binning when estimating the probability density function from empirical eigenvalues. We numerically integrate the theoretical pdf (Eq. (5)) to get its pdf. As seen below, the theoretical cdf only needs to be calculated at the empirical eigenvalues.

Motivated by methods of hypothesis testing on distributions, we measure the *L*^2^ norm cdf error using the Cramer-von Mises statistic

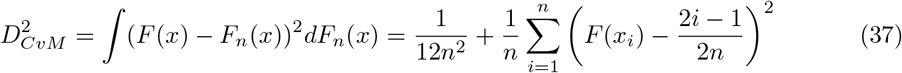

Here *n* is the number of samples and *x_i_* are the *i*-th empirical eigenvalues. Alternatively, the error can also be measure under *L*^∞^ norm based on the Kolmogorov-Smirnov statistic

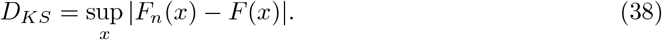

where *x_i_* are samples. Our code implements both measures.

In Fig. 8B-E, we fit the time-sampled theoretical spectrum with iid Gaussian connectivity (Section 3.7) to calcium imaging data in larval zebrafish [10]. The theoretical spectrum (once normalized by the mean, see Section 3.8) depends on two parameters *g* and *α*, but the latter is fixed to be *N/M* based on the data. Here *N* is the number of neurons in a cluster, and *M* is the number of time frames used in calculating the sample correlation matrix (Eq. (36), the calcium fluorescence Δ*F/F* traces of each neuron are normalized to z-scores [10]). *g* is then optimized to minimize the Cramer-von Mises error (Eq. (37)) between the data. The largest eigenvalue for each cluster is often much larger than the rest and is thus removed before the fitting. For comparison, we fit the same data to the Marchenko-Pastur law (Section 5.4) whose shape depends on the parameter *α*. Here we allow *α* to vary so that both models (random connectivity and MP law) have one parameter to be optimized to fit data.

### 5.7 Code

Codes for theoretical spectra, generating covariance for finite-size networks, fitting to the empirical spectrum, and making all figures are available at https://github.com/huyu00/netw_cov_spectrum

The larval zebrafish whole-brain calcium imaging data and functional clustering code are available from [10].

## Acknowledgements

This work has been partially supported by the Swartz Program in Theoretical Neuroscience at Harvard, an NIH grant from the NINDS (1U19NS104653), and the Gatsby Charitable Foundation. YH acknowledges support from HKUST. The authors would like to thank Eric Shea-Brown, Stefano Recanatesi, and Mark Reimers for helpful discussions, and Zhigang Bao for comments about the random matrix literature.

## References

[1] C. Amendola, P. Dettling, M. Drton, F. Onori, and J. Wu. “Structure Learning for Cyclic Linear Causal Models”. arXiv (2020), pp. 1–19.

[2] D. J. Amit, H. Gutfreund, and H. Sompolinsky. “Storing Infinite Numbers of Patterns in a Spin-Galss Model of Neural Networks”. Phys. Rev. Lett. September (1985), pp. 1530–1533.

[3] B. B. Averbeck, P. E. Latham, and A. Pouget. “Neural correlations, population coding and computation.” Nature reviews. Neuroscience 7.5 (2006), pp. 358–366.

[4] Z. D. Bai. “Circular law”. Annals of Probability 25.1 (1997), pp. 494–529.

[5] W. Bair, E. Zohary, and W. T. Newsome. “Correlated firing in macaque visual area MT: time scales and relationship to behavior.” The Journal of neuroscience: the official journal of the Society for Neuroscience 21.5 (2001), pp. 1676–1697.

[6] R. Ben-Yishai, R. Lev Bar-Or, and H. Sompolinsky. “Theory of orientation tuning in visual cortex”. Proceedings of the National Academy of Sciences of the United States of America 92.9 (1995), pp. 3844–3848.

[7] S. Bradde and W. Bialek. “PCA Meets RG”. Journal of Statistical Physics 167.3-4 (2017), pp. 462–475.

[8] Y. Burak and I. R. Fiete. “Accurate path integration in continuous attractor network models of grid cells”. PLoS Computational Biology 5.2 (2009).

[9] Z. Burda, A. Görlich, A. Jarosz, and J. Jurkiewicz. “Signal and noise in correlation matrix”. Physica A: Statistical Mechanics and its Applications 343.1-4 (2004), pp. 295–310.

[10] X. Chen et al. “Brain-wide Organization of Neuronal Activity and Convergent Sensorimotor Transformations in Larval Zebrafish”. Neuron 100.4 (Nov. 2018), 876–890.e5.

[11] D. Dahmen, S. Grün, M. Diesmann, and M. Helias. “Second type of criticality in the brain uncovers rich multiple-neuron dynamics”. Proceedings of the National Academy of Sciences of the United States of America 116.26 (2019), pp. 13051–13060.

[12] M. Farrell, S. Recanatesi, T. Moore, G. Lajoie, and E. Shea-Brown. “Recurrent neural networks learn robust representations by dynamically balancing compression and expansion”. bioRxiv (2019), pp. 1–20.

[13] D. J. Field. “Relations between the statistics of natural images and the response properties of cortical cells”. Journal of the Optical Society of America A 4.12 (1987), p. 2379.

[14] S. Fusi, E. K. Miller, and M. Rigotti. “Why neurons mix: High dimensionality for higher cognition”. Current Opinion in Neurobiology 37 (2016), pp. 66–74.

[15] J. A. Gallego, M. G. Perich, S. N. Naufel, C. Ethier, S. A. Solla, and L. E. Miller. “Cortical population activity within a preserved neural manifold underlies multiple motor behaviors”. Nature Communications 9.1 (2018), pp. 1–13.

[16] C. Gardiner. Handbook of Stochastic Methods for Physics, Chemistry and the Natural Sciences. Berlin: Springer-Verlag, 2009, p. 464.

[17] M. Gavish and D. L. Donoho. “The Optimal Hard Threshold for Singular Values is 4/sqrt(3)”. IEEE Transactions on Information Theory 60.8 (2014), pp. 5040–5053.

[18] I. Ginzburg and H. Sompolinsky. “Theory of correlations in stochastic neural networks”. Physical Review E 50.4 (1994), pp. 3171–3191.

[19] V. L. Girko. “Circular law”. Theory of probability and its applications (1983).

[20] J. a. Goldberg, U. Rokni, and H. Sompolinsky. “Patterns of Ongoing Activity and the Functional Architecture of the Primary Visual Cortex”. Neuron 42 (2004), pp. 489–500.

[21] F. Götze and A. Tikhomirov. “The circular law for random matrices”. Annals of Probability 38.4 (2010), pp. 1444–1491.

[22] D. Grytskyy, T. Tetzlaff, M. Diesmann, and M. Helias. “A unified view on weakly correlated recurrent networks”. Frontiers in Computational Neuroscience 7.October (2013), pp. 1–19.

[23] A. G. Hawkes. “Spectra of some self-exciting and mutually exciting point processes”. Biometrika 58.1 (1971), pp. 83–90.

[24] R. A. Horn and C. R. Johnson. Matrix Analysis. Cambridge University Press, 1990.

[25] Y. Hu, S. L. Brunton, N. Cain, S. Mihalas, J. N. Kutz, and E. Shea-brown. “Feedback through graph motifs relates structure and function in complex networks”. Physical Review E 062312 (2018), pp. 1–25.

[26] Y. Hu, J. Trousdale, K. Josić, and E. Shea-Brown. “Local paths to global coherence: Cutting networks down to size”. Physical Review E - Statistical, Nonlinear, and Soft Matter Physics 89.3 (2014), pp. 1–16.

[27] Y. Hu, J. Trousdale, K. Josić, and E. Shea-Brown. “Motif statistics and spike correlations in neuronal networks”. Journal of Statistical Mechanics: Theory and Experiment 2013.03 (2013), P03012.

[28] J. Kadmon and H. Sompolinsky. “Transition to chaos in random neuronal networks”. Physical Review X 5.4 (2015), pp. 1–28.

[29] R. B. Levy and A. D. Reyes. “Spatial profile of excitatory and inhibitory synaptic connectivity in mouse primary auditory cortex”. Journal of Neuroscience 32.16 (2012), pp. 5609–5619.

[30] B. Lindner, B. Doiron, and A. Longtin. “Theory of oscillatory firing induced by spatially correlated noise and delayed inhibitory feedback”. Physical Review E 72.6 (2005), p. 061919.

[31] A. Litwin-Kumar, K. D. Harris, R. Axel, H. Sompolinsky, and L. F. Abbott. “Optimal Degrees of Synaptic Connectivity”. Neuron 93.5 (Mar. 2017), 1153–1164.e7.

[32] V. A. Marchenko and L. A. Pastur. “Distribution of eigenvalues for some sets of random matrices”. Mathematics of the USSR-Sbornik 1.4 (1967), p. 457.

[33] F. Mastrogiuseppe and S. Ostojic. “Linking Connectivity, Dynamics, and Computations in Low-Rank Recurrent Neural Networks”. Neuron 99.3 (2018), 609–623.e29.

[34] L. Mazzucato, A. Fontanini, and G. La Camera. “Stimuli reduce the dimensionality of cortical activity”. Frontiers in Systems Neuroscience 10.FEB (2016).

[35] L. Meshulam, J. L. Gauthier, C. D. Brody, D. W. Tank, and W. Bialek. “Coarse graining, fixed points, and scaling in a large population of neurons”. Physical Review Letters 123.17 (2019), p. 178103.

[36] J. A. Mingo and R. Speicher. Free Probability and Random Matrices. New York, NY, 2017.

[37] M. Okun et al. “Diverse coupling of neurons to populations in sensory cortex”. Nature 521.7553 (2015), pp. 511–515.

[38] R. Perin, T. Berger, and H. Markram. “A synaptic organizing principle for cortical neuronal groups”. P Natl Acad Sci USA 108.13 (2011), pp. 5419–5424.

[39] V. Pernice, B. Staude, S. Cardanobile, and S. Rotter. “How Structure Determines Correlations in Neuronal Networks”. PLoS Computational Biology 7.5 (2011), e1002059.

[40] A. Peyrache, K. Benchenane, M. Khamassi, S. I. Wiener, and F. P. Battaglia. “Principal component analysis of ensemble recordings reveals cell assemblies at high temporal resolution”. Journal of Computational Neuroscience 29.1-2 (2010), pp. 309–325.

[41] K. Rajan and L. F. Abbott. “Eigenvalue spectra of random matrices for neural networks”. Phys Rev Lett 97 (2006), p. 188104.

[42] K. Rajan, L. F. Abbott, and H. Sompolinsky. “Interactions between Intrinsic and Stimulus-Evoked Activity in Recurrent Neural Networks”. Ed. by M. Ding and D. Glanzman. Oxford University Press, 2011. Chap. The Dynamic Brain: An Exploration of Neuronal Variability and Its Functional Significance, pp. 65–82.

[43] S. Recanatesi, G. K. Ocker, M. A. Buice, and E. Shea-Brown. “Dimensionality in recurrent spiking networks: Global trends in activity and local origins in connectivity”. PLoS Computational Biology 15.7 (2019), pp. 1–29.

[44] A. Rivkind and O. Barak. “Local Dynamics in Trained Recurrent Neural Networks”. Physical Review Letters 118.25 (2017), pp. 1–5.

[45] P. T. Sadtler, K. M. Quick, M. D. Golub, S. M. Chase, S. I. Ryu, E. C. Tyler-Kabara, B. M. Yu, and A. P. Batista. “Neural constraints on learning”. Nature 512.7515 (2014), pp. 423–426.

[46] J. D. Semedo, A. Zandvakili, C. K. Machens, B. M. Yu, and A. Kohn. “Cortical Areas Interact through a Communication Subspace”. Neuron 102.1 (2019), 249–259.e4.

[47] H. J. Sommers, a. Crisanti, H. Sompolinsky, and Y. Stein. “Spectrum of large random asymmetric matrices”. Physical Review Letters 60.19 (1988), pp. 1895–1898.

[48] H. Sompolinsky, a. Crisanti, and H. J. Sommers. “Chaos in random networks”. Phys. Rev. Lett. 61.3 (1988), pp. 259–262.

[49] S. Song, P. Sjöström, M. Reigl, S. Nelson, and D. Chklovskii. “Highly nonrandom features of synaptic connectivity in local cortical circuits”. PLoS Biol 3.3 (2005), e68.

[50] C. Stringer, M. Pachitariu, N. Steinmetz, M. Carandini, and K. D. Harris. “High-dimensional geometry of population responses in visual cortex”. Nature (2019).

[51] T. Tao. “Outliers in the spectrum of iid matrices with bounded rank perturbations”. Probability Theory and Related Fields 155.1-2 (2013), pp. 231–263.

[52] J. Trousdale, Y. Hu, E. Shea-Brown, and K. Josić. “Impact of Network Structure and Cellular Response on Spike Time Correlations”. PLoS Comput Biol 8.3 (2012), e1002408.

[53] N. Vladimirov, Y. Mu, T. Kawashima, D. V. Bennett, C.-T. Yang, L. L. Looger, P. J. Keller, J. Freeman, and M. B. Ahrens. “Light-sheet functional imaging in fictively behaving zebrafish”. Nature Methods 11.9 (2014), pp. 883–884.

[54] D. Voiculescu. “Multiplciation of certain noncommuting random variables”. Journal of Operator Theory 18 (1987), pp. 2223–2235.

[55] C. van Vreeswijk and H. Sompolinsky. “Chaos in Neuronal Networks with Balanced Excitatory and Inhibitory Activity”. Science 274.5293 (Dec. 1996), 1724 LP–1726.

[56] X. Wang. “Volumes of Generalized Unit Balls Published by: Mathematical Association of America Linked references are available on JSTOR for this article: Volumes of Generalized Unit Balls”. Mathematics Magazine 78.5 (2005), pp. 390–395.

[57] D. J. Watts and S. H. Strogatz. “Collective dynamics of ‘small-world’ networks.” Nature 393.6684 (1998), pp. 440–2.

[58] L. Zhao, B. Beverlin, T. Netoff, and D. Q. Nykamp. “Synchronization from Second Order Network Connectivity Statistics”. Front Comput Neurosci 5 (Jan. 2011), pp. 1–16.

