## Supplementary Materials for "The spectrum of covariance matrices of randomly connected recurrent neuronal networks"

**S6 Sparse connectivity and Excitatory–Inhibitory networks 22**

**S7 Frequency dependent covariance 23**

**S8 Time-sampled covariance matrix 24**

**S10 The spectrum of the correlation matrix 31**

**S11 Additional results on fitting the theoretical spectrum to data** 31

**S12 Ordered and deterministic connectivity** 33

### S1 Derivation of the covariance spectrum Eq. (5)

Here we consider a network with Gaussian random connectivity $J$, $J_{ij} \sim \mathcal{N}(0, \frac{g^2}{N})$. The edges $J_{ij}$ are independent except for possible nonzero $\kappa = \rho(J_{ij}, J_{ji})$ (the case of Eq. (5) corresponds to $\kappa = 0$). We want to calculate the eigenvalue distribution of $C$ in the large $N$ limit. This distribution can be easily obtained if we know the eigenvalue distribution $p_\eta(x)$ of the (scaled) *precision matrix* $P$

$$P = g^{-2}C^{-1} = (\eta I - J^0)^\dagger(\eta I - J^0), \quad \eta = 1/g, \;\; J^0 = J/g. \tag{S1}$$

Here $A^\dagger$ is the conjugate transpose as we will allow $\eta$ to be complex. To proceed, we can use the result in [10], which calculated the logarithmic potential of $p_\eta(x)$

$$\Phi(\epsilon, \eta) = -\int_0^\infty \log|\epsilon + x| p_\eta(x)dx, \quad \epsilon > 0. \tag{S2}$$

It is related to the Stieltjes transform of $p_\eta(x)$ by

$$\Delta_\eta(z) = -\frac{d\Phi}{d\epsilon}(-z). \tag{S3}$$

The Stieltjes transform is an important generating function defined as

$$\Delta_\eta(z) = \int_{\mathbb{R}} \frac{1}{x - z} p_\eta(x)dx, \tag{S4}$$

and is valid for all $z \in \mathbb{C}$ excluding the support of $p_\eta(x)$ (on the real line). Importantly, the pdf can be recovered via

$$p_\eta(x) = \lim_{\epsilon \to 0^+} \frac{\Delta_\eta(x - \mathbb{i}\epsilon) - \Delta_\eta(x + \mathbb{i}\epsilon)}{2\pi\mathbb{i}}. \tag{S5}$$

As shown in [10], in the large $N$ limit, $\Phi$ can be calculated using the replica method with an order parameter $\sigma$ satisfying

$$\frac{\epsilon}{\sigma^2} = \frac{1}{1+\sigma} - \frac{a^2}{(\sigma+1+\kappa)^2} - \frac{b^2}{(\sigma+1-\kappa)^2}, \quad \frac{d\Phi}{d\epsilon} = -\frac{1}{\sigma}. \tag{S6}$$

Here $\eta = a + b\mathbb{i}$. Together with Eq. (S3), we arrive at an equation for $\Delta_\eta(z)$,

$$z\Delta_\eta^2 = -\frac{1}{1+\Delta_\eta^{-1}} + \frac{a^2}{(1+\Delta_\eta^{-1}+\kappa)^2} + \frac{b^2}{(1+\Delta_\eta^{-1}-\kappa)^2}. \tag{S7}$$

Note that although the definition of $\Phi$ (Eq. (S2)) restricts $z = -\epsilon < 0$, the Stieltjes transform is analytic in $z$ outside of the support of $p_\eta(x)$, so the validity of Eq. (S7) can be extended to any complex $z$ not on the support.

For the iid Gaussian random connectivity with $\kappa = 0$ and $\eta = 1/g$, Eq. (S7) becomes a cubic equation for $\Delta_\eta$,

$$\Delta_\eta^3 + 2\Delta_\eta^2 + \frac{z + 1 - |\eta|^2}{z}\Delta_\eta + \frac{1}{z} = 0. \tag{S8}$$

Equation (S8) is also derived in [1] using a different method. Equation (S8) has three roots which can be either 3 real ones or 1 real and 2 complex ones. The roots and their type are closely related to $p_\eta(x)$ and its support. This is discussed in [1] and also [4]. Note that for our purpose, we restrict to $0 < g < 1$ or $|\eta| > 1$ so that the network is stable, which further simplifies the scenarios, and the exact support edges of $p_\eta(x)$ are [1]:

$$x_{1,2}^\eta = \frac{1}{8|\eta|^2}\left(-1 + 20|\eta|^2 + 8|\eta|^4 \pm \left(1 + 8|\eta|^2\right)^{\frac{3}{2}}\right). \tag{S9}$$

This translates to the support edges of $C$'s spectrum (Eq. (6)),

$$x_\pm = \frac{2 + 5g^2 - \frac{g^4}{4} \pm \frac{1}{4}g(8 + g^2)^{\frac{3}{2}}}{2(1 - g^2)^3}. \tag{S10}$$

Here $x_\pm$ are the left and right edges, respectively. They are also the roots of a quadratic polynomial based on the determinant of cubic equations (see Eq. (S12)).

Within the support, the relevant roots of $\Delta_\eta$ for the density $p_\eta(x)$ are the complex conjugate pair, and the inverse Stieltjes transform (Eq. (S5)) becomes $\frac{1}{\pi}|\mathrm{Im}(\Delta_\eta(z = x))|$. To derive an explicit expression for $p_\eta(x)$, we use the root formula for cubic equations (Cardano formula) and remember that we only need the imaginary part of the complex roots. Let $u = 1/x$, $u(1 - |\eta|^2) = A < 0$, $u|\eta|^2 = B > 0$, then $y = \Delta_\eta + 1$ satisfies

$$y^3 - y^2 + Ay + B = 0.$$

Now define the standard intermediate variables in the Cardano formula,

$$p = A - \frac{1}{3}, \quad q = B + \frac{A}{3} - \frac{2}{27}.$$

One can show that for $x_1^\eta < x < x_2^\eta$,

$$p < 0, \quad q > 0, \quad \frac{q^2}{4} + \frac{p^3}{27} > 0.$$

These help to select the correct square roots and cubic roots in the formula. In particular, in the expressions for intermediate variables $s, t$

$$s, t := -\left(\frac{q}{2} \pm \sqrt{\frac{q^2}{4} + \frac{p^3}{27}}\right)^{\frac{1}{3}}.$$

Here all quantities that are being taken roots of are positive and we take the positive root. Finally, the absolute value of the imaginary part of the complex root of $\Delta_\eta$ is $\frac{\sqrt{3}}{2}|s - t|$, which gives the eigenvalue density of the precision matrix $P$.

$$p_\eta(x) = \frac{3^{\frac{1}{6}}}{2\pi x^{\frac{1}{3}}}\left[\sum_{\xi=1,-1} \xi \left(\eta^2 + \frac{1}{2} - \frac{x}{9} + \frac{\xi}{\sqrt{3}}\sqrt{2\eta^4 + 5\eta^2 - \frac{1}{4} - \frac{(\eta^2 - 1)^3}{x} - \eta^2 x}\right)^{\frac{1}{3}}\right]. \tag{S11}$$

Using $p_C(x) = p_\eta\left(\frac{1}{g^2 x}\right)\frac{1}{g^2x^2}$, the eigenvalue distribution of the covariance matrix is then

$$p_C(x) = \frac{3^{\frac{1}{6}}}{2\pi g^2 x^2}\left[\sum_{\xi=1,-1}\xi\left((1+\frac{g^2}{2})x - \frac{1}{9} + \xi\sqrt{\frac{x}{3}}\sqrt{(2+5g^2-\frac{g^4}{4})x-(1-g^2)^3x^2-1}\right)^{\frac{1}{3}}\right], \tag{S12}$$

Note that the square root in the formula above becomes zero if and only if $x$ equals to the support edges $x_\pm$. Therefore $p_C(x)$ can also be written as (Eq. (5))

$$p_C(x) = \frac{3^{\frac{1}{6}}}{2\pi g^2 x^2}\left[\sum_{\xi=1,-1}\xi\left((1+\frac{g^2}{2})x - \frac{1}{9} + \xi\sqrt{\frac{(1-g^2)^3x(x_+-x)(x-x_-)}{3}}\right)^{\frac{1}{3}}\right].$$

#### S1.1 Density near the support edges

Let

$$A = (1+\frac{g^2}{2})x - \frac{1}{9}, \quad B = \sqrt{(1-g^2)^3x(x_+-x)(x-x_-)/3}.$$

As $x \to x_-$, $A$ approaches a non-zero constant, while $B \to 0$, thus $p_C(x) \to 0$. Using the identity $p - q = \frac{p^3-q^3}{p^2+q^2+pq}$ with $p = (A+B)^{\frac{1}{3}}$, $q = (A-B)^{\frac{1}{3}}$,

$$\begin{aligned} p_C(x) &\approx \frac{3^{\frac{1}{6}}}{\pi g^2 x_-^2}\left((1+\frac{g^2}{2})x_- - \frac{1}{9}\right)^{-\frac{2}{3}}\sqrt{\frac{(1-g^2)^3x_-(x_+-x_-)(x-x_-)}{3}} \\ &= c_1\sqrt{x-x_-}, \end{aligned}$$

holds to the leading order.

Similarly, as $x \to x_+$,

$$\begin{aligned} p_C(x) &\approx \frac{3^{\frac{1}{6}}}{\pi g^2 x_+^2}\left((1+\frac{g^2}{2})x_+ - \frac{1}{9}\right)^{-\frac{2}{3}}\sqrt{\frac{(1-g^2)^3x_+(x_+-x)(x_+-x_-)}{3}} \\ &= c_1\sqrt{x_+-x}. \end{aligned}$$

#### S1.2 Approximate power-law tail

Below we explain an alternative derivation of the leading-order power-law approximation for the iid Gaussian random connectivity as $g \to 1^-$. We will later use similar arguments for other cases of random connectivity.

Note that for any $g > 0$, the distribution of eigenvalues of $P = (\eta I - J^0)^\dagger(\eta I - J^0)$ is always well defined and finite supported on $[0, \infty)$. Set $g = 1$ in Eq. (S8)

$$\Delta_\eta(\Delta_\eta + 1)^2 = -\frac{1}{\alpha}. \tag{S13}$$

For $\alpha > 0$, there is a critical value above which the equation has 3 real roots. Importantly, this transition occurs at a stationary point of the left-hand side of Eq. (S13), which gives $\Delta_\eta = -1/3$. The corresponding $\alpha$ value is then $27/4$, which is the limiting upper bound for $p_\eta(x)$. For any $0 < \alpha < 27/4$, there is only one real root and thus $p_\eta(x)$ is non-zero so the limiting lower bound for $p_\eta(x)$ is zero.

As $\alpha \to 0$, the magnitude of the roots goes to infinity. The equation thus approximates to

$$\Delta_\eta^3 = -\frac{1}{\alpha}.$$

The imaginary part of the complex pair of roots gives the density as (divergent at 0)

$$p_\eta(x) \approx \frac{\sqrt{3}}{2\pi} x^{-\frac{1}{3}}, \quad \text{as } x \to 0^+.$$

Using the reciprocal relation of $P$ and $C$ (Eq. (S1)), this edge density of $p_\eta(x)$ translates to the power-law tail of $p_C(x)$ (Eq. (7)).

To evaluate the power-law approximation numerically, we consider the "distance" of $x$ from the boundary described by the ratios $x/x_-$ and $x_+/x$. The results in the main text show that the log error $|\log(p(x)) - \log(\hat{p}(x))|$, $\hat{p}(x) = \frac{\sqrt{3}}{2\pi} x^{-\frac{5}{3}}$ vanishes in the limit when both ratios go to infinity. When the ratios are large but finite, we can plot the log-error as a function of the two ratios (data not shown). We observe that the log-errors are similar when $x/x_- \approx \sqrt{x_+/x}$, meaning at the same ratio distance from the right edge leads to larger error than the left edge (Fig. 2AB). Therefore, in Fig. 2C, we plot the log-error against $\min(x/x_-, \sqrt{x_+/x})$.

##### S1.3 Diagonal entries of connectivity $J$

In our random connectivity models, we have allowed non-zero $J_{ii}$ (i.e., self-coupling) for simplicity. For example, in the iid Gaussian model, $J_{ii}$ has the same $\mathcal{N}(0, g^2/N)$ distribution as other entries. In the large-network limit, since individual connections are weak (e.g., $O(1/\sqrt{N})$), either allowing this self-coupling or setting $J_{ii} = 0$ should not affect the covariance spectrum. We verified this claim numerically (Fig. S1)

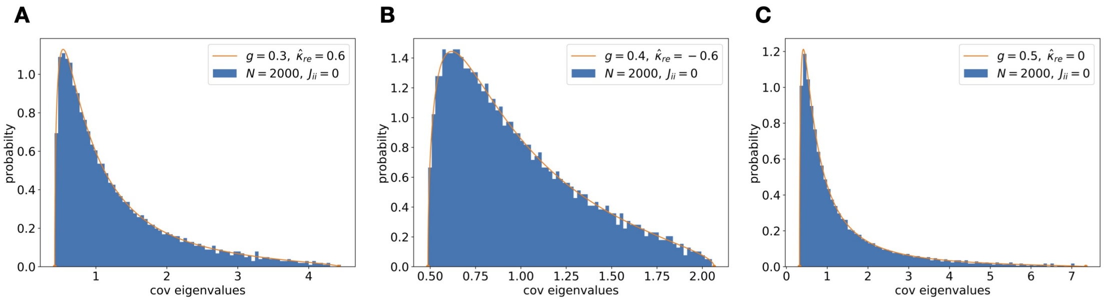

Figure S1: Setting $J_{ii} = 0$ does not affect the bulk spectrum in the large $N$ limit. Here are examples with various values of $g$ and the reciprocal motif $\hat{\kappa}_{re}$.

#### S2 Dimension and moments

The dimension $D$ (Eq. (4)) can be calculated from the first two moments of the eigenvalue distribution. We derive the moments using the equation for the Stieltjes transform and the moment generating function as explained in detail in Section S5. Let $M(z) = \sum_{n=1}^{\infty} \mu_n z^n$ where $\mu_n = \mathbf{E}(\lambda)^n$ is the $n$-th moment of the covariance eigenvalues. For iid Gaussian connectivity, as a special case of Eq. (S65) derived in Section S5,

$$\frac{1}{z} = -\frac{1}{M(g^2 M + z)} + \frac{1}{(g^2 M + z)^2}. \tag{S14}$$

Multiplying by the denominators $M(g^2M+z)^2$ and comparing the coefficient of $z$, $z^2$ gives Eq. (9),

$$\mathbf{E}(\lambda) = (1-g^2)^{-1}, \quad \mathbf{E}\left(\lambda^2\right) = (1-g^2)^{-4},$$

and hence $D = N(1-g^2)^2$. By similarly comparing higher-order terms, we can obtain higher-order moments (which are always explicitly expressed in terms of lower moments), for example,

$$\mathbf{E}\left(\lambda^3\right) = (1-g^2)^{-7}(1+2g^2), \quad \mathbf{E}\left(\lambda^4\right) = (1-g^2)^{-10}(1+g^2)(1+5g^2).$$

These results suggest that as $g \to 1^-$, $\mathbf{E}(\lambda^n) \propto (1-g)^{-3(n-1)-1}$ (Eq. (11)). We will derive this by establishing a general relation between a power-law pdf and the moments. Suppose the pdf $p_\delta(x) \to c_0 x^{-\beta}$ and the its left and right support edges $x_-(\delta) \to a > 0$, $x_+(\delta) \propto \delta^{-\alpha}$, as parameter $\delta \to 0$. Here $0 < \alpha$, $1 < \beta < 2$. The $n$-th moment is approximated as

$$\mathbf{E}(\lambda^n) \to \int_{x_-}^{x^+} x^n c_0 x^{-\beta} dx = \frac{c_0 \delta^{-\alpha(-\beta+k+1)}}{-\beta+n+1} - \frac{c_0 a^{-\beta+k+1}}{-\beta+n+1}. \tag{S15}$$

Note the second term is a constant, therefore

$$\mathbf{E}(\lambda^n) \propto \delta^{-\alpha(-\beta+k+1)} \text{ as } \delta \to 0. \tag{S16}$$

For the covariance spectrum with iid Gaussian random connectivity, $\delta = 1-g$, $\alpha = 3$ (Eq. (6)) and $\beta = \frac{5}{3}$ (Eq. (7)). Plugging these into Eq. (S16) gives Eq. (11). Interestingly, one can also use Eq. (S16) to solve for $\alpha$, $\beta$ and hence the power-law property, from the results of just two moments (assume such a power law near the critical parameter exists).

**First two moments as a corollary of results in [3]** The Eq. (2) and (3) in [3] give the mean and standard deviation of $C_{ij}$, copied below for easy reference,

$$\bar{C}_{ij} = (1-\mu)^{-1} D_\lambda (1-\mu^T)^{-1}, \quad D_\lambda = D/(1-\lambda_{\max,J}),$$

$$\delta C_{ij} = D_\lambda \sqrt{\frac{1+\delta_{ij}}{N}\left(\left(\frac{1}{1-\lambda_{\max,J}^2}\right)^2 - 1\right)}.$$

For our iid Gaussian random connectivity, mean connection $\mu$ is a zero matrix, the maximum real part of connectivity eigenvalues $\lambda_{\max,J} = g$, and the unperturbed variance $D = 1$. Using these results, the mean of the covariance eigenvalues is

$$\frac{1}{N}\mathrm{tr}(C) = \bar{C}_{ii} = \frac{1}{1-g^2}. \tag{S17}$$

For the second moment of the covariance eigenvalues, separate it into diagonal and off-diagonal terms

$$\frac{1}{N}\mathrm{tr}(C^2) = \frac{1}{N}\sum_{1\le i,j\le N} C_{ij}^2 = \frac{1}{N}\left(N\overline{C_{ii}^2} + N(N-1)\overline{C_{i\neq j}^2}\right). \tag{S18}$$

Note $\overline{C_{ii}^2} = D_\lambda^2 + \frac{2}{N}D_\lambda^2\left(\left(\frac{1}{1-\lambda_{\max,J}^2}\right)^2 - 1\right)$ and $\overline{C_{i\neq j}^2} = 0 + \frac{1}{N}D_\lambda^2(\left(\frac{1}{1-\lambda_{\max,J}^2}\right)^2 - 1)$. Plugging these into Eq. (S18) leads to

$$\begin{aligned}\frac{1}{N}\mathrm{tr}(C^2) &= (1-g^2)^{-2} + \frac{N+1}{N}((1-g^2)^{-4} - (1-g^2)^{-2}) \\ &\to (1-g^2)^{-4}, \quad \text{as } N \to \infty.\end{aligned}$$

These are the same results for the first two moments as Eq. (9).

#### S3 Robustness to low rank perturbations of connectivity and activity

Here we show that a low rank perturbation to the covariance $C$ or to the connectivity matrix $J$ does not change the continuous part, i.e., bulk spectrum, of the covariance matrix in large networks. Moreover, the number of the outliers aside from the bulk spectrum is bounded by the rank or two times the rank of the perturbation. Note that this is in contrast to the effect of similar low rank perturbations on the spectrum of connectivity $J$, which can result in an unbounded number of outliers even with a rank-1 perturbation [11]. We later explained how to predict the location of the outlying eigenvalues for large $N$ (Sections S3.2 and S3.3).

**Proposition S3.1.** *Assume that the covariance matrix $C$ has a limiting spectrum that has a continuous pdf and is finitely supported as network size $N \to \infty$. $C^0$ is a rank $k$ symmetric matrix, with $k$ being fixed. Then the eigenvalues of $\tilde{C} = C + C^0$ have the same limiting continuous bulk spectrum as $C$ as $N \to \infty$, with at most $k$ eigenvalues outside of the support of the bulk.*

*Proof.* Assume $C^0$ has $a$ positive eigenvalues and $b$ negative eigenvalues, $a + b \leq k$. Using Weyl's inequality on the eigenvalue changes when perturbing a symmetric matrix [5],

$$\lambda_{i-b} \leq \tilde{\lambda}_i \leq \lambda_{i+a}, \quad b+1 \leq i \leq N-a, \tag{S19}$$

where $\lambda_i$ and $\tilde{\lambda}_i$ are eigenvalues of $C$ and the perturbed $\tilde{C}$ sorted in ascending order. If $C$ has a limiting continuous spectrum with a finite support, then by Eq. (S19) there can be at most $a + b \leq k$ eigenvalues that lie outside of this support as $N \to \infty$. Moreover, the difference between the empirical distributions are bounded by

$$\sup_x |F_{\tilde{C}}(x) - F_C(x)| \leq \frac{k}{N}.$$

Therefore, the spectrum of $\tilde{C}$ converges to the same limit of $C$ as $N \to \infty$. □

**Proposition S3.2.** *Assume that the covariance matrix $C$, defined by connectivity $J$ as Eq. (2), has a limiting spectrum that has a continuous pdf and is finitely supported as network size $N \to \infty$. $\tilde{J}$ is a rank $k$ perturbation of $J$. Assume $I - \tilde{J}$ is invertible and $k$ is fixed, then the eigenvalues of its corresponding covariance $\tilde{C}$ have the same continuous limiting bulk spectrum as $C$, with at most $2k$ eigenvalues outside of the support of the bulk.*

*Proof.* Let $\tilde{J} = J + uv^T$ be the rank $k$ perturbation of $J$, where matrices $u_{N\times k}, v_{N\times k}$ have linearly independent columns.

$$\tilde{C}^{-1} = (I - \tilde{J})^T (I - \tilde{J}) = (I - J)^T (I - J) - vu^T (I - J) - (I - J)^T uv^T + v(u^T u)v^T.$$

Let $\tilde{u} = (I - J)^T u$, $U_{N\times 2k} = [\tilde{u}, v]$, we have

$$\tilde{C}^{-1} = C^{-1} + U \begin{bmatrix} 0 & -I \\ -I & u^T u \end{bmatrix} U^T. \tag{S20}$$

Using the Woodbury formula,

$$\tilde{C} = C + CU \left( \begin{bmatrix} u^T u & I \\ I & 0 \end{bmatrix} - U^T C U \right)^{-1} U^T C. \tag{S21}$$

Note that $\begin{bmatrix} u^T u & I \\ I & 0 \end{bmatrix} - U^T C U$ in the above equation must also be invertible, otherwise we can find a null direction for $\tilde{C}^{-1}$ leading to a contradiction with $I - J$ being invertible. Equation (S21) shows that $\tilde{C}$ is a perturbation of $C$ with rank at most $2k$. The claim is thus obtained using Prop. S3.1. □

#### S3.1 Gaussian random connectivity with second-order motifs

Following [6], we study a connectivity model with jointly Gaussian distributed entries that can produce all four types of second-order motifs (composed of two edges, Fig. S2). The entries of $J$ in this model can be written as

$$J_{ij} = a_i + b_j + \tilde{J}_{ij}, \;\; 1 \le i, j \le N, \tag{S22}$$

where $a_i$, $b_j$, $\tilde{J}_{ij}$ are zero-mean Gaussian variables that have the same variance across $i, j$

$$\mathrm{var}(a_i) = a^2, \;\; \mathrm{var}(b_j) = b^2, \;\; \mathrm{var}(\tilde{J}_{ij}) = c^2, \;\; 1 \le i, j \le N, \tag{S23}$$

and are independent except for

$$\mathrm{cov}(a_i, b_i) = k, \;\; \mathrm{cov}(\tilde{J}_{ij}, \tilde{J}_{ji}) = r, \;\; i \ne j. \tag{S24}$$

Such a structure and correlation introduces the second-order motifs [14, 7] to $J$,

$$\begin{aligned} &\kappa_{div} = \mathrm{cov}(J_{ik}, J_{jk}) = b^2, \;\; \kappa_{con} = \mathrm{cov}(J_{ki}, J_{kj}) = a^2, \;\; \kappa_{ch} = \mathrm{cov}(J_{ik}, J_{kj}) = k, \\ &\kappa_{re} = \mathrm{cov}(J_{ij}, J_{ji}) = 2k + r, \quad i \ne j \ne k. \end{aligned} \tag{S25}$$

We can achieve various second-order *motif cumulants* $\kappa_*$ by adjusting $\{a^2, b^2, c^2, k, r\}$. The relation between these covariances and the motifs can be seen in the network where $J_{ij}$ take binary values of $w_0$ or 0 [14, 7]. For example, in large networks $\frac{N^3}{w_0^2}\mathbf{E}\left(J_{ik}J_{jk}\right)$ is the number of diverging motifs. Note that $\mathbf{E}\left(J_{ij}\right)/w_0$ is the connection probability and $\frac{N^3}{w_0^2}\mathbf{E}\left(J_{ik}\right)\mathbf{E}\left(J_{jk}\right) = Np^2$ is the expected number of diverging motifs in a matching Erdős-Rényi (ER) random graph. Therefore $\kappa_{div}$ is proportional to the deviations of the diverging motif frequency from the (iid) ER graph.

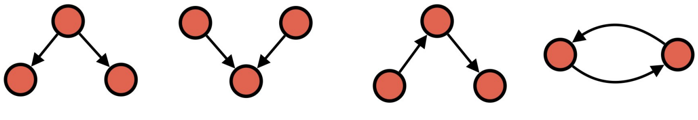

Figure S2: Four types of second-order motifs.

From Eq. (S22) it is clear that $J$ can be viewed as $\tilde{J}$, which only has reciprocal motifs, perturbed by a rank-2 matrix $ae^T + eb^T$, $a = (a_i)$, $b = (b_i)$, $e = (1, 1, \ldots, 1)^T$. As an example, the rank-1 perturbation in Fig. 5B,D corresponds to a connectivity with diverging motifs and the values of $x$ and $\hat{\kappa}_{div}$ can be converted using the formulas below. According to Prop. S3.2, the continuous spectrum of the covariance based on $J$ is the same as that for $\tilde{J}$ with at most four outliers for large $N$. This justifies focusing on studying connectivity with reciprocal motifs.

One can normalize $\kappa_*$ to correlations as in Eq. (13), $\hat{\kappa}_*$, by dividing $\mathrm{var}(J_{ij}) = a^2 + b^2 + c^2$. In particular, $\hat{\kappa}_{re} = \mathrm{cov}(J_{ij}, J_{ji})/\mathrm{var}(J_{ij}) = \rho(J_{ij}, J_{ji})$ ($\kappa$ in the main text). Note that when applying results based on the reciprocal-only model Eq. (13) to a general $J$ from Eq. (S22), the $\kappa = \hat{\kappa}_{re}$ and $g$ in the formulas (e.g., Eq. (18)) are referring to the $\tilde{J}$ component and need to be calculated from motifs of $J$

$$g^2(\tilde{J}) = g^2(J)(1 - \hat{\kappa}_{div}(J) - \hat{\kappa}_{con}(J)), \quad \hat{\kappa}_{re}(\tilde{J}) = \frac{\hat{\kappa}_{re}(J) - 2\hat{\kappa}_{ch}(J)}{1 - \hat{\kappa}_{div}(J) - \hat{\kappa}_{con}(J)}. \tag{S26}$$

This indicates that the diverging, converging, and chain motifs do not directly affect the bulk covariance spectrum, but channel their effect through changing the normalized reciprocal motifs.

#### S3.2 Outliers in the covariance spectrum when perturbing $C$

We consider a rank-1 perturbation to a covariance $C$ generated with iid Gaussian connectivity (Eq. (2)) with a fixed $0 \leq g < 1$ $\tilde{C} = C + xuu^T$ (note the perturbation has to be symmetric). Here $u$ is a $N \times 1$ vector with iid entries $u_i \sim \mathcal{N}(0, 1/N)$, and is independent of $J$. $x > 0$ is a scalar kept fixed as $N \to \infty$. We will see that this makes the outlying eigenvalue (if any) converge to a finite value.

According to Prop. S3.1, there is at most one outlying eigenvalue $z$. Since $z$ is outside the support of the spectrum of $C$, $zI - C$ is invertible, we have

$$\begin{aligned} 0 &= \det(zI - C - xuu^T) = (\det(zI - C))^{-1} \det(I - xuu^T(zI - C)^{-1}) \\ &= (\det(zI - C))^{-1} \det(1 - xu^T(zI - C)^{-1}u). \end{aligned} \tag{S27}$$

Here we used Sylvester's identity $\det(I - AB) = \det(I - BA)$. So we need to solve

$$1 = xu^T(zI - C)^{-1}u. \tag{S28}$$

Next, we use the following fact without proof: as $N \to \infty$, $u^T(zI - C)^{-1}u$ converges to its expectation (over random $J$ and $u$). The expectation is

$$\begin{aligned} &\mathbf{E}(u^T(zI - C)^{-1}u) = \mathbf{E}\left(\mathrm{tr}((zI - C)^{-1}uu^T)\right) \\ &= \mathbf{E}_J(\mathrm{tr}((zI - C)^{-1})\mathbf{E}_u(uu^T)) = \frac{1}{N}\mathbf{E}_J(\mathrm{tr}((zI - C)^{-1})) = -\Delta_C(z) \end{aligned}$$

Therefore, Eq. (S28) becomes

$$\Delta_C(z) = -\frac{1}{x}, \tag{S29}$$

where $\Delta_C(z)$ is the Stieltjes transform of the spectrum of $C$. For small $x > 0$, there is no outlier in $\tilde{C}$, and for sufficiently large $x$, one outlier appears to the right of the support of $p_C(x)$. The value of $x$ for this transition can be found using Eq. (S29) as

$$x_{\min} = -1/\Delta_C(z = x_+). \tag{S30}$$

Here $x_+$ is the right edge of the support (Eq. (6)). To determine $x_{\min}$, first, $\Delta_P(x_-^P)$, which is evaluated at the left edge of $P = \frac{1}{g^2}C^{-1}$, can be solved from Eq. (S61) (with $\kappa = 0$)

$$\Delta_P(x_\pm^P) = \frac{3 \mp \sqrt{1 + 8/g^2}}{4(1/g^2 - 1)}. \tag{S31}$$

Here we also include the case for the right edge $x_+^P$. Then using the following relation

$$\Delta_P\left(\frac{1}{g^2 z}\right) = -g^2 z^2 \Delta_C(z) - g^2 z, \tag{S32}$$

we can obtain $\Delta_C(z = x_+)$ and then $x_{\min}$.

To solve for the outlier location $z$, plugging Eqs. (S29) and (S32) into Eq. (S8), we get a fourth-order equation for $z$. For $x > x_{\min}$, there are two real roots and the smaller one corresponds to the outlying eigenvalue $z$. We confirm our analytical predictions of the outlier with generating covariance matrices of finite-size networks using Eq. (2) (Fig. S3A)

The results above can be generalized to the cases of independent rank-$k$ perturbation: $C + \sum_{i=1}^{k} x_i u_i u_i^T$, where $u_i$ has iid $\mathcal{N}(0, 1/N)$ entries and is independent of $u_{j\neq i}$ and $J$. As $N \to \infty$, $u_i^T(zI - C)^{-1}u_j \to 0$. This ensures we get a diagonal matrix in the generalized version of Eq. (S27). Therefore, the (at most) $k$ outlying eigenvalues can be solved separately for each $x_i$ in the same way as $k = 1$. In particular, we use the generalized results for $k = 2$ to determine the outliers in Fig. S11 (Fig. S3B).

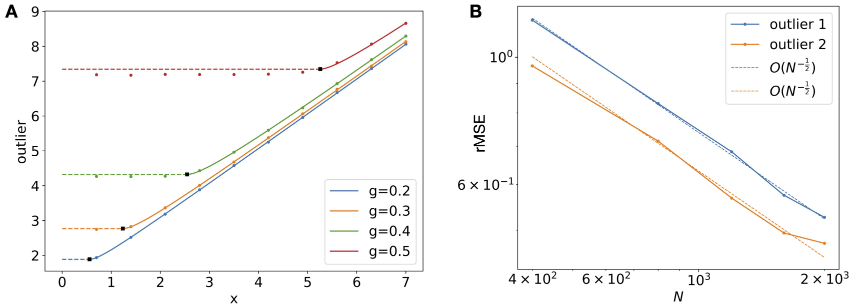

Figure S3: **A**. The outlier eigenvalue as a function of the perturbation strength $x$ for different $g$ (colors) when perturbing the random network covariance with a rank-1 matrix (see text). The analytical predictions (dash and solid curves) match well with simulations of the largest eigenvalue at $N = 1000$ (dots, averaged over 400 realizations of random $J$ and $u$). For $x < x_{\min}(g)$, there is no outlier, therefore the largest eigenvalue is close to the right edge of the support of the bulk spectrum (dashed line). The predicted transition $x_{\min}$ to having an outlier is marked by the black square. **B**. The root-mean-square error (rMSE) of predicting the two outliers (corresponding to $\sigma_{1,2}^2$ respectively) in Fig. S11. The simulations is at $N = 400, 800, 1200, 1600, 2000$ and is averaged over 1000 realizations. The dashed line is the fitted square-root decay with $N$ on the log-log plot.

##### S3.3 Outliers in the covariance spectrum when perturbing $J$

We consider a rank-1 perturbation of the iid Gaussian connectivity $J$: $\tilde{J} = J + xuv^T$, where $x$ is a fixed scalar and $u, v$ are vectors with unit 2-norm (as $N \to \infty$). They can be either deterministic or with iid $\mathcal{N}(0, 1/N)$ entries (independent within each vector - more on the relation between $u, v$ later) as long as being independent with $J$. This creates at most two outlying eigenvalues in the covariance $\tilde{C}$ (defined by Eq. (2) with $\tilde{J}$). We can equivalently find the outliers in $\tilde{C}^{-1}$ and then their reciprocals are outliers in $\tilde{C}$. From Eq. (S20), and using similar arguments as Eq. (S27), the outliers $z$ of $\tilde{C}^{-1}$ satisfy

$$\begin{aligned} 0 &= \det\left( \begin{bmatrix} 0 & -x \\ -x & x^2 \end{bmatrix}^{-1} - U^T(zI - C^{-1})^{-1}U \right) \\ &= \det\left( \begin{bmatrix} 1 + u^T(I-J)(zI - C^{-1})^{-1}(I-J)^T u & \frac{1}{x} + u^T(I-J)(zI - C^{-1})^{-1}v \\ \frac{1}{x} + v^T(zI - C^{-1})^{-1}(I-J)^T u & v^T(zI - C^{-1})^{-1}v \end{bmatrix} \right) \end{aligned} \tag{S33}$$

Here $U_{N\times 2} = [(I-J)^T u, v]$. Similarly as in Section S3.2, we will use the following fact without proof: as $N \to \infty$, the terms in the $2 \times 2$ matrix of Eq. (S33) converge to their expectations. At least one of these expectations is $O(1)$, so the outlier $z$ to the leading order of $N$ is determined by plugging these expectations in Eq. (S33).

The expectation of the $(2,2)$-entry is analogous to Eq. (S28): $-\Delta_{C^{-1}}(z)$. For the $(1,1)$-entry, note that the distribution of $(I-J)(zI-C^{-1})^{-1}(I-J)^T$ is invariant under any orthogonal similarity transform (see also Section S9). Therefore its expectation must be a scalar matrix $\lambda I$.

$$\begin{aligned}&\mathbf{E}(u^T(I-J)(zI-C^{-1})^{-1}(I-J)^T u) = \mathrm{tr}(\mathbf{E}_J((I-J)(zI-C^{-1})^{-1}(I-J)^T)uu^T)\\&= \lambda u^T u = \lambda = \frac{1}{N}\mathbf{E}_J(\mathrm{tr}((I-J)(zI-C^{-1})^{-1}(I-J)^T))\end{aligned} \tag{S34}$$

It is straightforward to show this as a matrix identity,

$$\mathrm{tr}((I-J)(zI-C^{-1})^{-1}(I-J)^T) = \mathrm{tr}(z(zI-C^{-1})^{-1} - I). \tag{S35}$$

This shows the expectation of $(1,1)$-entry of Eq. (S33) is $-z\Delta_{C^{-1}}(z)$.

To proceed, we need to calculate the off-diagonal entry of Eq. (S33) (the two entries are the same $(1,2) = (2,1)$). It turns out its expectation depends on the angle/overlap between $u, v$ and has a significant impact on the covariance outliers. For simplicity, we consider two extreme cases: (i) $u, v$ being orthogonal or $u, v$ being independent random vectors with iid $\mathcal{N}(0, 1/N)$ entries; (ii) $u = v$. General cases with $r := u^T v$ being a fixed value in $[-1, 1]$ can be similarly addressed using the results from (i) and (ii).

##### S3.3.1 Orthogonal $u, v$

Again, since the distribution of $(I-J)(zI-C^{-1})^{-1}$ is invariant under any orthogonal similarity transform, its expectation is a scalar matrix $\lambda I$. Using the independence of $J$ to $u, v$,

$$\mathbf{E}_J(u^T(I-J)(zI-C^{-1})^{-1}v) = \lambda \mathbf{E}(u^T v) = 0. \tag{S36}$$

The last equality applies to both orthogonal $u, v$ (can be deterministic) and independent $u, v$ with iid $\mathcal{N}(0, 1/N)$ entries. Together with results in Section S3.3, Eq. (S33) becomes

$$\det \begin{bmatrix} -z\Delta_{C^{-1}}(z) & \frac{1}{x} \\ \frac{1}{x} & -\Delta_{C^{-1}}(z) \end{bmatrix} = 0, \quad \text{or} \quad z\Delta^2_{C^{-1}}(z) = \frac{1}{x^2}. \tag{S37}$$

Clearly, the outliers only depend on $|x|$. As $|x|$ increases from 0, there can be two outliers on each side of the bulk spectrum. Using Eq. (S37), the transition value for the left and right outlier to emerge are, respectively,

$$x_{\min,\pm} = \frac{\sqrt{x_\pm}}{|\Delta_{C^{-1}}(1/x_\pm)|}. \tag{S38}$$

Here $x_\pm$ are edges of $p_C(x)$'s support. The value of $\Delta_{C^{-1}}(1/x_\pm)$ can be calculated using Eq. (S31) with the scaling relation

$$\Delta_{C^{-1}}(z) = \frac{1}{g^2}\Delta_{g^{-2}C^{-1}}\left(\frac{z}{g^2}\right). \tag{S39}$$

Interestingly, we found that $x_{\min,-} < x_{\min,+}$. This means that for $x_{\min,-} < |x| < x_{\min,+}$ there will be only be one outlier to the left of the bulk (Fig. S4A).

To find the location of the outliers. Plugging Eq. (S39) in Eq. (S7) with $\kappa = b = 0, a = 1/g$,

$$z\Delta^2_{C^{-1}} = -\frac{1}{g^2 + \Delta^{-1}_{C^{-1}}} + \frac{1}{(g^2 + \Delta^{-1}_{C^{-1}})^2}. \tag{S40}$$

Let $y = g^2 + \Delta^{-1}_{C^{-1}}(z)$ and use Eq. (S37), we get a quadratic equation for $y$ with two real roots

$$y_\pm = \frac{-x^2 \pm |x|\sqrt{x^2+4}}{2}. \tag{S41}$$

We can then get $\Delta_{C^{-1}} = 1/(y - g^2)$ and subsequently $z$ from Eq. (S37). Finally, the two (potential) outliers of $C$ are $1/z$,

$$w_\pm = \frac{4x^2}{(-x^2 - 2g^2 \pm |x|\sqrt{x^2+4})^2}. \tag{S42}$$

**Stability** Note that Eq. (S42) is valid when $|x| > x_{\min,\pm}$, for $w_\pm$ respectively. In particular, it appears $w_+$ could diverge to infinity when $|x| = \frac{g^2}{\sqrt{1-g^2}}$. But this value is smaller than $x_{\min,+}$ and therefore the divergence is spurious and $w_+$ is finite for any $|x| > x_{\min,+}$. The finite outlier in the covariance is consistent with the stability of the linear network dynamics. For any fixed $g < 1$ and $0 < \epsilon < 1 - g$, Theorem 1.7 in [11] shows that all eigenvalues of $\tilde{J}$ will be close to the circle at the origin with radius $g(1+\epsilon)$ with probability 1 as $N \to \infty$ (here the eigenvalue of the low rank component is zero). This ensures all eigenvalues of $\tilde{J}$ have real parts less than 1 and that the linear system is stable.

We apply the analytical predictions of rank-1 perturbation of $J$ corresponding to diverging motifs (Fig. 5, Fig. S4B). Here $u = (1,1,\ldots,1)^T/\sqrt{N}$, $v = (b_1, b_2, \ldots, b_N)^T/\sqrt{\kappa_{div}N}$, $x = \sqrt{\kappa_{div}}N$ (see Eq. (S22) and Section S3.1). It is easy to see $u^Tv \to 0$ as $N \to \infty$ thus satisfies the orthogonal condition. Another application is the EI network obeying Dale's law in Fig. 6C. We can decompose the connectivity of the EI network as $J = (J - xuv^T) + xuv^T$, where $u = (1,1,\ldots,1)^T/\sqrt{N}$, $v = (-1,-1,\ldots,-1,1,1,\ldots,1)^T/\sqrt{N}$, $x = pw_0N$. For the rank-1 part, $u^Tv = 0$. The first component $J - xuv^T$ has entry distributions that are zero-mean with equal variance, which by itself leads to a covariance with the same spectrum as an iid Gaussian connectivity with matching $g$ (data not shown). Therefore, we can apply the above outlier results based on iid Gaussian connectivity to the EI network (Fig. S4C), which are confirmed by the simulations.

###### S3.3.2 Identical $u = v$

Here Eq. (S36) becomes

$$\begin{aligned}&\mathbf{E}\left(u^T(I-J)(zI-C^{-1})^{-1}u\right) = \mathbf{E}_J(\mathbf{E}_u(u^T(I-J)(zI-C^{-1})^{-1}u)) \\ &= \frac{1}{N}\mathbf{E}_J(\mathrm{tr}((I-J)(zI-C^{-1})^{-1})) =: q.\end{aligned} \tag{S43}$$

Consider

$$F = \frac{1}{N}\log(\det\left(zI - C^{-1}\right)) = \frac{1}{N}\log(\det\left(zI - (I-gJ_0)^T(I-gJ_0)\right)) \tag{S44}$$

as a function of $g$, where $J_0 = J/g$ has iid $\mathcal{N}(0, 1/N)$ entries. We have

$$\begin{aligned}\frac{\partial F}{\partial g} &= \frac{2}{g}\frac{1}{N}\mathrm{tr}((zI-C^{-1})J^T(I-J)) \\ &= -\frac{2}{g}\frac{1}{N}\mathrm{tr}((zI-C^{-1})(I-J)^T(I-J)) + \frac{2}{g}\frac{1}{N}\mathrm{tr}((zI-C^{-1})(I-J)) \\ &= \frac{2}{g}(1 + z\Delta_{C^{-1}}(z) + q)\end{aligned} \tag{S45}$$

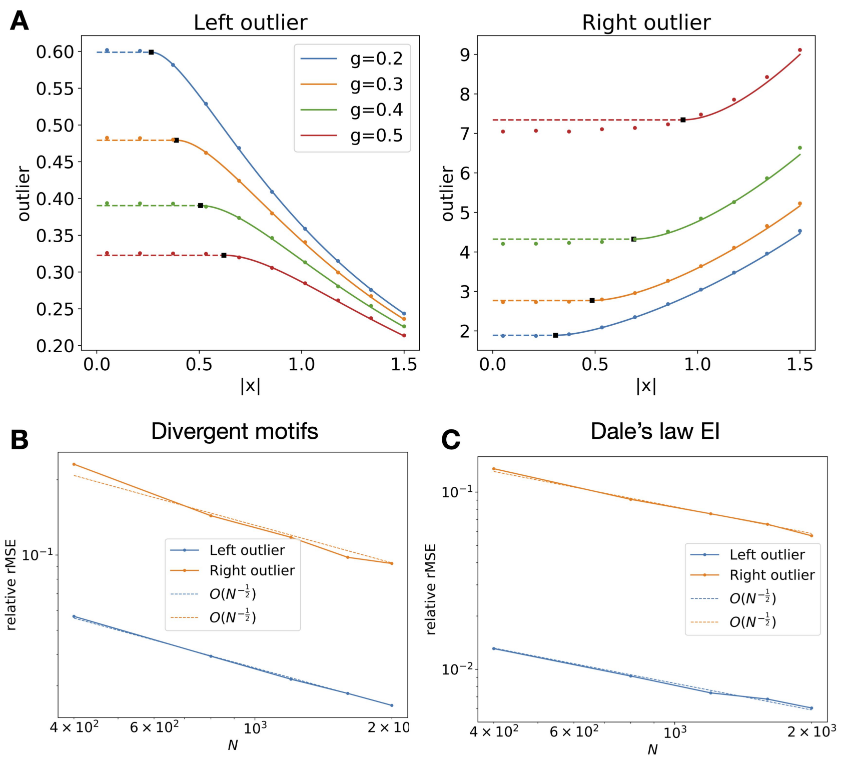

Figure S4: **A**. Similar as Fig. S3A but for the two covariance outliers when perturbing $J$ with a rank-1 matrix with orthogonal column and row vectors (see text). The parameters and line-style meanings are the same as in Fig. S3A. **B**. The root-mean-square error (rMSE) of predicting the two outliers in the random connectivity with diverging motifs (Fig. 5). Here $N = 400$ and $x = 2.066$. For networks with different $N$, we scale $\kappa_{div}$ (Section S3.1) such that $x, g$ and consequently the locations of the outliers remain fixed. The rest of the parameters and lines are the same as in Fig. S3B. For clarity of the plot, the y-axis is the (log scale) rMSE divided by the theoretical prediction of the outlier. **C**. Same as B. but for the Dale's law EI network in Fig. 6B. Again, we fix $x = 8/3$, which corresponds to the parameters in Fig. 6B, by adjusting $p$ properly for different $N$.

We have used Eq. (S35). Note that $F$ is essentially the logarithm potential Eq. (S2),

$$F = \Phi\left(-\frac{z}{g^2}\right) - \log(g^2). \tag{S46}$$

As shown in [10],

$$\Phi(\epsilon) = \text{constant} - \left(-\frac{\epsilon}{\sigma} + \log(\sigma+1) + \frac{g^{-2}}{\sigma+1}\right), \tag{S47}$$

where $\sigma$ is related to $\epsilon, g$ and $\Phi$ by Eq. (S6). In particular, we can replace it with $\Delta_{C^{-1}}(z) = 1/(g^2\sigma)$. After this replacement, we calculate the derivative against $g$ using Eqs. (S46) and (S47),

$$\frac{\partial F}{\partial g} = \frac{2g}{\Delta_{C^{-1}}^{-1} + g^2} - \frac{2g}{(\Delta_{C^{-1}}^{-1} + g^2)^2} = -2gz\Delta_{C^{-1}}^2. \tag{S48}$$

In the calculation, terms containing $\partial_g \Delta_{C^{-1}}$ canceled because of Eq. (S40). This shows

$$\mathbf{E}(u^T(I-J)(zI - C^{-1})^{-1}u) = q = -g^2 z\Delta_{C^{-1}}^2 - z\Delta_{C^{-1}} - 1. \tag{S49}$$

The determinant equation for $z$ Eq. (S33) is then

$$\left(g^2 z\Delta_{C^{-1}}^2 + z\Delta_{C^{-1}} + 1 - \frac{1}{x}\right)^2 = z\Delta_{C^{-1}}^2.$$

Note that the outlier $z$ in $C^{-1}$ is positive and $\Delta_{C^{-1}}$ is real for $z$ outsize of the support of $C^{-1}$. Taking the square root of the above equation,

$$g^2 z\Delta_{C^{-1}}^2 + z\Delta_{C^{-1}} + 1 - \frac{1}{x} = \pm\sqrt{z}\Delta_{C^{-1}} \tag{S50}$$

**Stability** Similar to the case in Section S3.3.1, we can apply the result in [11] to determine stability. Here the rank-1 perturbation has an eigenvalue equals $x$ (as $N \to \infty$). Therefore, for any $g < 1$ the network is stable in the large $N$ limit if and only if $x < 1$.

To proceed with solving the outliers, consider the limiting case of $g \to 0$, there is *one* outlier in $C^{-1}$ at $z = (1-x)^2$. At the same time, $\Delta_{C^{-1}}(z) \to \frac{1}{1-z}$ and we can solve Eq. (S50) explicitly. This confirms that we should take the *negative* sign in Eq. (S50). The outlier for a general $g$ is also qualitatively similar to the case of $g = 0$ (Fig. S5A). For $x < x_{\min,-} < 0$, a unique outlier appears to the left of the bulk of $C$; for $1 > x > x_{\min,+} > 0$, the unique outlier is to the right of the bulk of $C$. The transition value $x_{\min,\pm}$ can be calculated using Eq. (S50) by plugging in $z = x_\pm$ respectively. Here $x_\pm$ are support edges of $p_C(x)$ (Eq. (6)) and the values of $\Delta_{C^{-1}}$ are calculated in the same way as in Section S3.3.1.

Next, we plug in Eq. (S40) to Eq. (S50) to eliminate $z$, and obtain an equation of $\Delta_{C^{-1}}$. This can be solved numerically, for example, using a bisection search. For $x < x_{\min,-} < 0$, we search between $\Delta_{C^{-1}}(z = 1/x_-)$ and $\Delta_{C^{-1}}(z \to \infty) = 0$. For $1 > x > x_{\min,+} > 0$, the search is between $\Delta_{C^{-1}}(z = 0) = (1-g^2)^{-1}$ and $\Delta_{C^{-1}}(1/x_+)$.

Finally, we can check the consistency between the covariance spectrum outlier and the stability of the network by showing the divergence of the outlier as $x \to 1$. For $x > 0$, the outlier in $\tilde{C}$ is to the right of the bulk, hence $z > 0$ is to the left of the pdf of $C^{-1}$. This means $\Delta_{C^{-1}}^{-1}(z) > 0$ (Eq. (S86)). Let $\epsilon := \frac{1}{x} - 1 > 0$, $y := g^2 + \Delta_{C^{-1}}^{-1}(z)$, $\delta := y^2 z\Delta_{C^{-1}}^2 \geq 0$, Eq. (S40) is then $1 - y = \delta$. Multiplying Eq. (S50) (with the negative sign as discussed above) with $y$ and plugging in Eq. (S40), we have

$$-\sqrt{\delta} = \delta - \epsilon y. \tag{S51}$$

Using Eq. (S40), we have $y < 1$. Therefore $\delta - \epsilon \leq \delta - \epsilon y < 0$ and $\delta \to 0$ as $x \to 1$. Under this limit, $y = 1 - \delta \to 1$, $\Delta_{C^{-1}}$ has a positive lower bound, so we must have $z \to 0$ to achieve $\delta = y^2 z\Delta_{C^{-1}}^2 \to 0$. This vanishing limit of $z$ corresponds to an outlier diverging to $+\infty$ in $\tilde{C}$ as expected.

#### S4 Symmetric and anti-symmetric random connectivity

For $\kappa = \pm 1$, $J$ is symmetric and anti-symmetric (or skew-symmetric) respectively and is thus a normal matrix. Therefore, in both cases, we can calculate the eigenvalue distribution of $C$ directly from the spectrum of $J$, which are semicircle laws on either the real or imaginary axis [10]. We derive below the eigenvalue distribution of the frequency dependent covariance (Section S7, Eq. (S81)) which includes the long time window covariance as a special case with $\omega = 0$ (Eqs. (14) and (15)).

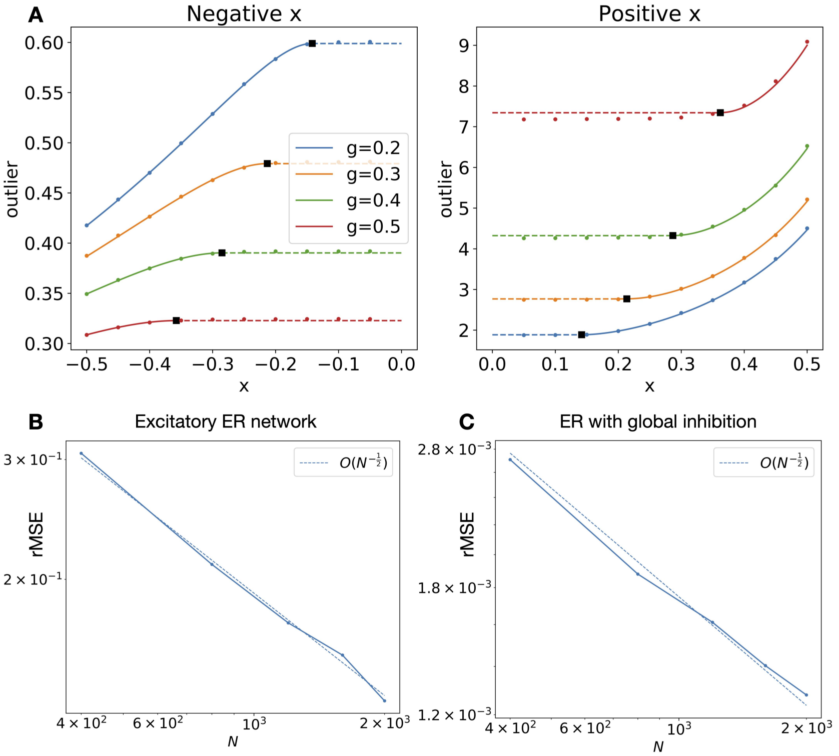

Figure S5: **A**. Similar to Fig. S3A but for the outlier when perturbing $J$ with a symmetric rank-1 matrix (see text). The parameters and line-style meanings are the same as in Fig. S3A. **B**. Similar to Fig. S4 but for the outlier in an all excitatory Erdős-Rényi network. Here we fix $x = 0.57143$ and $g = 0.2$ according to Fig. S8A and adjust $p$ for different $N$. The outlier is to the right of the bulk spectrum. **C**. Same as B. but for the outlier in an Erdős-Rényi network with a global inhibition that dominates. Here we fix $x = -4/3$ and $g = 0.4$ according to Fig. S8B and adjust $p$ for different $N$. The outlier is to the left of the bulk spectrum.

#### S4.1 Random symmetric

For $\kappa = 1$ the symmetric case, as usual we start by calculating the eigenvalue distribution of $P$ (Eq. (S1)) with $\eta = a + b\mathbb{i}$. The result depends on the value of $a$. For $|a| \geq 2$,

$$p_\eta(x) = \frac{\sqrt{4 - a^2 + b^2 - x + 2|a|\sqrt{x - b^2}}}{4\pi\sqrt{x - b^2}}, \quad x \in (b^2 + (|a| - 2)^2, b^2 + (|a| + 2)^2).$$

For $|a| < 2$,

$$p_\eta(x) = \begin{cases} \frac{\sqrt{4-a^2+b^2-x-2|a|\sqrt{x-b^2}}+\sqrt{4-a^2+b^2-x+2|a|\sqrt{x-b^2}}}{4\pi\sqrt{x-b^2}}, & x \in (b^2, b^2 + (|a| - 2)^2) \\ \frac{\sqrt{4-a^2+b^2-x+2|a|\sqrt{x-b^2}}}{4\pi\sqrt{x-b^2}}, & x \in (b^2 + (|a| - 2)^2, b^2 + (|a| + 2)^2) \end{cases}.$$

The stability condition in this case requires $a > 2$, which is what we assume below. For the frequency dependent covariance in Eq.(S81)

$$C = g^{-2}P^{-1}, \quad z = 1/g + \mathbb{i}\omega/g,$$

$$p_C(x) = \frac{\sqrt{(4g^2-1)x - (1-\omega^2 x) + 2\sqrt{x(1-\omega^2 x)}}}{4\pi g^2 x^2 \sqrt{1-\omega^2 x}}, \tag{S52}$$

where the stability condition is $g < \frac{1}{2}$ and the support of the distribution is

$$x \in \left( \frac{1}{\omega^2 + (1+2g)^2}, \frac{1}{\omega^2 + (1-2g)^2} \right). \tag{S53}$$

For $g < 1/2$ and for any frequency, the density at both edges scales as

$$p_C(x) \propto |x - x_0|^{\frac{1}{2}}. \tag{S54}$$

As $g \to 1/2$, the network approaches instability, but this is reflected differently depending on the frequency. At $\omega = 0$, the upper limit of the distribution diverges to infinity as $(1-2g)^{-2}$. The lower edge remains goes to $\frac{1}{4}$ and has the same density scale as Eq. (S54). For non-zero frequency, the upper limit remains finite, but the probability density function at this edge diverges to $+\infty$ (as $(1-\omega^2 x)^{-1/4}$) when $g \to 1/2$. In particular, the shape of the probability density function can become bimodal for large $g$ at non-zero $\omega$.

Let's now look at the tail approximation for $\omega = 0$ as $g \to 1/2$.

$$p_{C,\omega=0}(x) = \frac{\sqrt{(4g^2-1)x - 1 + 2\sqrt{x}}}{4\pi g^2 x^2}.$$

For $x$ far away from the edge of the support, that is,

$$1 \ll x \ll x_+ = (1-2g)^{-2},$$

$(4g^2-1)x = (2g-1)(2g+1)x \ll \sqrt{x}$. Hence the leading-order term of $p_C(x)$ as $g \to 1/2$ is

$$p_C(x) \approx \frac{\sqrt{2}}{\pi} x^{-\frac{7}{4}}. \tag{S55}$$

#### S4.2 Random anti-symmetric

For $\kappa = -1$ the anti-symmetric case, the result of $p_P(x)$ is the same as the symmetric case with real and imaginary axes flipped. Note that the network is stable for any $g$ (this does not follow from flipping the axes).

At high frequencies $|\omega| > 2g$, the density of $p_C(x)$ is smooth.

$$p_C(x) = \frac{\sqrt{(4g^2+1-\omega^2)x - 1 + 2|\omega|\sqrt{x(1-x)}}}{4\pi g^2 x^2 \sqrt{1-x}}, \quad x \in \left( \frac{1}{1+(|\omega|+2g)^2}, \frac{1}{1+(|\omega|-2g)^2} \right) \tag{S56}$$

The density at both of the edges both scales as $|x - x_0|^{\frac{1}{2}}$.

For low frequencies $|\omega| < 2g$,

$$p_C(x) = \begin{cases} \frac{\sqrt{(4g^2+1-\omega^2)x-1+2|\omega|\sqrt{x(1-x)}}}{4\pi g^2 x^2 \sqrt{1-x}}, & x \in \left( \frac{1}{1+(|\omega|+2g)^2}, \frac{1}{1+(|\omega|-2g)^2} \right) \\ \frac{\sqrt{(4g^2+1-\omega^2)x-1-2|\omega|\sqrt{x(1-x)}}+\sqrt{(4g^2+1-\omega^2)x-1+2|\omega|\sqrt{x(1-x)}}}{4\pi g^2 x^2 \sqrt{1-x}}, & x \in \left( \frac{1}{1+(|\omega|-2g)^2}, 1 \right) \end{cases}.$$

The left edges similarly has a density of $|x-x_0|^{\frac{1}{2}}$. The density diverges at the right edge $x=1$ as $(1-x)^{-1/2}$. At $\frac{1}{1+(|\omega|-2g)^2}$ with the support, $p_C(x)$ is continuous but differentiable: its derivative is finite on the left side and is $+\infty$ on the right side ($p_C'(x) \propto |x-x_0|^{-\frac{1}{2}}$).

At the critical frequency (i.e., $|\omega| \to 2g^+$), the density diverges as $(1-x)^{-1/4}$ at the right edge $x_+$. In all cases, the distribution never has a "long tail" as the upper edge of the support is always bounded by 1

$$x_+ = \begin{cases} \frac{1}{1+(|\omega|-2g)^2}, & |\omega| > 2g \\ 1, & |\omega| \le 2g \end{cases} . \tag{S57}$$

#### S5 Random networks with reciprocal motifs: general case

Here we study the eigenvalue distribution of the covariance $C$ when the connectivity is given by a Gaussian random matrix with reciprocal motifs

$$J_{ij} \sim \mathcal{N}(0, g^2/N), \quad \rho(J_{ij}, J_{ji}) = \kappa. \tag{S58}$$

To calculate the spectrum of $C$ (Eq.(2)), we set $\eta = 1/g = a$ and $b = 0$ in Eq. (S7) following the derivations explained in Section S1,

$$z = -\frac{1}{\Delta_\eta(\Delta_\eta+1)} + \frac{\eta^2}{(\Delta_\eta(1+\kappa)+1)^2}. \tag{S59}$$

This in general is a quartic equation for $\Delta_\eta$,

$$\begin{aligned}(\kappa+1)^2\Delta_\eta^4 + (\kappa+1)(\kappa+3)\Delta_\eta^3 + \frac{z(2\kappa+3)-\eta^2+(\kappa+1)^2}{z}\Delta_\eta^2 \\ + \frac{z-\eta^2+2(\kappa+1)}{z}\Delta_\eta + \frac{1}{z} = 0,\end{aligned} \tag{S60}$$

and the roots can be solved using Ferrari's formula. Note we will select the pair of complex roots that is associated with the branch that approaches $-\frac{1}{z}$ as $z \to \infty$.

The branching points of when a double root occurs, contain the edges of the support of $p_P(x)$ and can be found analytically by solving for $f'(\Delta_{\eta,0}) = 0$, and $z = f(\Delta_{\eta,0})$, where $f(\Delta_\eta)$ is the right hand side of Eq. (S59)

$$f'(\Delta_\eta) = \frac{1}{\Delta_\eta^2} - \frac{1}{(\Delta_\eta+1)^2} - \frac{2\eta^2(1+\kappa)}{(\Delta_\eta(1+\kappa)+1)^3}. \tag{S61}$$

The above leads to a quartic equation in $\Delta_\eta$. We explain below the behavior of the roots under the stability condition that $g(1+\kappa) < 1$. For $-1 < \kappa \le 0$, Eq. (S61) has two real roots and they correspond to the support of $p_P(x)$. For $0 < \kappa < 1$, Eq. (S61) has four real roots, corresponding to four branch points. In such cases, the corresponding $\mathrm{Im}(\Delta_\eta)$ are two finitely supported continuous functions associated with two pairs of complex roots of Eq. (S60), with edges matching the branching points. However, only one of the functions corresponds to $p_P(x)$, moreover, the support edges of the two functions can interlace. We can select the correct density function/complex roots, based on continuity when we vary $\kappa$ from 0 which is the iid Gaussian case that we already know $p_P(x)$ (Eq. (S11)). In this process of increasing $\kappa$ from 0, one function appears at $+\infty$ and moves towards 0 and coincides with the other function when $\kappa = 1$. This shows that the function/root branch near 0 is the one that corresponds to $p_P(x)$.

To find the dimension $D$ for the covariance eigenvalues, we will derive an equation based on Eq. (S59) for the moment generating function

$$M_A(z) = \sum_{n=1}^{\infty} \mu_n z^n, \tag{S62}$$

where $\mu_n = \int x^n p_A(x) dx$. $M(z)$ is related to the Stieltjes transform by

$$M_A(z) = -\Delta_A\left(\frac{1}{z}\right)\frac{1}{z} - 1. \tag{S63}$$

For reciprocal variables $y = 1/x$ with distributions on $\mathbb{R}^+$, their Stieltjes transforms are related by

$$-\frac{1}{z^2}\Delta_x\left(\frac{1}{z}\right) - \frac{1}{z} = \Delta_y(z). \tag{S64}$$

Using Eqs. (S59), (S63) and (S64) we get

$$\frac{1}{z} = -\frac{1}{M(M+z)} + \frac{\eta^2}{(M(1+\kappa)+z)^2}, \tag{S65}$$

where $M(z)$ is the moment generating function for the eigenvalues of matrix $P$ (Eq. (S1)).

We can expand Eq. (S65) in series of $z$ and matching the coefficients of lower-order terms to calculate the first two moments $\mu_1, \mu_2$. The lowest order term after multiplying both sides by the denominators $M(M+z)(M(1+\kappa)+z)^2$ is $z^3$. This leads to a quadratic equation for $\mu_1$,

$$\mu_1^2(\eta^2 - \beta^2) - 2\mu_1(\beta - \frac{\eta^2}{2}) - 1 = 0.$$

Here $\beta = 1+\kappa < \eta = 1/g$. The equation has two real roots one of them is positive, which corresponds to the moment of eigenvalues, so

$$\mu_1 = \frac{2\beta - \eta^2 + \eta\sqrt{\eta^2 - 4\beta + 4}}{2(\eta^2 - \beta^2)}, \quad \text{or } \mu_1(C) = \frac{2(1+\kappa)g^2 - 1 + \sqrt{1 - 4g^2\kappa}}{2(1 - g^2(1+\kappa)^2)g^2}. \tag{S66}$$

Note that the "$\mu_1$" in the main text is the $\mu_1(C)$ here. Matching the coefficient at the next order, $z^4$, leads to a linear equation of $\mu_2$,

$$\mu_2 = \frac{(\mu_1\beta + 1)^2(\mu_1 + 1)\mu_1}{\eta^2(2\mu_1 + 1) - 2\beta(\mu_1\beta + 1)}, \quad \mu_2(C) = \mu_2/g^4.$$

From these moments, the dimension is

$$D = N\frac{\mu_1(C)^2}{\mu_2(C)} = N\frac{(\eta^2(2\mu_1 + 1) - 2\beta(\mu_1\beta + 1))\mu_1}{(\mu_1\beta + 1)^2(\mu_1 + 1)}.$$

These are the same as Eq. (18).

For the special case of symmetric ($\kappa = 1$) and anti-symmetric ($\kappa = -1$) random connectivity,

$$\mu_1(C, \kappa = 1) = \frac{2}{(1 + \sqrt{1 - 4g^2})\sqrt{1 - 4g^2}}, \tag{S67}$$

$$\mu_1(C, \kappa = -1) = \frac{2}{1 + \sqrt{1 + 4g^2}}, \tag{S68}$$

$$D(\kappa = 1) = N\frac{(1 - 4g^2)^{\frac{3}{2}}(1 - \sqrt{1 - 4g^2})^2}{4g^4}, \tag{S69}$$

$$D(\kappa = -1) = N\frac{\sqrt{1 + 4g^2}(\sqrt{1 + 4g^2} - 1)^2}{4g^4}. \tag{S70}$$

#### S5.1 Power-law approximation

We use the alternative approach described in Section S1.2 to derive a power-law approximation for $p_C(x)$ with reciprocal motifs as $g$ approaches the critical $g_c = 1/(1+\kappa)$ and for large $x \to \infty$. Here we consider the general non-degenerate case of $-1 < \kappa < 1$, since the special cases of $\kappa = \pm 1$ are already studied in Sections S4.1 and S4.2.

Starting from Eq. (S60) and setting $a = 1/g_c = 1 + \kappa$ (no singularity for the distribution of $P$) and $z = x \in \mathbb{R}^+$.

$$xa^2\Delta_\eta^4 + xa(a+2)\Delta_\eta^3 + x(2a+1)\Delta_\eta^2 + (x + a(2-a))\Delta_\eta + 1 = 0 \tag{S71}$$

We will study the leading-order approximation of the roots to this quartic equation as $z = x \to 0^+$, because this will translate to the large $x$ approximation in $p_C(x)$. Following methods in perturbation theory of polynomial equations, we enumerate the possible order (in terms of $x$) of the roots by assuming each term in Eq. (S71) as being the leading term as $x \to 0^+$. The first possibility is when $a(2-a)\Delta_\eta$ is leading and this corresponds to a real root that approaches $-\frac{1}{a(2-a)}$ ($O(1)$). This does not correspond to $p_P(x)$.

The remaining three roots have magnitude $|\Delta_\eta| \to \infty$ and to the leading order, Eq. (S71) becomes

$$xa^2\Delta_\eta^4 + a(2-a)\Delta_\eta = 0 \;\text{ or }\; xa^2\Delta_\eta^3 + a(2-a) = 0.$$

Here two complex roots correspond to $p_P(x)$, so we have

$$\lim_{x\to 0^+} p_{P,g=g_c}(x) / \left( \frac{\sqrt{3}}{2\pi} x^{-\frac{1}{3}} \left( \frac{1-\kappa}{1+\kappa} \right)^{\frac{1}{3}} \right) = 1. \tag{S72}$$

Using relation Eq. (S1), this means

$$\lim_{x\to\infty} \lim_{g\to g_c} p_C(x) / \left( \frac{\sqrt{3}}{2\pi} x^{-\frac{5}{3}} (1-\kappa)^{\frac{1}{3}} (1+\kappa) \right) = 1. \tag{S73}$$

Importantly, this has the same power-law exponent as the iid Gaussian case Eq. (7).

#### S5.2 Strong non-normal effects at $\kappa < 0$

As stated in Section 3.3.2, unlike the extreme (and normal $J$) case of $\kappa = -1$ (Fig. 3B), for all $\kappa > -1$ and $g < g_c$, there is no divergence in $p_{C,g,\kappa}(x)$ with the asymmetric random connectivity $J$, and the pdf is unimodal for all but a minuscule set of combinations of $\kappa$ and $g$ where $\kappa$ is very close to $-1$ (Fig. S6B-E). Note that the main plot in Fig. S6E is a highly zoomed-in view to show the bimodal region, which is hard to see within the whole attainable parameter region (the inset of panel E).

This is in sharp contrast to the case where $J$ is normal. To show this, we compare $p_{C,g,\kappa}(x)$ to the covariance eigenvalue distribution $p_{C^n,g,\kappa}(x)$ from a network with a *normal* connectivity matrix $J^n$ that has a matching eigenvalue distribution to $J$ (Section S5.3). Note that at $\kappa = -1$, $J$ itself is normal and $p_{C,g,\kappa=-1}(x) = p_{C^n,g,\kappa=-1}(x)$. Interestingly, the normal matching $p_{C^n,g,\kappa}(x)$ shows similar features as the $\kappa = -1$ (Fig. 3B and the magenta line in Fig. S6B). First, the diverging density of $p_{C,g,\kappa=-1}(x)$ at the upper edge $x = 1$ becomes a non-differentiable peak in $p_{C^n,g,\kappa}(x)$ when $g$ is larger than the value shown by the black dashed curve in Fig. S6F (derived from the tangent condition above). This non-smooth peak continues to exist throughout $g < g_c$, so there is no second transition as in the non-normal connectivity case (the magenta curve in Fig. S6E). Second, the distribution is bimodal in a much larger region of the parameter space $(g, \kappa)$, compared to the case of $p_C(x)$ (Fig. S6F vs Fig. S6E-inset ).

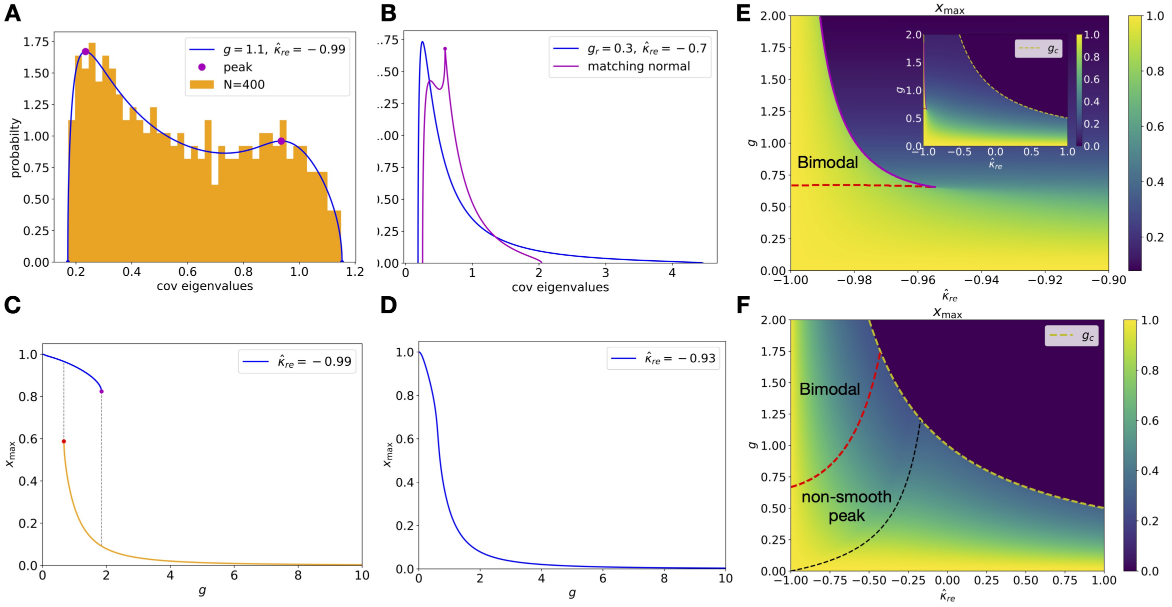

Figure S6: **Strong non-normal effect at** $\hat{\kappa}_{re} < 0$. **A**. An example bimodal distribution at $\hat{\kappa}_{re} = -0.99$ compared with $N = 400$ simulation (a single realization). **B**. A typical unimodal spectrum (blue) for $\hat{\kappa}_{re} < 0$ compared with the covariance spectrum predicted from the spectrum of connectivity $J$ assuming it is a normal matrix (magenta). The normal prediction is very different and bears similarity to the anti-symmetric case (Fig. 3B) by being bimodal with a non-differentiable point (dot). **C**. Location of the peak(s) in the covariance spectrum as a function of $g$ at $\hat{\kappa}_{re} = -0.99$. As $g$ increases, a left peak emerges at $g = \frac{2}{3}$, and the right peak disappears after $g \approx 1.85$. For a range of $g$ between the bifurcations, the distribution is bimodal. **D**. Same as C. but for $\hat{\kappa}_{re} = -0.93$. There is no bifurcations of peaks and the distribution is always unimodal. **E**. The location of the peak (rightmost if multiple) as a function of $g$ and $\hat{\kappa}_{re}$. The red dashed line and magenta solid lines label the location of the peak bifurcations as in C. (same colored dots) and the triangular region enclosed is when the covariance spectrum is bimodal. Note the $\hat{\kappa}_{re}$-axis is highly enlarged and the bimodal region is minuscule within the whole parameter space only for $\hat{\kappa}_{re} \lesssim -0.95$ (inset, the yellow dashed line shows the stability bound $g_c$). **F**. Same as E. but for the normal matching spectrum $p_{C^n}(x)$. The bimodal region is enclosed between the red (left peak emerges) and yellow ($g_c$) dashed lines. The region is much larger compared to the plot for the actual covariance spectrum (E-inset), where the bimodal region is almost invisible under the same axes scales.

#### S5.3 Matching normal connectivity

Here we consider the spectrum of a covariance $C^n$ with connectivity $J^n$ that is a normal matrix and has the same eigenvalue distribution as the Gaussian random connectivity $J$. With reciprocal motifs (Eq. (S58)), the eigenvalues of $J$ in the large $N$ limit is uniformly distributed in an ellipse on the complex plane [10], centered at the origin with horizontal and vertical half-axes being $g(1 \pm \kappa)$, respectively. Since $J^n$ is a normal matrix, the eigenvalues of its covariance matrix $C^n$ (defined via Eq. (2) with $J^n$) are directly related to those of $J_n$ (and $J$), that is,

$$\lambda(C^n) = |1 - \lambda(J^n)|^{-2} = |1 - \lambda(J)|^{-2}.$$

Consider a circle centered at $(1, 0)$ with radius $1/x^2$. The probability $p_{C^n}(x)dx$ is then proportional to the arc length of the part of the circle inside the ellipse (Fig. S7) where $\lambda(J)$ is distributed in

$$p_{C^n}(x)dx = \frac{1}{\pi ab}|\Omega(\theta)|rdr, \quad x = 1/r^2, \ a = g(1 + \kappa), \ b = g(1 - \kappa) \tag{S74}$$

Here $\Omega(\theta)$ is the range of angle of the circle that is inside the ellipse, which can be one single or two intervals whose endpoints can be easily solved. Given the endpoints, $|\Omega(\theta)|$ is the length of the angle interval(s) (see Fig. S7) and $p_{C^n}(x)dx$ is derived by a change of variable.

When $b^2 - a^2 - a > 0$ (thus $\kappa$ must be negative), the largest $r$ having an intersection with the ellipse occurs when the circle is tangential to the ellipse (Fig. S7),

$$r_t = b\sqrt{\frac{b^2 - a^2 + 1}{b^2 - a^2}}.$$

In this case the support of $p_{C^n}(x)$ is $[r_t^{-2}, (1 - g(1+\kappa))^{-2}]$, otherwise the support is $[(1 + g(1 + \kappa))^{-2}], (1 - g(1+\kappa))^{-2}]$.

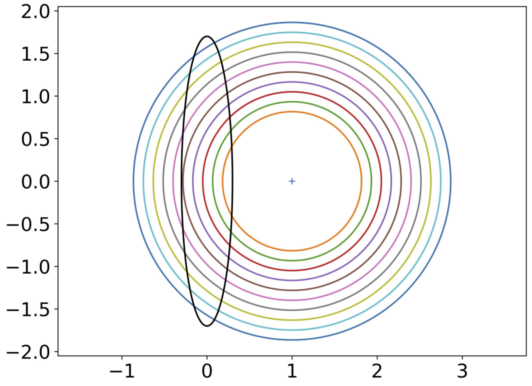

Figure S7: Deriving the covariance spectrum pdf for a normal $J$ with matching eigenvalue distribution (i.e., the ellipse centered at the origin) based on the arc length. See text for details.

For the case of iid $J$ with $\kappa = 0$, the eigenvalue distribution of normal matching covariance $C^n$ is

$$p_{C^n}(x) = \frac{1}{\pi g^2 x^2} \arccos\left(\frac{(1-g^2)x^{\frac{1}{2}} + x^{-\frac{1}{2}}}{2}\right) dx, \quad \frac{1}{(1+g)^2} \le x \le \frac{1}{(1-g)^2}. \tag{S75}$$

Using this formula, we can make quantitative comparisons to the iid Gaussian random connectivity case (Eq. (5)). First, the density at both edges has the same scaling of $|\Delta_\eta x|^{\frac{1}{2}}$ (Section S1.1). This can be shown by the Taylor expansion near the edges. Let $f(x) = \frac{1}{2}((1-g^2)x^{\frac{1}{2}} + x^{-\frac{1}{2}})$. Near the edges, $f(x) \to 1$,

$$\arccos(f(x)) = \arcsin(\sqrt{1 - f^2(x)}) = \sqrt{1 - f(x)}\sqrt{1 + f(x)} + o(\sqrt{1 - f^2(x)}). \tag{S76}$$

The Taylor expansion near the edges is

$$1 - f(x) \approx -f'(x)dx. \tag{S77}$$

The derivative at the edges

$$f'(x) = \frac{1}{4}x^{-\frac{1}{2}}(1 - g^2 - x^{-1}) = -\frac{1}{2}(1 \pm g)^2 g.$$

Thus for $g < 1$, $f'(x)$ is non-zero and finite, thus the approximation in Eq. (S77) indeed gives the leading term approximation near the edges. Plugging this into Eq. (S76) shows the densities at the two edges scale as $|x - x_0|^{\frac{1}{2}}$.

As $g \to 1^-$, the right edge of $p_{C^n}(x)$ diverges slower as $O((1-g)^{-2})$ comparing to $O((1-g)^{-3})$ of $p_C(x)$; and the tail is approximated by a faster decaying power law

$$p_{C^n}(x) \approx \frac{1}{2}x^{-2}. \tag{S78}$$

### S6 Sparse connectivity and Excitatory–Inhibitory networks

Here we include results of several additional sparse random (i.e., Erdős-Rényi (ER)) networks in addition to the cases in the main text and Fig. 6.

**All-excitatory network** Here each pair of neurons is independently decided to be connected with a weight $w0 > 0$ at probability $0 < p < 1$ (and unconnected with probability $1 - p$). Similar to the EI network discussed in the main text, the covariance matrix spectrum from the iid Gaussian connectivity (Eq. (5)) also describes the bulk spectrum of an ER network once substituting $g = w_0\sqrt{Np(1-p)}$ to match the variance of connections $\mathrm{var}(J_{ij})$ (Fig. S8A). The non-zero mean connections which is a rank-1 matrix generates an outlier to the right of the bulk covariance spectrum (Fig. S8A) and its location is derived in Section S3.3.2. Note this right outlier can affect the stability of the network. In particular, the network becomes unstable and the outlier goes to infinity as $w_0pN \to 1$ (Section S3.3.2).

**Adding all-to-all inhibition** The previous case means either $w_0$ or $p$ has to be small to ensure stability. Such a limitation can be overcome by adding inhibition [8] as discussed below. One option is to add a *global inhibition*, where the summed output of all neurons is fed back to each neuron with a strength of $-w_I < 0$. This is equivalent to having a connectivity $J - w_I\mathbf{1}$. Since this is again a rank-1 perturbation, this ER network with inhibition has the same bulk spectrum described by Eq. (5) (Fig. S8BC). The added inhibition precisely reduces the right outlier described above, thus allowing for a larger effective $g$ while keeping the network stable. In particular, when $w_I = w_0p$ there is no outliers in the covariance spectrum (Fig. S8C).

**All inhibitory network** In the ER network, if we have all non-zero connections being $-w_0 < 0$, then the non-zero mean will not cause instability. In this case, there is an outlier to the left of the bulk spectrum (Fig. S8D) similar to the case where we add a large all-to-all inhibition (inhibitory dominant $-w_i < w_0p$, Fig. S8B).

**Mixed sparse E and I connections** Another E-I model is a network with

$$J_{ij} = \begin{cases} w_e, & \text{with prob. } p_e \\ -w_i, & \text{with prob. } p_i \\ 0, & \text{with prob. } 1 - p_e - p_i \end{cases} \tag{S79}$$

where $w_e, w_i > 0$ and satisfies the balance $w_ep_e - w_ip_i = 0$ to ensure a zero mean. Note that since $J_{ij}$ is chosen independently, this network does not obey Dale's law. Here there is no outlier (no low rank structure due to non-zero mean), and the covariance spectrum of $J$ is again described by Eq. (5) (Fig. S8E).

**Unequal fractions of E and I neurons** The key for applying the theory based on the Gaussian random connectivity to the Dale's law EI network is that the variance of connections $\mathrm{var}(J_{ij})$ is (for large $N$) the same regardless of the cell type of $i, j$. This allows us to generalize to have unequal fractions of E and I neurons (Fig. S8F), as long as the variance is kept equal by adjusting the connection probability $p_e$, $p_i$ and individual non-zero connection weights $w_e$, $w_i$

between cell types. Here for simplicity we assume the probability and weights of connections only depend on the cell type of the sending neuron (a rank-1 perturbation due to the mean).

In summary, for all sparse random networks discussed here and in the main text, the bulk spectrum is well described by the iid Gaussian random connectivity theory Eq. (5) with an effective $g$ matching the variance of connection strength.

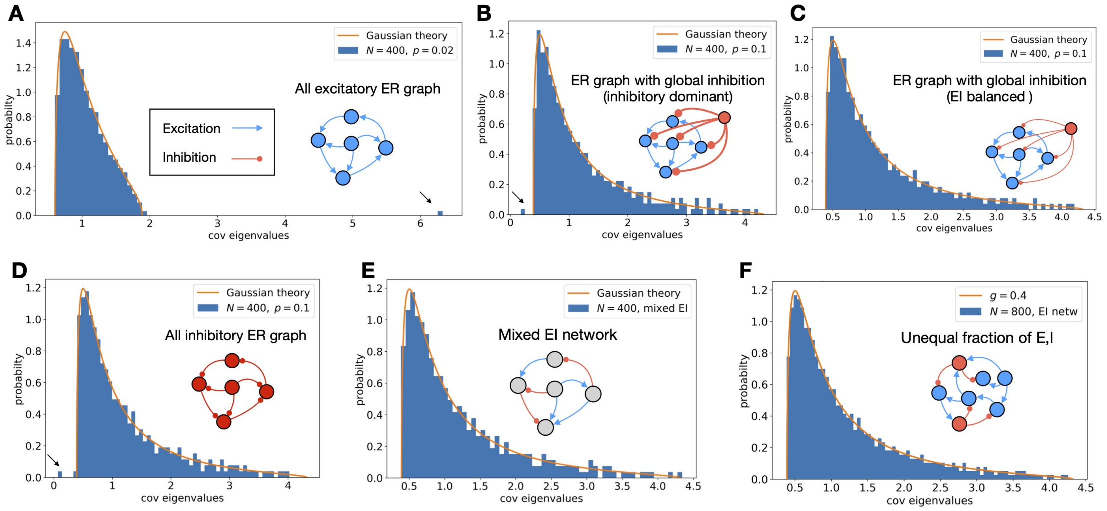

Figure S8: **Additional Erdős-Rényi and EI networks**. **A**. An all-excitatory Erdős-Rényi network with connection probability $p = 0.02$. There is a large outlier caused by the positive mean connection (arrow). The connection strength $w_0 = 0.0714$ is chosen to set the effective $g = 0.2$ and while keeping the network stable. $N = 400$. **B**. An ER network with global inhibition but is inhibitory dominant (negative average connection strength). $N = 400$, $p = 0.1$, effective $g = 0.4$. $w_i$ is 1.5 times the value to balance excitation. This resulted in a small outlying eigenvalue to the left of the bulk (arrow). The bulk spectrum is still closely described by the Gaussian random connectivity theory (curve). **C**. Similar to B. with a global inhibition that balances the mean connection to 0 and eliminates the outlier (see text). $N = 400$, connection probability $p = 1$, the effective $g = 0.4$. **D**. An all inhibitory ER network that has a small outlier to the left of the bulk (arrow). $N = 400$, $p = 0.1$, effective $g = 0.4$. **E**. A mixed EI network with randomly assigned E or I edges (Eq. (S79)). $p_e = 0.025$, $p_i = 0.075$, $N = 400$, the effective $g = 0.4$. **F**. The *bulk spectrum* (excluding two outlying eigenvalues similar to Fig. 6B) for a Dale's law EI network with 70% E and 30% I neurons ($N = 800$). The connection probabilities and weights are chosen to ensure the variance of E and I connection strengths are equal: $p_e = 0.028$, $p_i = 0.15$, $w_e = 0.0864$, $w_i = 0.0396$. The effective $g = 0.4$.

#### S7 Frequency dependent covariance

Correlations beyond the long time window covariance can be described by the frequency dependent covariance $C(\omega)$ (Eq. (19)). Note that by definition,

$$C(\omega)^\dagger = C(\omega), \quad C(\omega)^T = \overline{C(\omega)} = C(\bar{\omega}),$$

and $C(\omega)$ is positive definite ($X^\dagger$ is the conjugate transpose of a matrix $X$). These also ensure the eigenvalues of $C(\omega)$ are always real and nonnegative so the spectrum $p_{C(\omega)}(x)$ is still a distribution on $\mathbb{R}^+$. The time-lagged matrix $C_{ij}(\tau) = \langle \Delta y_i(t) \Delta y_j(t - \tau) \rangle$ is related to $C(\omega)$ by the Fourier transform (Wiener-Khinchin theorem)

$$C(\omega) = \int_{-\infty}^{\infty} e^{-\mathbb{i}\omega\tau} C(\tau) d\tau. \tag{S80}$$

In particular, this shows $C(\omega = 0)$ is the long time window covariance (Eq. (2)).

For the iid Gaussian random connectivity, the spectrum of $C(\omega)$ can be obtained by replacing $g$ with $g(\omega)$ (Eq. (21)). This is because the Stieltjes transform of $P$ (Eq. (S8)) only depends on $|\eta| = 1/g(\omega)$. For other connectivity in general, the shape of the eigenvalue distribution of $C(\omega)$ depends on the phase of $\omega$ in addition to its norm. To illustrate with a concrete example, consider $C\omega)$ in Eq. (20) with $\tau = 1$ and $\sigma^2 = 1$,

$$C(\omega) \propto g^{-2}(\eta I - J^0)^{-1}(\eta I - J^0)^{\dagger -1}, \quad \eta = (1 + \mathrm{i}\omega)/g. \tag{S81}$$

Here $J^0 = J/g$, $\mathrm{var}(J^0_{ij}) = 1/N$, and we omitted a factor of $(1 + \omega^2)$ for simplicity. In Section S4, we calculate the eigenvalue distribution of Eq. (S81) for symmetric and anti-symmetric random connectivity and show that the distribution shape does not only depend on $|\eta|$.

### S8 Time-sampled covariance matrix

Here we give an alternative derivation of the key relation Eq. (22) which is a result from [2]. We later use a similar approach for the space-sampled case. Following [2], we assume the neural activity data follows a zero-mean Gaussian distribution. The sample covariance matrix can then be written as

$$\hat{C} = \frac{1}{M} X X^T$$

where the columns of $X$, $X_{\cdot,t}$, are independent (across $t$) and are drawn from the zero-mean Gaussian distribution with covariance $C$. We can rewrite $\hat{C}$ equivalently (in distribution) as

$$\hat{C} = (I - J)^{-1} Z Z^T (I - J)^{-T}.$$

where $Z_{ij} \sim \mathcal{N}(0, 1/M)$ are iid Gaussian variables which can also be interpreted as the white noises received by individual neurons. Note that this matrix has the same non-zero eigenvalues as

$$(I - gJ)^{-T}(I - gJ)^{-1} Z Z^T =: \check{C}(Z Z^T), \tag{S82}$$

where $\check{C} = (I - gJ)^{-T}(I - gJ)^{-1}$. Importantly, Eq. (S82) is a *free product* of $\check{C}$ and $ZZ^T$. This can be shown by checking a sufficient condition in [9]: (i) $\check{C}$ and $ZZ^T$ are independent and (ii) the distribution of $ZZ^T$ is invariant under any orthogonal similarity transformation. Condition (i) is true since $J$ and $Z$ are independent. Condition (ii) is satisfied because for any $UU^T = I$, $U(ZZ^T)U^T = (UZ)(UZ)^T$ and the columns of $UZ$ are still independent with iid $\mathcal{N}(0, 1/M)$ entries. Note that the eigenvalues of $\check{C}$ are the same as $C$, so we can substitute $\check{C}$ with $C$ in the free product for the purpose of studying the eigenvalue distribution.

#### S8.1 Dimension

For a free product, the moments $\mu_n$ of the eigenvalue distribution can be calculated from its factors using decomposition rules from free probability theory [9] expressed using *free cumulants* $\kappa_n$. The $\kappa_n$ and $\mu_n$ are related by enumerating non-crossing partitions (which is the same as the usual cumulant/moment relation for random variables up to the third order). For a free product,

$$\mu_1(\hat{C}) = \mu_1(C)\mu_1(ZZ^T) = \mu_1(C),$$

$$\begin{aligned}
\mu_2(\hat{C}^2) &= \phi(\hat{C}^2) = \phi(C(ZZ^T)C(ZZ^T)) \\
&= \kappa_2(C)\kappa_1(ZZ^T)^2 + \kappa_2(ZZ^T)\kappa_1(C)^2 + \kappa_1(C)^2\kappa_1(ZZ^T)^2, \\
&= \kappa_2(C) + (1+\alpha)\kappa_1(C)^2 = \mu_2(C) + \alpha\mu_1(C)^2.
\end{aligned}$$

Here $\phi(\cdot) = \frac{1}{N}\mathrm{tr}(\cdot)$ and by definition of the free cumulant, $\kappa_1(A) = \mu_1(A)$, $\kappa_2(A) = \mu_2(A) - \mu_1(A)^2$. We have used the properties of the Marchenko–Pastur law: $\kappa_1(M) = 1$, $\kappa_2(M) = \alpha$. The (relative) dimension of $\hat{C}$ (Eq. (23)) then follows immediately from these relations on first two moments

$$\hat{D}(\hat{C}) = \frac{\hat{D}(C)}{1 + \alpha\hat{D}(C)}, \quad D(\hat{C}) = D(C)\frac{N}{N + \alpha D(C)}.$$

For $\alpha = N/M > 1$ ($M$ is the number of samples), there are $N - M$ trivial zero eigenvalues in $\hat{C}$. We can remove them and calculate the dimension of the non-zero eigenvalues. This does not change $D$ but the relative $\hat{D}(\hat{C})$ is now $D/M = \alpha D/N$. Therefore,

$$\hat{D}(\hat{C}) = \frac{\hat{D}(C)}{\frac{1}{\alpha} + \hat{D}(C)}, \quad \alpha > 1. \tag{S83}$$

This shows that $\hat{D}$ increases with $\alpha > 1$.

##### S8.2 Equation relating the sampled and exact spectrum

Beyond the dimension, it is possible to extend the relation to determine the spectrum of $\hat{C}$ [2]. For a free product, the limiting eigenvalue distribution can be determined from those distributions of the factors using a generating function called S-transform [12]

$$S_{\hat{C}}(z) = S_{\check{C}}(z) S_{ZZ^T}(z). \tag{S84}$$

The S-transform of the spectrum of matrix $A$ is defined by

$$S_A(z) = \frac{1+z}{z} M_A^{(-1)}(z), \tag{S85}$$

Here $M_A(z)$ is the moment generating function (Eq. (S62)) and $M_A^{(-1)}(z)$ denotes the inverse function. Using the relation between $M_A(z)$ and the Stieltjes transform (Eq. (S63)), we have

$$-zS(z) = \Delta\left(\frac{1+z}{zS(z)}\right). \tag{S86}$$

The eigenvalue distribution of $ZZ^T$ follows the Marchenko–Pastur law

$$p_{MP}(x) = \frac{\sqrt{(\alpha_+ - x)(x - \alpha_-)}}{2\pi\alpha x}, \quad \alpha_\pm = (1 \pm \sqrt{\alpha})^2, \quad 0 < \alpha = N/M < 1. \tag{S87}$$

For $\alpha > 1$, there is an additional delta distribution at 0 with probability $1 - 1/\alpha$. The eigenvalues of $\check{C}$ are the same as $C$, so the distribution is given by the spectrum of the exact covariance. This points to a general strategy to calculate the eigenvalue distribution of the sample covariance:

$$\begin{aligned} &p_C(x) \to \Delta_C(z) \to S_C(z) \to S_{\hat{C}}(z) = S_C(z) S_{MP}(z) \\ &\to \Delta_{\hat{C}}(z) \to p_{\hat{C}}(x). \end{aligned} \tag{S88}$$

In particular, this allows us to re-derive the same result of [2] (Eq. (22)). First, we need to derive $S_{MP}(z)$. As a standard result (for example see [9]), the free cumulants of the Marchenko–Pastur law is $\kappa_n = \alpha^{n-1}$. Then its R-transform $R(z) = \sum_{n=1}^{\infty} \kappa_n z^{n-1}$ is simply $R_{MP}(z) = \frac{1}{1-\alpha z}$. The R transform is related to the Stieltjes transform by [9]

$$\frac{1}{\Delta(z)} + z = R(-\Delta(z)). \tag{S89}$$

Using Eqs. (S86) and (S89) and $R_{MP}(z)$ we find the S-transform for Marchenko–Pastur law is

$$S_{MP}(z) = \frac{1}{1+\alpha z}. \tag{S90}$$

Note that the generating function $W_A(z) = M_A(1/z)$ and from Eq. (S85)

$$S(z) = \frac{z+1}{z} \frac{1}{W^{(-1)}(z)}. \tag{S91}$$

Plugging this equation in the free product identity (Eq. (S84)) and using the expression for $S_{MP}(z)$, we have

$$\hat{W}^{(-1)}(z) = W^{(-1)}(z)(1+\alpha z), \tag{S92}$$

where $\hat{W}(z)$ is the corresponding generating function for $\hat{C}$. Finally, substitute $z$ with $W(z)$ and apply $\hat{W}(\cdot)$ on both sides,

$$\hat{W}(z \cdot (1+\lambda W(z))) = W(z).$$

If we instead substitute $z$ with $\hat{W}(z)$ and apply $W(\cdot)$ on both sides after dividing by $(1+\alpha\hat{W}(z))$, we get the other identity of Eq. (22).

For $\alpha > 1$, we remove the $N - M$ zero eigenvalues. The Stieltjes transform and $W(z)$ after the removal $\Delta_{\hat{C}_+}$ and $W\hat{C}_+$ satisfy

$$\Delta_{\hat{C}_+} = \alpha\Delta_{\hat{C}} + (\alpha-1)\frac{1}{z}, \quad W_{\hat{C}_+}(z) = \alpha W_{\hat{C}}(z). \tag{S93}$$

So we can simply replace $\hat{W}(z)$ with $\hat{W}(z)/\alpha$ in Eq. (22) when $\alpha > 1$.

#### S8.3 Time-sampled covariance spectrum for iid Gaussian connectivity

Here we use the general relation Eq. (22) to derive the time-sampled covariance spectrum when the connectivity is iid Gaussian. Using $W(z) = -\Delta(z)z - 1$ and Eq. (S64) in Eq. (S8), we have the following equation for $W(z)$ of the matrix $(\eta I - J)^{-1}(\eta I - J)^{\dagger -1}$,

$$z^2W^3 + 2zW^2 + (1 + z(1 - |\eta|^2))W + 1 = 0. \tag{S94}$$

Using the relation with its sampled counterpart $\hat{W}$ (Eq. (22)),

$$(z+\alpha)^2\hat{W}^3 + (2z + 2\alpha + z\alpha(1-|\eta|^2) + \alpha^2)\hat{W}^2 + (1 + z(1-|\eta|^2) + 2\alpha)\hat{W} + 1 = 0. \tag{S95}$$

We can solve $\hat{W}$ from this cubic equation using the Cardano formula for $z = x \in \mathbb{R}$, $\eta = 1/g$. Then the corresponding $\hat{\Delta}(z) = -(1+\hat{W}(z))/z$ is the Stieltjes transform of the eigenvalue distribution of $g^2\hat{C}$, and $p_{\hat{C}}(x)$ is obtained by a simple rescaling $p_{\hat{C}}(x) = g^2 p_{g^2\hat{C}}(g^2x)$. For $\alpha > 1$, according to Eq. (S93), we only need to additionally multiply the above solution by $\alpha$: $p_{\hat{C}_+}(x) = \alpha p_{\hat{C}}(x)$. We see that for $\alpha > 1$, the time-sampled spectrum shifts rightwards when increasing $\alpha$ (Fig. S9A).

The support of $p_{\hat{C}}(x)$ can be found by considering when the determinant of the cubic equation Eq. (S95) turns zero (then scale by $g^2$), because the imaginary part of $\hat{W}$ and $\hat{\Delta}$ have the same support on $\mathbb{R}_{\geq 0}$. The determinant equals zero leads to a cubic equation in $z$ (after factoring out two trivial roots of $z = 0$) and can be solved in closed-form. For $0 < g < 1$ and $\alpha \geq 0$, this has 3 real roots of $z$, one negative and two positive. The two positive roots correspond to the edges of

$p_{\hat{C}}(x)$. As a function of $\alpha$, the left edge of the support attains its minimum of 0 at $\alpha = 1$ (Fig. S9B). For any fixed $\alpha$, the left edge decreases with $g$ towards the curve of $g \to 1^-$ where

$$x_- = \frac{2}{27}\left((1+3\alpha)^{\frac{3}{2}} + 1 - 9\alpha\right). \tag{S96}$$

The right edge $x_+$ increases with both $\alpha$ and $g$ (Fig. S9B).

We can also use Eq. (S95) to show a power-law approximation for the sample covariance spectrum as $g \to 1^-$. Plugging in $g = 1$ in Eq. (S95), we get the equation for $\hat{W}$ of $\hat{C}$ (here $g^2\hat{C} = \hat{C}$)

$$(z+\alpha)^2\hat{W}^3 + (2z + 2\alpha + \alpha^2)\hat{W}^2 + (1+2\alpha)\hat{W} + 1 = 0. \tag{S97}$$

To determine the tail of the distribution of $p_C(x)$, we consider the roots of the above equation in the limit of $z \to +\infty$. We can determine the order of $\hat{W}$ in $z$ by assuming each term in the equation is the leading one and check for consistency. There are three possibilities corresponding to different terms in Eq. (S97) being dominant: $\hat{W} = O(1), O(z^{-\frac{1}{2}})$, or $o(z^{-\frac{1}{2}})$. The only consistent case is the last one, where the equation approximates to the leading order as

$$z^2\hat{W}^3 + 1 = 0, \quad \text{thus } \hat{W}_i = -\zeta_i z^{-\frac{2}{3}}.$$

Here $\zeta_i^3 = 1$ are the cubic roots of unity. The imaginary parts of $\hat{W}_i/z$ correspond to $p_{\hat{C}}(x)$, therefore,

$$p_{\hat{C}}(x) \approx \frac{\sqrt{3}}{2\pi}x^{-\frac{5}{3}}. \tag{S98}$$

This power-law approximation holds for any fixed $0 \le \alpha \le 1$ as $g \to 1$ (Fig. S9C). Importantly, it is exactly the same approximation as the non-sampled covariance (Eq. (7)). For $\alpha > 1$, the power-law approximation is modified to $p_{\hat{C}}(x) \approx \frac{\alpha\sqrt{3}}{2\pi}x^{-\frac{5}{3}}$ due to re-scaling of non-zero eigenvalues.

#### S9 Space-sampled covariance matrix

Consider the random spatial subsampling where we randomly select a subset of $N_s$ neurons to measure their activity. The spatially subsampled covariance matrix can be written as

$$\tilde{C} = V^T C V, \tag{S99}$$

where $V$ is an $N \times N_s$ matrix and $V_{ij} = \delta_{i,p_j}$, where $p_j, j = 1, \ldots, N_s$ are distinct and randomly chosen from $1, \ldots, N$. As in the time-sampled case (Section S8), the eigenvalues of $\tilde{C}$ are the same as the *non-zero* eigenvalues of

$$CVV^T. \tag{S100}$$

If this is a free product of $C$ and $VV^T$, we can similarly use the tools of free probability to determine the eigenvalue distribution of $\tilde{C}$ as in Section S8. We need to check a sufficient condition for free product [9]: the factors are independent and one of the them is distribution invariant under any orthogonal similarity transformation (referred below as orthogonal invariant for short). The first part of the condition is satisfied and we discuss below three cases that satisfy the second condition.

**Case (i)**: the distribution of $C$ is orthogonal invariant. This is the case for iid Gaussian random connectivity. For any orthogonal matrix $U$

$$U^T C U = (I - U^T J U)^{-1}(I - U^T J U)^{-T}.$$

Since $U^T J U$ has the same distribution as $J$, $U^T C U$ has the same distribution as $C$.

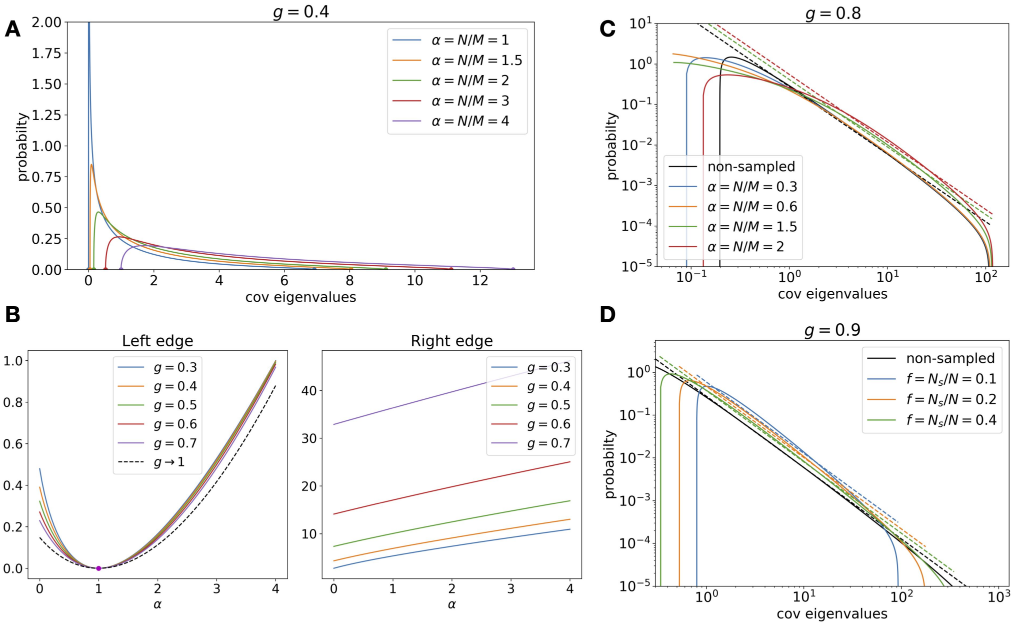

Figure S9: **A**. Same as Fig. 7A. but for $\alpha \geq 1$. **B**. How the left and right edge of the support changes with $\alpha$ in temporal sampling. **C**. Log-log scale plot showing the approximate power-law for time-sampled spectra. The dashed lines (with corresponding color) are power-law approximations (see text). Note that for $\alpha \leq 1$, the power-law approximations are identical (black dashed line). **D**. Same as C. but for space-sampled spectra.

The other possibility for satisfying the condition is $VV^T$ being orthogonal invariant. However, this is not the case for the subsampling described above, where $VV^T = D$ is a diagonal matrix with $N_s$ 1's appear on random locations of the diagonal and everywhere else being 0. After an orthogonal transformation, $D$ can be a dense matrix. Nonetheless, we can modify the subsampling procedure to be *random orthogonal projections* rather than insisting that the directions has to be along one of the coordinates/neurons. This leads to **Case (ii)** where the columns of $V$ are orthonormal vectors in $\mathbb{R}^N$ chosen randomly (for example as the first $N_s$ columns of an orthogonal matrix which is chosen randomly according to the Haar measure). By construction, $VV^T$ is orthogonal invariant. Moreover, it is easy to see that $VV^T$ has the same eigenvalues as $D$. So the results on the eigenvalue distribution of $\tilde{C}$ will be *identical* for Case (i) and (ii).

A common variant of random projection is **Case (iii)** where the entries of $V$ are drawn as iid zero-mean Gaussian variables with variance $1/N$. It is easy to check that $VV^T$ is also orthogonal invariant. For large $N$ and $N_s \ll N$, the columns of $V$ are indeed approximately orthonormal and one expects the same result as Case (ii). However, for the case where $N_s = fN$, $0 < f < 1$ being a constant, the eigenvalue distribution of $VV^T$ is different from $D$. Note the free product Eq. (S100) in Case (iii) is the same as Eq. (S82) in time-sampled case with $\alpha = 1/f$. Hence Case (iii) can be analyzed (but not presented here) using results in Sec.S8 with an additional step of removing $(1 - f)N$ zero eigenvalues (this step is described below for Case (i)/(ii)).

Based on the above discussion, we derive below the results on eigenvalue distribution of the space-sampled covariance for Case (i) and (ii) which has the same resulting spectrum. We conjecture that the spectrum developed under Case (i) also applies to networks with entry distributions other than Gaussian, for example Erdős-Rényi and EI connectivity in Sections 3.5 and S6 , which we have confirmed by numerical simulations (data not shown).

##### S9.1 Dimension

Given the free product $C(VV^T) = CD$, it is straightforward to use the free moment-cumulant relations (see Section S8.1) to show

$$\kappa_n(\tilde{S}) = f^{n-1}\kappa_n(S), \quad n = 1, 2, \ldots. \tag{S101}$$

$$\mu_1(\tilde{C}) = \frac{1}{f}\phi(CD) = \frac{1}{f}\kappa_1(C)\kappa_1(D) = \mu_1(C).$$

$$\begin{aligned}
\mu_2(\tilde{C}^2) &= \frac{1}{f}\phi(CDCD) \\
&= \frac{1}{f}\left(\kappa_2(C)\kappa_1(D)^2 + \kappa_2(D)\kappa_1(C)^2 + \kappa_1(C)^2\kappa_1(D)^2\right), \\
&= \kappa_2(C)f + \kappa_1(C)^2
\end{aligned}$$

We have used $\mu_n(D) = 1/f$, $n \geq 1$. Combining the above equations on first two moments, we have the following relation between the (relative) dimensions

$$\hat{D}(\tilde{C}) = \frac{\hat{D}(C)}{f + (1-f)\hat{D}(C)}, \quad \text{and } D(\tilde{C}) = D(C)\frac{N}{N + \frac{1-f}{f}D(C)}. \tag{S102}$$

Note the relative dimension is with respect to the number of sampled neurons $N_s = fN$, $\hat{D}(\tilde{C}) = D(\tilde{C})/(fN)$. In particular, we always have

$$\hat{D}(\tilde{C}) \geq \hat{D}(C), \quad \text{and } D(\tilde{C}) \leq D(C), \tag{S103}$$

meaning that the relative dimension increases under spatial subsampling while the raw dimension decreases. Interestingly, if we compare Eq. (23) and Eq. (S102), we see that the relation between raw dimension $D$ is the same as in the temporal sampling if we replace $\frac{1-f}{f}$ with $\alpha$.

##### S9.2 Equation relating the sampled and exact spectrum

We can use the similar method of S-transform and free product as in the time-sampled case to calculate the eigenvalue distribution under spatial sampling. First, it is straightforward to calculate the Stieltjes transform of the diagonal matrix $D$

$$\Delta_D(z) = -\frac{1-f}{z} + \frac{f}{1-z}.$$

Using this along with Eq. (S86) we have,

$$S_D(z) = \frac{1+z}{f+z}.$$

This replace the role of $S_{MP}(z)$ in the derivation. We also need to remove the trivial zero eigenvalues in the $\Delta_{CD}(z)$ in the second last step of Section S8.2 before inverting the Stieltjes transform.

This can be done similarly as Eq. (S93). The rest of the derivation follows the same steps as in Section S8.2 and we get

$$f\tilde{W}\left(\frac{z(f+W(z))}{1+W(z)}\right)=W(z),\quad \text{and conversely } f\tilde{W}(z)=W\left(\frac{z(1+f\tilde{W}(z))}{f(1+\tilde{W}(z))}\right). \tag{S104}$$

We can also describe the combined effect of temporal and spatial sampling on the spectrum. Let $W_1(z)$ be the generating function the spectrum time-sampled with $\alpha$ and then space-sampled with $f$. For simplicity, assume $\alpha f \leq 1$ so that we don't have zero eigenvalues in the end. Then by combining Eqs. (22) and (S104) we have,

$$fW_1\left(\frac{z(1+\alpha W(z))(f+W(z))}{1+W(z)}\right)=W(z),\quad fW_1(z)=W\left(\frac{z(1+fW_1(z))}{f(1+\alpha f W_1(z))(1+W_1(z))}\right). \tag{S105}$$

One can also apply the spatial sampling first then temporal sampling and get the same result as Eq. (S105). However, in this case the sampling parameters should be $f$ and $\alpha f$ since the number of neurons changed after spatial sampling.

##### S9.3 Space-sampled covariance spectrum for iid Gaussian random connectivity

Here we apply the general result of spatial subsampling to the random iid Gaussian connectivity. Similar to the time-sampled case, we start from Eq. (S94) and plug in Eq. (S104). However, unlike the time-sampled case, the resulting equation for $\hat{W}(z)$ is a fifth-order polynomial and does not appear to be analytically tractable even for calculating the edges of the support. We can nonetheless solve the equation numerically. To select the correct pairs among potentially multiple complex roots that correspond to $p_{\tilde{C}}(x) =: p_{g,f}(x)$, we can use the continuity when varying $f$ from 1, which is the original spectrum $p_C(x)$. We found that for $0 < g, f < 1$, the equation of $\tilde{W}$ only has one pair of complex roots for $z = x > 0$, which gives $p_C(x)$. We have verified our choice of branch/roots by comparing the numerical solution with simulations of random connectivity (data not shown).

We observe that for any $f > 0$, the pdf of the space-sampled distribution narrows inwards comparing to the non-sampled case (Fig. 7). We therefore numerically solved for the support edges of $\tilde{W}(z)$ using a bisection search inside the known support of the non-sampled case (Eq. (6)) for when $\mathrm{Im}(\tilde{W}(z))$ first becomes to 0.

To determine the power-law tail approximation, we let $g \to 1^-$ in the equation of $\tilde{W}(z)$ which gives

$$\begin{aligned}&-f^3z^2\tilde{W}^5-2f^2z(-1+f(1+r)z)\tilde{W}^4+(f+2f^2(2+r)z-f^3(1+r)^2z^2)\tilde{W}^3\\&+(1+2f+2f^2(1+r)z)\tilde{W}^2+(2+f)\tilde{W}+1=0.\end{aligned}$$

Here $r = (1-f)/f$. By assuming whether each term in the above equation has an $O(1)$ contribution (i.e., the leading term in the equation) as $z \to \infty$, we have the following possible orders of $\tilde{W}$: $O(1)$, $O(z^{-1/2})$, $O(z^{-2/3})$, $O(z^{-1/2})$, $O(z^{-2/5})$, or $o(z^{-2/3})$. The only consistent one is $\tilde{W} \sim O(z^{-2/3})$, and to the leading order the equation is approximated as

$$f^3(1+r)^2z^2\tilde{W}^3=1,\quad \Rightarrow \quad \tilde{W}=\zeta_i z^{-2/3}f^{-1}(1+r)^{-2/3}. \tag{S106}$$

Here $\zeta_i$ are cubic roots of unity. The pair of complex roots that correspond to a probability density function gives (Fig. S9D)

$$\mathrm{Im}\tilde{W}=-\frac{\sqrt{3}}{2}f^{-\frac{1}{3}}z^{-\frac{2}{3}},\quad p_{\tilde{C}}(x)\approx\frac{\sqrt{3}}{2\pi}f^{-\frac{1}{3}}x^{-\frac{5}{3}}. \tag{S107}$$

We see that this has the same power-law exponent as the original covariance spectrum Eq. (7).

Motivated by the results shown for the random iid Gaussian connectivity (Eqs. (S98) and (S107)), we conjecture that a power-law tail in the critical spectrum ($g \to 1$) will persist with the same exponent after temporal and spatial sampling. Specifically, if the original distribution satisfies $\beta > 1$ (for integrability of pdf $p(x)$),

$$\lim_{x\to\infty} p(x)x^{\beta} = C_0,$$

then the spectra of the temporal and space-sampled covariance for any fixed $0 \leq \alpha \leq 1$, $0 < f \leq 1$ satisfy

$$\lim_{x\to\infty} p_{\hat{C}}(x)x^{\beta} = C_0, \quad \lim_{x\to\infty} p_{\tilde{C}}(x)x^{\beta} = C_0 f^{\beta-2}. \tag{S108}$$

#### S10 The spectrum of the correlation matrix

Here we explain our heuristic argument of how the results for the spectrum of the covariance matrix in the large $N$ limit can be easily translated to that of the correlation matrix for the random connectivity models we considered.

The important observation is that in the large $N$ limit, the diagonal entries of the covariance $C$ converge to the same (positive) value $\mu = \lim_{N\to\infty} \mathrm{tr}(C)/N$, which is the mean/first moment of the eigenvalues. More precisely,

$$\lim_{N\to\infty} \frac{1}{N} \sum_i (C_{ii} - \frac{1}{N}\mathrm{tr}(C))^2 = 0. \tag{S109}$$

This uniform limiting diagonal motivates that the spectrum of the correlation matrix $R$ is

$$p_R(x) = \mu p_C(\mu x). \tag{S110}$$

For the Gaussian random connectivity model, iid or with reciprocal motifs, we verified Eq. (S109) numerically (Fig. S10). Note that the mean $\mu$ for these models are also available analytically (Eqs. (9) and (18), which makes it easy to apply the rescaling.

#### S11 Additional results on fitting the theoretical spectrum to data

Here we describe the details on fitting to simulated covariance eigenvalues under low rank activity perturbation (Section 3.8). We add a rank-2 matrix to the covariance matrix generated from iid Gaussian connectivity (Eq. (2))

$$C = \sigma_1^2 u_1 u_1^T + \sigma_2^2 u_2 u_2^T + (I - J)^{-1}(I - J)^{-1} \tag{S111}$$

Here we set $\sigma^2 = 1$ without loss of generality. Column vectors $u_1$ and $u_2$ are chosen as random vectors with unit 2-norm. $\sigma_1$ and $\sigma_2$ are set to be sufficiently large to create two large outlying eigenvalues which nonetheless may not obviously stand out in PCA due to the long tail of the bulk (see inset of Fig. S11). The location of these outliers can be predicted analytically for large $N$ (Section S3.2). But this knowledge and the value of $\sigma_i^2$ are not needed in the fitting due to the robustness of the bulk (Section S3).

We fit the theoretical spectrum Eq. (5) to all the eigenvalues including the outliers (unknown to the fitting algorithm). The inferred $g$ is highly accurate despite the presence of outliers for a moderate size network ($N = 200$): the root mean square error (rMSE) in $g = 0.6$ is 0.0321, or 5.35%. Here we fit a theoretical spectrum to each realization of Eq. (S111) and record the error

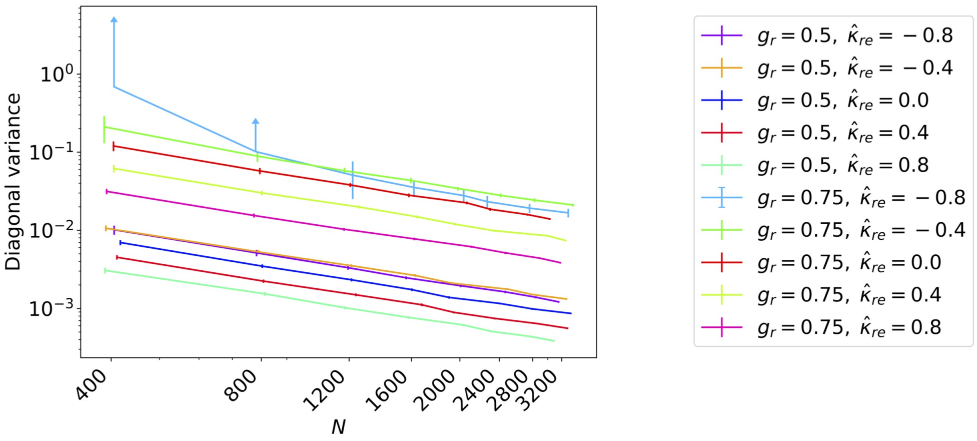

Figure S10: The variance of diagonal entries (Eq. (S109)) for various combinations of $g$ and reciprocal motif cumulants $\hat{\kappa}_{re} = \rho(J_{ij}, J_{ji})$ ($\hat{\kappa}_{re} = 0$ is the iid random connectivity). The lines and the error bars shows the average and standard deviation across 100 trials. $N = 400, 800, 1200, 1600, 2000, 2400, 2800, 3200$. For $g_r = g(1 + \hat{\kappa}_{re}) = 0.75$ and $\hat{\kappa}_{re} = -0.8$, the average is smaller than the standard deviation, thus only the upper error bar (with an arrow) is shown in this log-log scale plot. A small jitter in the x-axis is added to each line for clearer visualization.

$\hat{g} - g$, and then average these errors over 100 realizations to get the above rMSE. We can use the fitted $\hat{g}$ to calculate the upper edge of the support (Eq. (6)) and use it as a cut-off to separate the outlying eigenvalues. We can then re-fit the spectrum leaving out the putative outliers, and iterate the process a few times or until convergence. This process further improves the fitting accuracy (rMSE of $g$ is 0.01) and correctly identifies the two outliers (Fig. S11) in 86 of the 100 realizations of Eq. (S111).

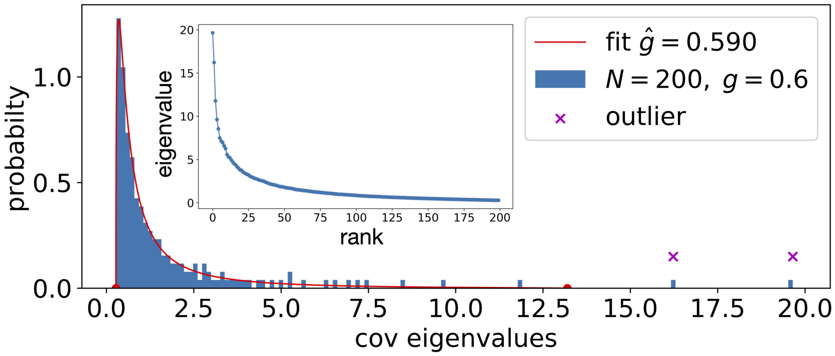

Figure S11: Fitting the theoretical spectrum (red curve) to the eigenvalues of a covariance matrix with iid Gaussian connectivity perturbed by a rank-2 matrix (see text). Shown is one realization with $N = 200$, $g = 0.6$, $\sigma_1^2 = 17$, $\sigma_2^2 = 15$. The theoretical support at the fitted $g$ correctly separates out the outliers (highlighted by magenta crosses). The inset plots the eigenvalues vs. rank, where there is no obvious visual separation between the bulk and outliers.

Figure S12 shows the fitting results of additional clusters in the whole-brain calcium imaging data in larval zebrafish (Fig. 8). The details, including the numbering of the clusters, are explained in the main text and Fig. 8.

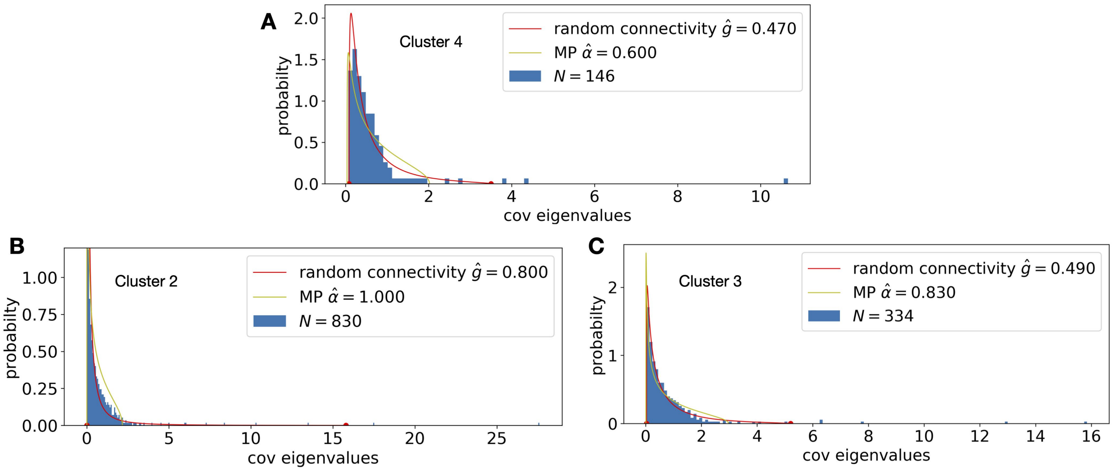

Figure S12: Same as Fig. 8D,E. but for the rest of functional clusters in Fig. 8A.

### S12 Ordered and deterministic connectivity

For the examples of deterministic connectivity considered in the main text, their eigenvectors are given by the discrete Fourier basis and are therefore orthogonal to each other. This means that the connectivity $J$ is a normal matrix and the eigenvalues of the covariance matrix can be directly calculated from those of $J$.

#### S12.1 Long-range ring network

Here $J$ is given by Eq. (35),

$$J_{ij} = \frac{1}{N} f(x_i - x_j) = \frac{1}{N} f\left(\frac{i-j}{N}\right), \quad x_i = i/N,\ i = 0, \dots, N-1$$

where $f(x)$ is a smooth function. The all-to-all uniform network is a special case with $J_{ij} = g/N$ ($f(x) \equiv g$). It is convenient to allow self-coupling so that $J_{ii} = g/N$. $J$ is a rank-1 matrix, with one eigenvalue being $g$ and $N-1$ eigenvalues being 0 (without self-coupling these are $g\frac{N-1}{N}$ and $-\frac{g}{N}$) and

$$p_C(x) = \frac{1}{N}\delta\left(x - \frac{1}{(1-g)^2}\right) + \frac{N-1}{N}\delta\left(x-1\right). \tag{S112}$$

In the large $N$ limit, $p_C(x)$ converges to be a delta function at 1 and an outlier at $(1-g)^{-2}$.

For a general long-range ring network (again, for simplicity we allow self-connection but this would have negligible effect in the large $N$ limit), the eigenvalues of $J$ in the large $N$ limit correspond to the Fourier coefficients of $f(x)$. For each fixed $k$,

$$\lambda_k = \frac{1}{N}\sum_{i=0}^{N-1} e^{-\frac{2\pi \mathbb{i} k}{N} i} f(x_i) \to \int_0^1 e^{-2\pi \mathbb{i} k x} f(x) dx =: \lambda_k^\infty.$$

For a smooth ($C^\infty$) $f(x)$ , $\lambda_k^\infty \to 0$ (faster than any polynomial of $k$) , as $k \to \infty$. This indicates that most eigenvalues of $J$ are near 0, except for "a few" eigenvalues that approach their fixed locations $\lambda_k^\infty$ as $N \to \infty$. More precisely, for any $\delta > 0$, there exists $N_0$, such that for all $N > N_0$, all but $M(\delta)$ eigenvalues of $J$ lie outside of circle $|z| < \delta$ on the complex plane. This means that the eigenvalue distribution of $J$ in the large $N$ limit is a delta function at 0.

The property of eigenvalues of $J$ translates to that of $C$ as a delta function at 1, except for a number of discrete eigenvalues near $|1 - \lambda_k^\infty|^{-2}$ (Fig. 9A). We note that such a "sparse tail" of discretely located eigenvalues is very different from the continuous density function in the case of random networks. The above results also hold for $d$-dimension rings (i.e., a torus $S^1 \times \cdots \times S^1$) that have long-range connectivity. The eigenvectors of the torus is the tensor products of the 1D discrete Fourier basis.

#### S12.2 Nearest-Neighbor ring

Here neuron $i$ is connected to the two neighboring neurons $i \pm 1$ and the connectivity has two parameters: $x = J_{i-1,i}$ and $y = J_{i+1,i}$. By Fourier transform the eigenvalues of $J$ are

$$\lambda_k = a\cos(\theta_k) + \mathbb{i}b\sin(\theta_k), \quad a = x + y, \; b = x - y, \quad \theta_k = \frac{2\pi k}{N}, \; k = 0, \ldots, N-1.$$

Since $\theta_k$ are evenly spaced over $[0, 2\pi)$, we can re-define

$$a = |x + y| \geq 0, \;\; b = |x - y| \geq 0.$$

without changing the set of eigenvalues $\{\lambda_k\}$. The stability of the network dynamics requires $a = |x + y| < 1$.

The eigenvalues of $C$ are

$$c_k = |1 - \lambda_k|^{-2} = ((1 - a\cos(\theta_k))^2 + b^2\sin^2(\theta_k))^{-1}. \tag{S113}$$

Since

$$(1 - a\cos(\theta_k))^2 + b^2\sin^2(\theta_k) \geq (1 - a)^2 + 0,$$

which attains equality at $k = 0$, the maximum of $c_k$ is $(1 - a)^{-2}$, which is the upper edge of the covariance spectrum support.

For the minimum of $c_k$, a natural candidate is $\theta_k = \pi$, and $c_k = (1 + a)^{-2}$ (we neglect the small difference due to an odd $N$ since $N$ is large and write $\theta_k = \theta$ below to signify the limit where $\theta$ is considered continuous). However, $c(\theta = \pi)$ is the global minimum if and only if

$$b^2 < a^2 + a. \tag{S114}$$

Otherwise, the global minimum is attained at

$$\theta = \pm\arccos(-a/(b^2 - a^2)), \quad c_{\min} = \frac{b^2 - a^2}{b^2(b^2 - a^2 + 1)}. \tag{S115}$$

Here we have two solutions for $\theta$ due to symmetry. We proceed to derive the pdf $p_C(x)$ for each of these two cases respectively.

**Regular case** When Eq.(S114) is satisfied, we call this the *regular case.* First, consider the precision matrix eigenvalues

$$p = c^{-1} = (1 - a\cos(\theta))^2 + b^2\sin^2(\theta) = 1 + b^2 + (a^2 - b^2)\cos^2(\theta) - 2a\cos(\theta). \tag{S116}$$

The density of $p$ can be expressed using its mapping with $\theta$ (only consider $\theta \in [0, \pi)$ due to symmetry) and the density of $c$ follows,

$$p_P(x) = \frac{1}{\pi}\left|\frac{dp}{d\theta}\right|^{-1}, \quad p_C(x) = \frac{1}{x^2}p_P\left(\frac{1}{x}\right). \tag{S117}$$

The density of $c$ also follows easily from that of its reciprocal $p$. Solving $\cos(\theta)$ in Eq.(S116)

$$\cos(\theta) = \frac{a \pm \sqrt{D}}{a^2 - b^2}, \quad D = b^2 - (b^2 - p)(a^2 - b^2). \tag{S118}$$

Due to Eq. (S114), only the root with "−" sign corresponds to $\cos(\theta)$. To proceed,

$$\frac{dp}{d\theta} = 2\sin(\theta)(-(a^2 - b^2)\cos(\theta) + a).$$

Plugging Eq. (S118) to the above and use Eq. (S117), we have the density for the covariance

$$p_C(x) = \frac{|a^2 - b^2|}{2\pi x^2\sqrt{D(a^2 - b^2 - a + \sqrt{D})(a^2 - b^2 + a - \sqrt{D})}}, \tag{S119}$$

where

$$x \in \left[(1+a)^{-2}, (1-a)^{-2}\right], \quad D = b^2(1 - a^2 + b^2) + \frac{a^2 - b^2}{x}.$$

**Folded case** When Eq. (S114) is not satisfied, i.e., $b^2 > a^2 + a$, we call this the *folded case.* We only need to consider $\theta \in [0, \pi)$ due to symmetry. Let $\theta_0$ be the angle where the maximum of $p$ is attained (given in Eq. (S115)). The relation between $\theta$ and $p$ is monotonic and thus one-to-one on each segment $[0, \theta_0]$ (increasing) and $[\theta_0, \pi]$ (decreasing) respectively, and hence we will consider them separately.

Note that both of the two roots of $\cos(\theta)$ in Eq. (S118) are meaningful if $p \geq p(\pi)$. Moreover, since $b^2 > a^2$, the root with the "−" sign corresponds to a $\theta \in [0, \theta_0]$ and the root with the "+" sign corresponds to a $\theta \in [\theta_0, \pi)$. For $p < p(\pi)$, only the negative sign root is meaningful (and $\theta \in [0, \theta_0]$).

Based on the above analysis and select the correct root $\cos(\theta)$ in Eq. (S118), we derive the density of $p$ and then $c$ in the same way as the regular case on each $\theta$ segment. For $\theta \in [0, \theta_0]$, the density function for $c_k$ is

$$p_1(x) = \frac{b^2 - a^2}{2\pi x^2\sqrt{D(a^2 - b^2 - a + \sqrt{D})(a^2 - b^2 + a - \sqrt{D})}},$$

where $D$ is the same as in Eq.(S119). For $\theta \in [\theta_0, \pi]$,

$$p_2(x) = \frac{b^2 - a^2}{2\pi x^2\sqrt{D(a^2 - b^2 - a - \sqrt{D})(a^2 - b^2 + a + \sqrt{D})}}.$$

Combining together,

$$p_C(x) = \begin{cases} p_1(x) + p_2(x), & x \in [c_{\min}, (1+a)^{-2}) \\ p_1(x), & x \in [(1+a)^{-2}, (1-a)^{-2}] \end{cases} , \tag{S120}$$

where $c_{\min}$ is given by Eq. (S115).

Using the analytic expressions Eqs. (S119) and (S120), we now summarize the shape of the covariance spectra $p_C(x)$ in the two cases of NN ring. In the regular case (Fig. 9B), $p_C(x)$ diverges at the two edges as $(\Delta x)^{-\frac{1}{2}}$. In the folded case (Fig. S13), the same diverging densities (due to $p_1(x)$ term) now appear at the upper edge of $p_C(x)$ and $c_{\min} + 0^-$, which is inside the support. In addition, at the left edge corresponding to $\sqrt{D} = 0$, $p_C(x)$ diverges as $(\Delta x)^{-\frac{1}{2}}$. The situation of having singularity in the middle of the distribution is qualitatively similar to that in the anti-symmetric random network (at low frequency $\omega$, Eq. (S57)). Because the density is more complex in the folded case with the additional singularity, we focus on the regular case to further explore comparisons with the random network.

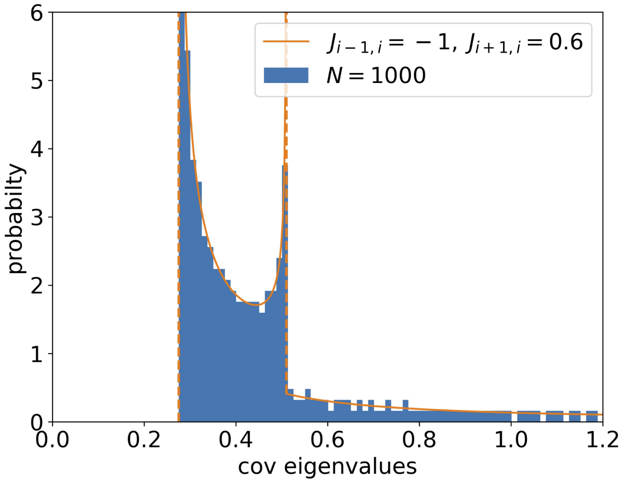

Figure S13: Th covariance spectrum a Nearest-Neighbor ring under the folded case: $J_{i-1,i} = -1$, $J_{i+1,i} = 0.6$. The theory (Eq. (S120)) matches well with the finite-size simulation at $N = 1000$. For visualizing the interior singularity at $(1+a)^{-2}$ (see text), the full tail of the distribution is cropped in the plot.

##### S12.3 Multi-dimensional Nearest-Neighbor ring

One outstanding difference between the $p_C(x)$ for the Nearest-Neighbor (NN) ring network and the random networks is the diverging density at the edges of the support. This divergence is because the eigenvalues of $J$ are distributed along a 1D curve rather than in a 2D region of the complex plane. We thus consider the generalization of the NN ring network to higher dimensions and in particular examine how the edge density may change.

In a $d$-dimension NN ring, the neurons are located on an equally spaced lattice of $T^d = S^1 \times S^1 \times \cdots S^1$. Each neuron is connected to $2d$ nearest neighbors with connection strength along the $k$-th coordinate axis being $J^k_{i_k-1,i_k} = x_k$ and $J^k_{i_k+1,i_k} = y_k$. Since the tensor products of the 1D discrete Fourier basis are eigenvectors, the eigenvalues of $J$ are

$$\lambda_{\vec{k}} = \sum_i a_i \cos(\theta_{k_i}) + \mathbb{i} \sum_i b_i \sin(\theta_{k_i}), \quad a_i = |x_i + y_i| \geq 0, \; b_i = |x_i - y_i| \geq 0.$$

The stability requires that $\sum_i a_i < 1$. Based on the results of the 1D ring, we will focus on an analogue of the regular case, that is, the global minimum and maximum of

$$p_{\vec{k}} = p(\vec{\theta}_k) = |1 - \lambda_{\vec{k}}|^2, \quad p(\vec{\theta}) = \sum_{i=1}^{d} (1 - a_i \cos(\theta_i))^2 + \sum_{i=1}^{d} b_i^2 \sin(\theta_i) \tag{S121}$$

should be obtained at the natural candidates

$$\vec{\theta}_1 = (0, \ldots, 0), \text{ and } \vec{\theta}_2 = (\pi, \ldots, \pi).$$

Here $\vec{k} = (k_1, k_2, \ldots, k_d)^T$, $0 \leq k_i \leq N - 1$ is the frequency vector. First, it is easy to see that $\vec{\theta}_1$ is the minimum of $p$, since

$$p \geq (1 - \sum_i a_i)^2 + 0 = p(\vec{\theta}_1).$$

Similarly as the 1D case, we need additional conditions on $a_i$, $b_i$ to ensure that the maximum of $p$ is obtained at $\vec{\theta}_2$ (thus the regular case). The following lemma gives a sufficient condition. For 1D, the condition is equivalent to Eq. (S114).

**Lemma S12.1.** *Under the stability condition $\sum_{i=1}^{d} a_i < 1$ and the following inequality*

$$\sum_i \left| \frac{b_i^2}{a_i} - a_i \right| < 1, \quad \text{and } a_i > 0, i = 1, \ldots, d, \tag{S122}$$

*the global maximum of $p(\vec{\theta})$ (minimum of $c(\vec{\theta})$) is obtained at $\vec{\theta}_2 = (\pi, \ldots, \pi)$.*

*Proof.* Consider a stationary point of $p(\vec{\theta})$, the derivative along each $\theta_i$ should be 0,

$$0 = \frac{1}{2}\frac{\partial p}{\partial \theta_i} = (1 - \sum_j a_j \cos(\theta_j)) a_i \sin(\theta_i) + (\sum_j b_j \sin(\theta_j)) b_i \cos(\theta_i).$$

$$- a_i \sin(\theta_i) = \mu b_i \cos(\theta_i), \quad \mu = \frac{\sum_j b_j \sin(\theta_j)}{1 - \sum_j a_j \cos(\theta_j)}, \quad i = 1, \ldots, d. \tag{S123}$$

If $\mu = 0$, then all $\theta_i = 0$, so that all $\cos(\theta_i) = \pm 1$. It is easy to see among these points, the maximum of $p$ is obtained at $\vec{\theta}_2$.

If $\mu \neq 0$, then $\sum_j b_j \sin(\theta_j) \neq 0$. Multiplying Eq. (S123) with $b_i/a_i$ and sum across $i$,

$$-\sum_i b_i \sin(\theta_i) = \mu \sum_i \frac{b_i^2}{a_i} \cos(\theta_i).$$

Now the left-hand side is the numerator of $\mu$. Divide the above by $\mu$ and move the $\cos(\cdot)$ terms to the right,

$$-1 = \sum_i (\frac{b_i^2}{a_i} - a_i) \cos(\theta_i).$$

This, however, with Eq. (S122),

$$1 = \left| \sum_i (\frac{b_i^2}{a_i} - a_i) \cos(\theta_i) \right| \leq \sum_i \left| \frac{b_i^2}{a_i} - a_i \right| < 1,$$

leads to a contradiction. Therefore, there are no stationary points with $\mu = 0$, and we conclude $p(\vec{\theta}_2)$ is the global minimum. □

We will from now on assume condition (S122). This means the support of $c(\vec{\theta})$ is

$$c(\vec{\theta}) \in \left[(1+\sum_i a_i)^{-2}, (1-\sum_i a_i)^{-2}\right].$$

To analyze the density at the upper edge of $c$, consider the Taylor expansion of $p$ around $\vec{\theta}_1$ and take the leading-order terms,

$$p \approx (1-\sum_i a_i)^2 + (1-\sum_i a_i)\sum_i a_i\theta_i^2 + (\sum_i b_i\theta_i)^2. \tag{S124}$$

For a small $\delta > 0$, the region in $\vec{\theta}$ and satisfies $p(\vec{\theta}) - p(\vec{\theta}_1) < \delta$ is approximately an ellipsoid (because the second-order terms above are positive definite)

$$(1-\sum_i a_i)\sum_i a_i\theta_i^2 + (\sum_i b_i\theta_i)^2 \leq \delta.$$

Its volume is $V(\delta) = C_0\delta^{\frac{d}{2}}$, where $C_0$ is a constant that depends on $a_i, b_i$. Since the density for $\theta_i$ is uniform, the leading-order term is

$$P(p(\vec{\theta}) - p(\vec{\theta}_1) < \delta) \approx \frac{1}{(2\pi)^d}V(\delta) = \frac{C_0}{(2\pi)^d}\delta^{\frac{d}{2}}. \tag{S125}$$

Hence the density near the upper edge of $p_C(x)$ is

$$p_C(x) \approx C_1(\Delta x)^{\frac{d}{2}-1}. \tag{S126}$$

The analysis around the lower edge $\vec{\theta} = \vec{\theta}_2$ is similar (expand in terms of $\hat{\theta}_i = \pi - \theta_i$), and we obtain the same result as Eq. (S126).

##### S12.3.1 Power-law tail approximations

Here we study the approximate power-law tail of $p_C(x)$ as the connection strength approaches the critical value $\sum_i a_i \to 1^-$. In this limit, the lower support of the distribution of $p$ goes to 0, which corresponds to the upper edge of $p_C(x)$ diverging to infinity. The leading-order approximation of the probability of $p$ near the 0 edge similar to Eq. (S125) will translate to a power law of $p_C(x)$ for $x \gg 1$.

First consider the 1D ring and start with the approximation Eq. (S116), but expand near 0 to a further order since here $a = 1$,

$$p \approx \frac{\theta^4}{4} + b^2\theta^2 - b^2\frac{\theta^4}{3}.$$

If $b \neq 0$, then the leading term is $b^2\theta^2$, and an analysis of the density for $p \in [0, \delta]$ similar as in Eq. (S125) shows that $p(p) \propto (\Delta x)^{-\frac{1}{2}}$. This in turn means

$$p_C(x) \approx \frac{1}{2\pi|b|}x^{-\frac{3}{2}}. \tag{S127}$$

If $b = 0$, then the leading term is $\theta^4/4$, and correspondingly $p(p) \propto (\Delta x)^{-\frac{3}{4}}$ and

$$p_C(x) \approx \frac{\sqrt{2}}{4\pi}x^{-\frac{5}{4}}. \tag{S128}$$

We can also directly verify these tail approximations using the explicit formulas of $p_C(x)$ (Eqs. (S119) and (S120)).

For a multi-dimensional ring (in the regular case assuming condition Eq. (S122)), there could be various situations for the leading term approximation for $p(\vec{\theta})$ in Eq. (S121) depending on, for example, how many $b_i$ are 0. For simplicity, we consider two cases that correspond to the two cases we see in the 1D ring (Eq. (S127), (S128))).

The first case is when all $b_i \neq 0$. An expansion of $p$ near 0 shows its leading terms are

$$p \approx \left( \sum_{i=1}^{d} b_i \theta_i \right)^2 .$$

The probability for $p \in [0, \delta)$ is proportional to the volume of the region of a scaled $L^1$-ball [13] with a half-axis being $\sqrt{\delta}/|b_i|$ (the $(2\pi)$ factor comes from the density of the angles $\theta_i$). Therefore,

$$P(0 \leq p \leq \delta) \approx (2\pi)^{-d} \frac{2^d}{d!} \delta^{\frac{d}{2}} (\Pi_i |b_i|)^{-1}.$$

In turn the power-law tail approximation of $p_C(x)$ is

$$p_C(x) \approx \frac{1}{2(\pi)^d (d-1)! \Pi_i |b_i|} x^{-\frac{d}{2}-1}. \tag{S129}$$

Another case is when all $b_i = 0$, this also includes the symmetric NN ring we considered in Fig. 9. The leading terms of $p$ are

$$p \approx \left( \sum_{i=1}^{d} \frac{a_i}{2} \theta_i^2 \right)^2 .$$

Here the related volume is an ellipsoid,

$$P(0 \leq p \leq \delta) \approx (2\pi)^{-d} \frac{\pi^{d/2}}{\Gamma(d/2+1)} 2^{d/2} \delta^{\frac{d}{4}} (\Pi_i a_i)^{-1/2}.$$

The power-law tail approximation in this case is

$$p_C(x) \approx \frac{d}{4(2\pi)^{\frac{d}{2}} \Gamma(\frac{d}{2}+1)(\Pi_i a_i)^{\frac{1}{2}}} x^{-\frac{d}{4}-1}. \tag{S130}$$

Note that Eq. (S130) can be directly verified for $d = 1, 2$ using the explicit formulas of $p_C(x)$ (Eqs. (S119) and (S132))

###### S12.3.2 Analytical expression of the covariance spectrum

To further characterize the shape of the covariance spectrum quantitatively in multi-dimensional ring networks, we derive explicit expressions for $p_C(x)$ in the special case of homogeneous symmetric connections: $a_i = a$, $b_i = 0$, $i = 1, \ldots, d$ (i.e., $x_i = y_i = a/2$, corresponding to Fig. 9C-F). The stability condition here becomes $ad < 1$. First consider the density of $x = \sum_{i=1}^{d} \cos(\theta_i)$, where each $\theta_i$ is uniformly distributed on $[0, 2\pi)$. We can express the pdf $p(x)$ using the delta function replacement (equivalent to using the characteristic function)

$$\begin{aligned}
p(x) &= \int_{-\infty}^{\infty} \delta\left(x - \sum_{i=1}^{d} \cos(\theta_i)\right) \prod_i \frac{d\theta_i}{2\pi} d\omega \\
&= \frac{1}{2\pi} \int_{-\infty}^{\infty} e^{\mathbb{i}\omega(x - \sum_{i=1}^{d} \cos(\theta_i))} \prod_i \frac{d\theta_i}{2\pi} d\omega \\
&= \frac{1}{2\pi} \int_{-\infty}^{\infty} d\omega e^{\mathbb{i}\omega x} \left(\int_0^{2\pi} e^{-\mathbb{i}\omega\cos(\theta)} \frac{d\theta}{2\pi}\right)^d \\
&= \frac{1}{2\pi} \int_{-\infty}^{\infty} d\omega e^{\mathbb{i}\omega x} J_0(\omega)^d.
\end{aligned}$$

Here $J_0(\cdot)$ is the Bessel function and we have used its integral definition. Now $p_C(x)$ and $p(x)$ above is related through a change of variable $c = (1 - ax)^{-2}$ or $x = (1 - c^{-1/2})/a$, hence the covariance spectrum for a $d$-dimensional symmetric NN ring is

$$p_{C,d}(x) = \frac{1}{4a\pi} x^{-\frac{3}{2}} \int_{-\infty}^{\infty} \cos(\omega(1 - 1/\sqrt{x})/a) J_0(\omega)^d d\omega. \tag{S131}$$

Note that this is a regular case as Eq.(S119) is satisfied, so the support of $p_{C,d}(x)$ is $[(1+ad)^{-2}, (1-ad)^{-2}]$.

For the 1D ring, Eq. (S131) can be integrated to a closed-form and the result is identical to Eq. (S119) with $b = 0$. For the 2D ring, the integral can be expressed using a special function $K(k)$, the complete elliptic integral of the first kind,

$$p_C(x) = \frac{x^{-\frac{3}{2}} K(1 - \frac{(1-1/\sqrt{x})^2}{4a^2})}{2a\pi^2}, \quad K(k) = \int_0^{\frac{\pi}{2}} \frac{d\theta}{\sqrt{1 - k^2 \sin(\theta)^2}} \tag{S132}$$

For $d > 2$, we can numerically integrate Eq. (S131). Importantly, as a 1D integral it is much more efficient than a naive $d$-dimensional integral.

### References

[1] Z. D. Bai. "Circular law". *Annals of Probability* 25.1 (1997), pp. 494–529.

[2] Z. Burda, A. Görlich, A. Jarosz, and J. Jurkiewicz. "Signal and noise in correlation matrix". *Physica A: Statistical Mechanics and its Applications* 343.1-4 (2004), pp. 295–310.

[3] D. Dahmen, S. Grün, M. Diesmann, and M. Helias. "Second type of criticality in the brain uncovers rich multiple-neuron dynamics". *Proceedings of the National Academy of Sciences of the United States of America* 116.26 (2019), pp. 13051–13060.

[4] F. Götze and A. Tikhomirov. "The circular law for random matrices". *Annals of Probability* 38.4 (2010), pp. 1444–1491.

[5] R. A. Horn and C. R. Johnson. *Matrix Analysis.* Cambridge University Press, 1990.

[6] Y. Hu, S. L. Brunton, N. Cain, S. Mihalas, J. N. Kutz, and E. Shea-brown. "Feedback through graph motifs relates structure and function in complex networks". *Physical Review E* 062312 (2018), pp. 1–25.

[7] Y. Hu, J. Trousdale, K. Josić, and E. Shea-Brown. "Motif statistics and spike correlations in neuronal networks". *Journal of Statistical Mechanics: Theory and Experiment* 2013.03 (2013), P03012.

[8] J. Kadmon and H. Sompolinsky. "Transition to chaos in random neuronal networks". *Physical Review X* 5.4 (2015), pp. 1–28.

[9] J. A. Mingo and R. Speicher. *Free Probability and Random Matrices.* New York, NY, 2017.

[10] H. J. Sommers, a. Crisanti, H. Sompolinsky, and Y. Stein. "Spectrum of large random asymmetric matrices". *Physical Review Letters* 60.19 (1988), pp. 1895–1898.

[11] T. Tao. "Outliers in the spectrum of iid matrices with bounded rank perturbations". *Probability Theory and Related Fields* 155.1-2 (2013), pp. 231–263.

[12] D. Voiculescu. "Multiplciation of certain noncommuting random variables". *Journal of Operator Theory* 18 (1987), pp. 2223–2235.

[13] X. Wang. "Volumes of Generalized Unit Balls Published by : Mathematical Association of America Linked references are available on JSTOR for this article : Volumes of Generalized Unit Balls". *Mathematics Magazine* 78.5 (2005), pp. 390–395.

[14] L. Zhao, B. Beverlin, T. Netoff, and D. Q. Nykamp. "Synchronization from Second Order Network Connectivity Statistics". *Front Comput Neurosci* 5 (Jan. 2011), pp. 1–16.
